# Structure-Dependent Effects of Lipid Conjugation on Cytosolic Accumulation in *Escherichia coli* and Mammalian Cells

**DOI:** 10.64898/2026.06.10.731343

**Authors:** Dylan J. Coffin, Sobika Bhandari, Liora E. Wittle, Karl L. Ocius, George M. Ongwae, Marcos M. Pires

## Abstract

Lipidation is a widely used strategy to promote membrane permeation, yet whether the factors governing lipid-driven accumulation are shared across mammalian and Gram-negative envelopes remains unresolved. Here, we apply the Chloroalkane Azide-based Membrane Penetration (CHAMP) assay to a systematically designed library of lipid conjugates in both HeLa and ***E. coli*** cells. CHAMP, developed by our group, pairs a minimally disruptive azide tag with a cytosolically anchored HaloTag to quantify cytosolic accumulation directly. The two systems show divergent trends: most lipid modifications reduce ***E. coli*** cytosolic accumulation, whereas larger, more hydrophobic conjugates, including medium-chain, cyclized, and heteroatom-containing lipids, are preferentially internalized by mammalian cells. By systematically editing the compound scaffold and, separately, perturbing individual layers of the bacterial envelope, we resolve how charge, scaffold composition, and specific envelope barriers shape these patterns. Together, these results indicate that lipidation is a context-dependent permeation strategy rather than a general one. By showing that hydrophobic modifications frequently hinder cytosolic entry into ***E. coli***, this work offers a permeability-based hypothesis for the scarcity of lipidated antibacterials with cytosolic targets, and delineates where lipophilicity-driven optimization is likely to fail in the diderm envelope.

**TABLE OF CONTENTS FIGURE:** 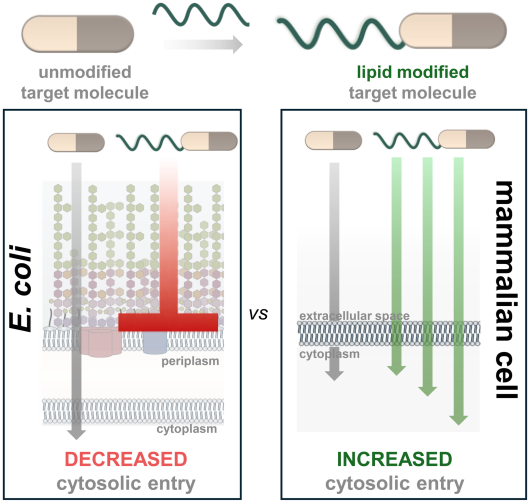

## INTRODUCTION

A central obstacle in drug development is the cell envelope, which restricts delivery of biologically active molecules to the cytosol [1–4]. While this challenge is shared by mammalian and bacterial systems, it is especially pronounced in Gram-negative pathogens such as ***Escherichia coli* (*E. coli*),** whose dual-membrane envelope, low-permeability outer membrane, and active efflux together limit antibiotic access to the cytosol [5–8]. Understanding the parameters that govern membrane permeability remains a major challenge, owing to a shortage of methods that distinguish cytosolic arrival from bulk cell association [9–11]. There is thus a need for widely adoptable, high-throughput methods that measure cytosolic accumulation across cell types to guide drug development [12–13].

Natural products frequently carry structural modifications that promote membrane permeation; lipid conjugation in particular increases hydrophobicity, favoring partitioning into the bilayer and, in many cases, access to the cytosol (**Fig. 1a**) [14–15]. A recent chemoinformatic analysis of the natural product lipidome by the Trauner group revealed that lipidated natural products predominantly carry medium-chain lipids (MCLs), which are appreciably shorter than the long-chain lipids found in membranes and lipidated proteins [16]. Building on this, they conjugated a series of small-molecule chemotypes (including fluorescent dyes and model drugs) to lipid tails of varying length and showed that medium-chain conjugates can render otherwise impermeable molecules cell-permeable while modulating their subcellular localization and bioactivity [16]. Such modifications appear across diverse drug classes that must cross the cellular envelopes of different organisms. For instance, bioactive natural products with lipid tails include myriocin [17], kahalalide F [18], icaritin [19], fidaxomicin [20], and andrimid [21], which engage cytosolic targets. The prevalence of lipid pharmacophores has motivated similar strategies in drug development, with lipidation becoming an increasingly common way to facilitate delivery into the cytosol [22–30]. Lipidated natural products that act on cytosolic targets in Gram-negative pathogens are notably rare (**Fig. 1b**), a prominent exception being the lipopeptides such as polymyxin B, which act at the outer membrane rather than in the cytosol. Whether this scarcity reflects poor cytosolic accumulation, restricted target space, or discovery bias remains unresolved [8].

**Figure 1.**
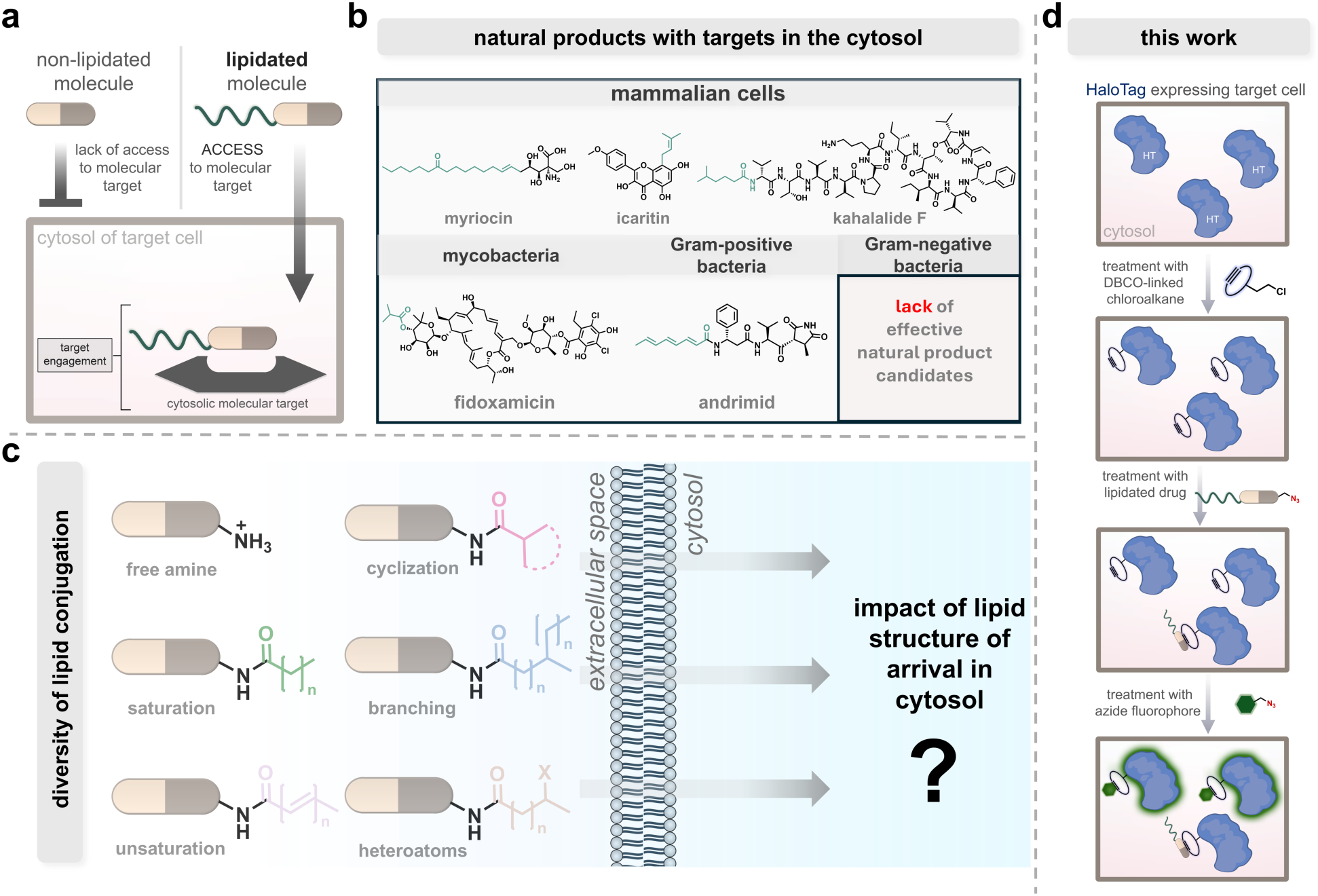
**a**, Schematic representation of the idea that lipidation of a target drug molecule enhances arrival into the cytosol and subsequent access to intracellular targets. In addition, a table identifying various natural products with cytosolic targets amongst a variety of organisms highlighted by the lipid pharmacophore. **b,** Schematic highlighting the differences in lipid structure and the gap in knowledge of how structure of the lipid moiety impacts permeation through a membrane. **c,** Synthetic pathway of the scaffold, **P0**, and scale of the lipid library used for the study. Lipid variations range from simple chain length (**Sat**), presence of double bonds (**Unsat**), cyclization (**Cyc**), presence of heteroatoms (**Het**), branching (**Brn**), and mixture of these different modifications (**Mix**). **d,** Overall workflow of CHAMP, symbolic of both mammalian cells (HeLa) and Gram- negative bacteria (*E. coli*). Cells carrying a plasmid that expresses HaloTag in the cytosol are treated with a strained alkyne dibenzocyclooctyne (DBCO) that effectively anchors the target azide-modified molecule to the cytosol. This “pulse step” can be used to assess the level of cytosolic arrival, which is subsequently followed by a “chase step” with an azide-modified fluorophore to anchor to any free DBCO remaining in the cytosol. In the case of molecules with high levels of accumulation, the cellular fluorescence level is expected to be low. The opposite should be observed for molecules with low levels of cytosolic accumulation.

Despite the prevalence of lipid modifications and the assumption that they drive accumulation, methods to empirically measure their impact on small-molecule arrival in the cytosol remain lacking [13]. Current methods of measuring cytosolic accumulation in different systems vary in their advantages and drawbacks. In bacteria, the reference method is liquid chromatography–tandem mass spectrometry (LC-MS/MS) [31], which is advantageous because it requires no chemical tag [32–33]. However, the low-throughput nature of this method limits its utility, especially in addressing the urgency of the antimicrobial resistance crisis [34,35]. Furthermore, measurements made on cell lysates cannot distinguish cell-associated molecules from those free in the cytosol, increasing the rate of false-positive hits. In mammalian cells, more standardized permeability assays, such as the Parallel Artificial Membrane Permeability Assay (PAMPA) and the Caco-2 assay, while successful, suffer from similar limitations of low throughput and poor spatial resolution [36–39].

To address these limitations, molecular tagging has emerged as a powerful strategy for visualizing intracellular accumulation [40–41]. Fluorescently tagged molecules can be visualized by confocal microscopy, which improves spatial localization [42]. Recently, the Kritzer lab developed the chloroalkane penetration assay (CAPA), in which cells are pulsed with a chloroalkane-tagged molecule that reacts with a cytosolic HaloTag self-labeling protein and then chased with a chloroalkane-tagged dye [43]. This approach is intrinsically compatible with flow cytometry, which substantially improves throughput and enables readout on a per-cell basis. CAPA has also been adapted to bacterial systems, expanding the range of membrane microenvironments that can be studied [44–46]. One consideration when applying CAPA is the size of the chloroalkane tag, which for small parent molecules can constitute a substantial fraction of the conjugate and may itself contribute to accumulation [47]. CHAMP was designed to reduce this contribution while retaining the per-cell, flow-cytometry readout that CAPA established.

Whether the lipidation patterns reported for mammalian cells translate to diderm bacteria has not been tested directly (**Fig. 1c**). To address this, we applied the Chloroalkane Azide-based Membrane Penetration (CHAMP) assay (**Fig. 1d**), a high-throughput platform recently developed to quantify cytosolic accumulation *via* strain-promoted azide-alkyne cycloaddition (SPAAC) [48–51]. Using a minimally disruptive azide tag that covalently reacts with a dibenzocyclooctyne (DBCO) moiety anchored in the cytosol by a HaloTag protein, CHAMP limits the confounding effects of larger tags and separates true cytosolic arrival from membrane binding. Here, we deploy CHAMP in both HeLa and *E. coli* cells to systematically investigate the cytosolic accumulation of lipid conjugates, derived from a hydrophilic parent molecule, which vary in structure and physicochemical properties. By comparing these two distinct biological contexts, we sought to determine whether the membrane-partitioning and translocation advantages conferred by lipidation translate into a universal, cross-domain principle for facilitating molecular access to the cytosol.

## RESULTS

### A Minimal Reporter Scaffold Isolates the Contribution of the Lipid Tail

To ensure that observed changes in cellular behavior could be attributed to the conjugated lipid rather than to the scaffold bearing it, we designed a minimal baseline reporting scaffold, **P0**, prepared by solid-phase synthesis (**Fig. 2a**) [16,52]. We note at the outset that **P0** carries a free, protonated N-terminal amine, whereas every lipid conjugate carries a neutral amide; comparisons to **P0** therefore report the combined effect of acylation and loss of the terminal ammonium. For this reason, **Sat1** (the acetylated congener) serves as the parent for all comparisons of lipid structure, and **P0** is used only as a non-lipidated baseline. Minimizing the non-lipid portion of the molecule limits its intrinsic contribution while retaining three functional motifs: (1) an aniline chromophore for quantification of concentration by UV/Vis spectroscopy, (2) an azide tag required for the CHAMP assay, and (3) a free *N*-terminal amine for on-resin acylation with the various lipid tails [48–49, 53]. The first two motifs provide the analytical readouts, and the third serves as the site of lipid attachment, such that lipid structure constitutes the sole variable across the series. Using this scaffold, we synthesized and screened a library of lipid conjugates incorporating a structurally diverse set of lipids that varied in saturated chain length, unsaturation, cyclization, branching, and the presence of heteroatom-containing functional groups (**Fig. 2b**). Many of these lipids overlapped with those examined in the Trauner study, allowing the corresponding mammalian-cell findings to be corroborated [16]. **P0** also allowed us to examine acylation of a free amine, a transformation common among natural products and in biological systems [23, 54].

**Figure 2.**
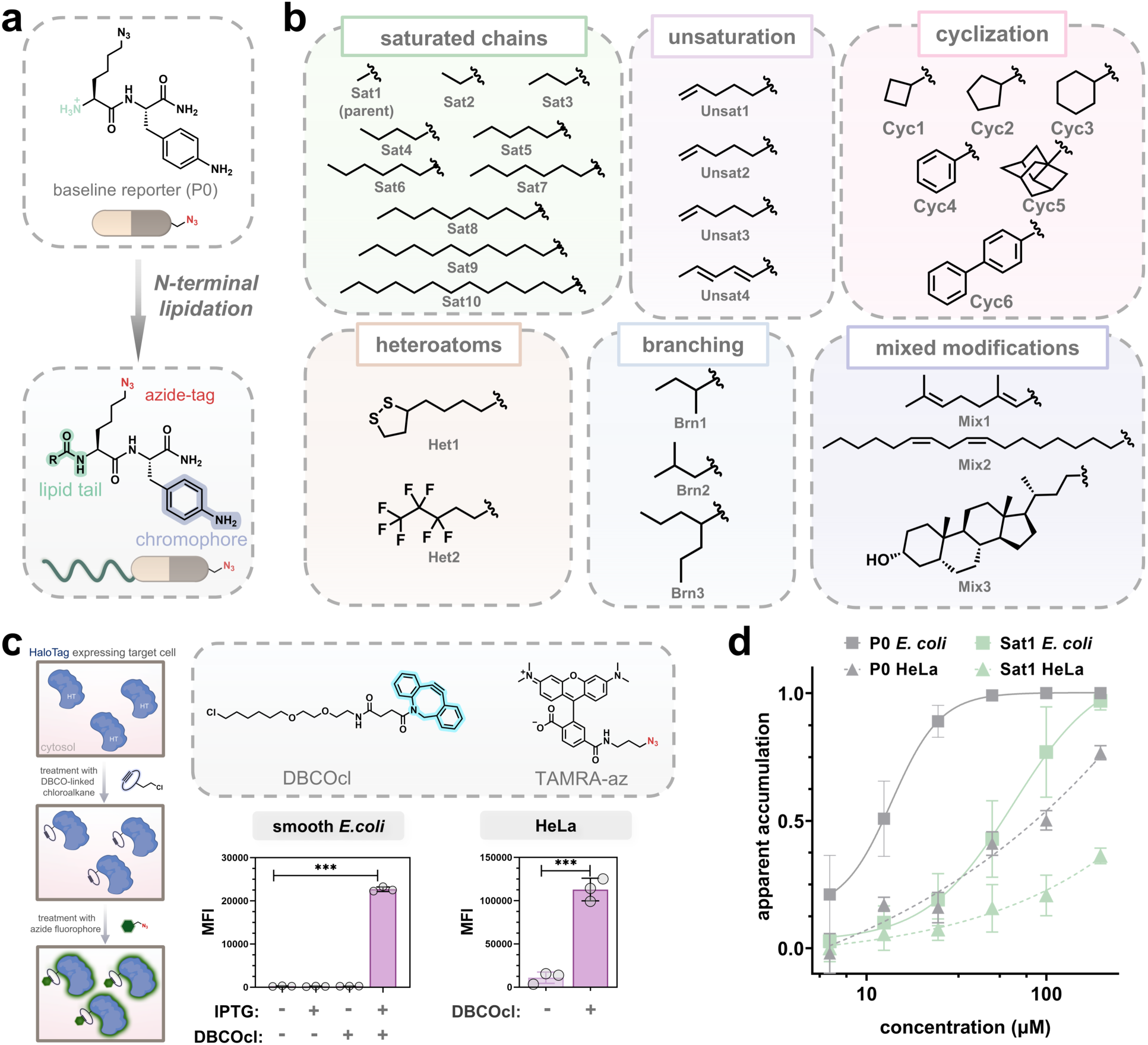
**a**, Synthetic scheme describing the baseline reported scaffold, **P0** and the eventual lipid derivatives. The notable features of the compound, including the azide-tag, aniline chromophore, and modifiable N-terminus are highlighted and give the scaffold its intrinsic advantages. **b,** Structures of the library of lipid tails used in this study along with their classification into the various subsets. **Sat1** is denoted as the parent molecule of the library **c,** Schematic representation of the one-step labeling process used to verify HaloTag expression within cells, paired with structures of **DBCOcl** and **TAMRA-az.** Testing in *E. coli* revealed the necessity for both IPTG and **DBCOcl** to gain a signal from eventual anchoring of the **TAMRA-az,** while in HeLa cells, in which HaloTag is constitutively expressed, it indicated the need for **<u>DBCOcl</u>** for anchoring of azide- molecules to the HaloTag. Signal-to-noise ratios in both systems were deemed optimal for subsequent usage. **d,** CP_50_ curves of **P0** and **Sat1** in both *E. coli* and HeLa cells across various concentrations during a 1 hr incubation time. Data are represented as mean +/− SD (*n* = 3) of biological replicates. *P*-values were determined by a two-tailed *t*-test (* denotes a *p*-value < 0.05, ** < 0.01, ***<0.001, ns = not significant).

Compatibility with the CHAMP assay is necessary to ensure that our scaffold and its derivatives provide a reliable platform for measurement. We isolated two such features that may affect CHAMP assay performance: reactivity and aggregation. Because CHAMP depends on SPAAC proceeding to completion during the incubation, differences in azide reactivity across the library could masquerade as differences in accumulation. We therefore benchmarked azide reactivity using DBCO-modified beads in a workflow analogous to the two-step CHAMP assay (**Fig. S1**) [48–49]. Conjugating a lipid tail can also promote self-association into micelles or colloidal aggregates [55]. We assessed whether this would be present within the library through fluorescence testing with 1- anilinonaphthalene-8-sulfonic acid (ANS) (**Fig. S2**). ANS reports on aggregation by binding to hydrophobic pockets exposed upon self-assembly, producing a marked increase in fluorescence intensity [56–58]. Across the concentration range used here, no compound produced the fluorescence enhancement characteristic of aggregate formation. We note that ANS reports on solvent-exposed hydrophobic pockets and is not a direct measure of colloidal state; for the longest chains in the series we therefore interpret this result as evidence against extensive aggregation rather than as proof of a purely monomeric state. Together, these controls indicate that the library is suitable for screening.

Previous work in our laboratory has established a standard protocol for the CHAMP assay in both WT smooth *E. coli* and HeLa cells [48–49] (**Fig. S3**). HaloTag expression and incubation conditions have been benchmarked previously in both systems: cells are exposed to the test compound during the pulse step at a fixed concentration of 50 µM for 1 hr at 37°C. Subsequent treatment with the azide-tagged fluorophore occurs under the same conditions as the pulse step. To ensure consistency between the systems, we used **TAMRA-az** as the assay readout, as it is compatible with the workflow in both systems. We confirmed its utility with a two-step labeling process, verifying that the system was functional and gave a strong signal-to-noise ratio (**Fig. 2c**). HaloTag expressing *E. coli* were treated with **DBCOcl** to anchor the cytosolic landmark, DBCO. This yielded a >100- fold signal-to-background ratio upon covalent capture of **TAMRA-az**. In HeLa cells, **DBCOcl** was likewise required to anchor the fluorophore, giving a >30-fold signal-to-background ratio. We note that this ∼3-fold difference in dynamic range between the two systems is one reason we compare compounds within a system rather than across systems (see below). Establishing these parameters confirmed that both systems were suitable for analysis of the library.

A second requirement is responsiveness to concentration. To serve as a measure of cytosolic arrival, the assay must respond to changes in compound concentration, from which a half-maximal accumulation concentration (CP_50_) can be derived to quantify compound-specific permeability [59–60]. To confirm that both systems were responsive, we performed concentration scans using the two-step CHAMP protocol with the baseline compound **P0** and the acetylated parent **Sat1** (**Fig. 2d**). Bearing only an acetyl moiety, this structure more closely mimics the rest of the library, allowing us to separate the effect of acylation itself from the effect of lipid tail structure [16]. To aid interpretation, the data were processed as “1 − (normalized fold change in fluorescence intensity)” to yield a value we term “apparent accumulation,” which positively correlates with cytosolic accumulation [48–49]. For a small number of compounds, treated samples gave fluorescence above the untreated (maximum-signal) reference, most plausibly reflecting partial membrane permeabilization that increases dye entry. Because apparent accumulation is calculated as 1 − normalized fluorescence, these values can yield apparent accumulation below zero. Such values reflect the limits of the fluorescence readout and are interpreted as minimal or undetectable accumulation rather than true negative accumulation. Negative values were censored to zero before any correlation or ratio analysis. The two scaffolds behaved differently in the two systems. In *E. coli*, both **P0** and **Sat1** reached half-maximal accumulation (CP₅₀) at lower concentrations than in HeLa cells. Acylation of **P0** to give **Sat1** reduced accumulation in both systems, although in *E. coli* this drop conflates the added acetyl group with the loss of the N-terminal ammonium. That small, polar scaffolds accumulate efficiently in *E. coli* is consistent with the physicochemical constraints reported for the diderm envelope [61–63], and raised the possibility that the two systems respond differently to lipidation.

### Lipid Structure Drives Opposite Accumulation Trends in the Two Systems

Lipidation has been studied as a permeability-enhancing strategy in mammalian cells, but those readouts largely tracked cellular uptake and bioactivity and could not definitively distinguish true cytosolic arrival from membrane association [16]. No comprehensive counterpart exists for Gram-negative bacteria, to our knowledge. The expectation is that increased hydrophobicity aids desolvation and promotes partitioning into the hydrophobic membrane environment [14–15]. Yet this reasoning derives largely from the single lipid bilayer of the mammalian plasma membrane. The asymmetric, LPS-rich outer membrane of Gram-negative bacteria presents a chemically distinct interface and may impose different requirements [64–66]. Because CHAMP reports cytosolic arrival directly in both systems, the library allowed us to test this expectation.

Screening the library at a single concentration in both systems revealed clear differences in how lipid structure relates to cytosolic accumulation (**Fig. 3a**). Relative to **Sat1**, the acetylated parent for the series, extending or elaborating the lipid tail generally reduced accumulation in *E. coli*; no conjugate recovered most of the accumulation seen for the non-acylated baseline **P0**. Only **Sat2** and **Sat3**, the shortest-chain conjugates, exceeded **Sat1**; every longer chain and every added structural motif reduced accumulation. The relationship with chain length is therefore biphasic, with an optimum at C3–C4 rather than a monotonic preference for the shortest tail. This runs counter to the expectation that greater hydrophobicity aids permeation, an expectation drawn largely from monolayer-bilayer systems [64–66].

**Figure 3.**
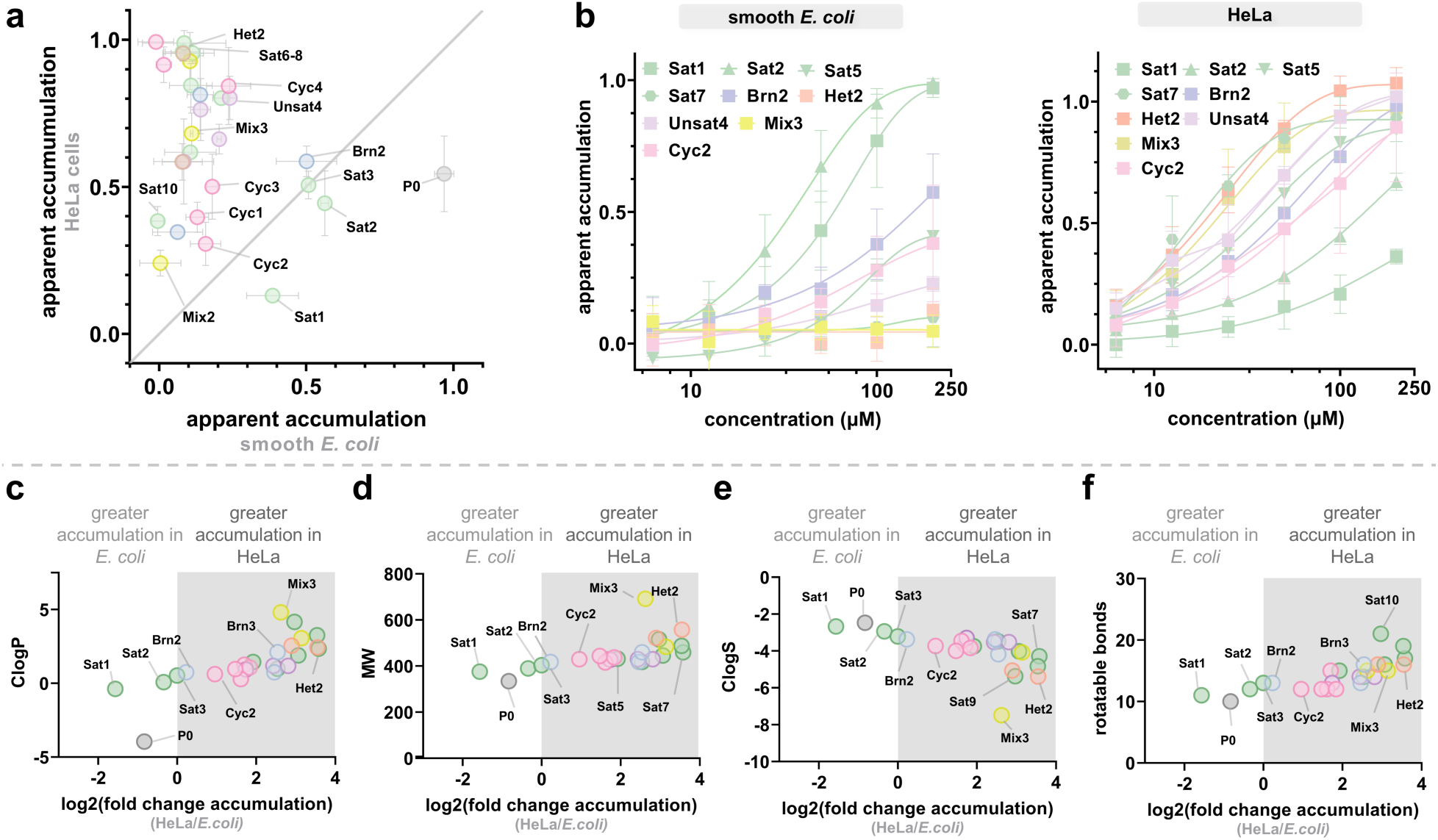
**a**, Graphical representation of the accumulation of the library members between HeLa cells (y-axis) and smooth *E. coli* (x-axis). Data represents single-point, apparent accumulation values derived from a treatment of 1 hr at 50 μM of compound. All compounds and their normalized fluorescence intensity can be found in the supplementary data (**Fig. S6** and **Fig. S9**). **b,** CP_50_ curves are generated for a select group of compounds in both smooth *E. coli* and HeLa cells. Concentration **c-e,** Comparison of accumulation of the library between HeLa and *E. coli* based on physicochemical parameters impacted by lipidation: **c** partition coefficient (ClogP), **d** molecular weight (MW), **e** aqueous solubility (ClogS), and **f** rotatable bonds. Data was normalized by taking the log2(fold accumulation difference), with “fold accumulation difference” being the normalized apparent accumulation in HeLa divided by that of *E. coli.* Data are represented as mean +/- SD (n = 3) of biological replicates.

To rule out complete membrane partitioning as a contributor, we used two fluorescence-based reporters of membrane integrity, NPN and SYTOX Green [67]. N-phenyl-1- naphthylamine (NPN) is excluded by the intact LPS outer leaflet; when the outer membrane is perturbed, NPN partitions into the bilayer and its fluorescence increases [68]. Under typical CHAMP conditions, we measured NPN fluorescence over the standard incubation period (**Fig. S7**). Relative to untreated cells, only **Sat10** and **Mix3** produced a measurable increase in NPN fluorescence, and both remained far below the polymyxin B positive control [67]. Because these two compounds also feature in the size- and hydrophobicity-dependent trends described below, we treat their apparent accumulation values as potentially confounded by membrane perturbation and interpret them with caution. SYTOX Green, which enters only cells with a compromised cytoplasmic membrane and fluoresces upon binding nucleic acids, showed no significant signal for any library member, indicating that the inner membrane remained intact (**Fig. S8**) [69]. We speculate that larger, lipid-containing molecules embed in the outer membrane, a common feature of lipidated species in nature and drug discovery, which may prevent their arrival in the cytosol [70–71].

We observed different accumulation patterns in HeLa cells, which were expected given the weaker accumulation of **P0** compared to *E. coli*. Within HeLa cells, **Sat1** ranked among the weakest accumulators, whereas in *E. coli* it ranked among the strongest, the first indication that the two systems order the same compounds differently. Consistent with the findings of Morstein et al. [16], the medium-chain saturated lipids **Sat6**, **Sat7**, and **Sat8** accumulated most strongly in HeLa cells, relative to both shorter and longer chains. [16]. For the flexible saturated cyclic lipids (excluding the aromatic and adamantane scaffolds), larger rings generally accumulated more in HeLa cells, paralleling the chain-length trend in the saturated series. Likewise, the more rigid rings, including the phenyl groups in **Cyc4** and **Cyc6** and the adamantane in **Cyc5**, generally correlated with stronger accumulation in mammalian cells. Notably, this rigid, cage-like scaffold was among those that did not enhance uptake in the earlier study [16], suggesting that a bioactivity- or association-based readout and a direct cytosolic-arrival readout can rank the same modification differently. Whereas cyclization largely preserved accumulation relative to the straight-chain counterparts, unsaturation of a straight-chain lipid tended to impede it. Comparison of the unsaturated series with its parent, **Sat5**, revealed that one or more double bonds generally restricted accumulation in HeLa cells; **Unsat1** was the only member to retain most of the parent’s accumulation. Branching the linear chain, whether small as in **Brn1** and **Brn2** or larger as in **Brn3**, hindered accumulation in *E. coli* but conversely increased accumulation in HeLa compared to the parent linear chain (**Sat3** and **Sat4**, respectively). **Het2**, the polyfluoroalkyl analogue of **Sat5**, accumulated strongly in HeLa cells and weakly in *E. coli*. This result is consistent with previous reports suggesting that similar modifications can aid cytosolic accumulation of larger peptides [72–73]. Similarly, **Mix3**, which is modified with a lithocholic acid (sterol) moiety, selectively favored permeation into mammalian cells. Sterol conjugation has been reported to enhance membrane association and, in some systems, uptake [74, 75]. We note, however, that **Mix3** also produced a small increase in NPN fluorescence, so this result should be treated as provisional. Overall, we observed that permeation into *E. coli* tolerates lipid modification far less than permeation into HeLa cells, which accommodate a wide range of lipid structures that enhance intracellular accumulation.

To further the analyses and isolate key quantitative differences between *E. coli* and HeLa lipid-driven accumulation, we generated CP_50_ curves with a select group of compounds from the library (**Fig 3b**) [43]. This subset was chosen to span every structural class in the library while including the strongest accumulators in each system. Across a concentration range of 6.25–200 µM, the selected compounds differed markedly in their concentration dependence. In *E. coli*, the small, polar moieties of **Sat1-2** showed greater than 2–4-fold accumulation compared to the rest of the conjugates. Within this subset, **Sat2** showed a CP_50_ ∼ 50 µM while **Sat1** was measured ∼ 100 µM. In this concentration range, the rest of the compounds were unable to reach saturation, but **Cyc4** and **Brn2** again show that smaller modifications partially preserve accumulation in *E. coli*. In HeLa cells, medium chain moieties like **Sat6-7** and **Het2** show the overall strongest accumulation patterns across the concentration range. Comparing **Het2** to parent compound **Sat5** reemphasizes the nature of fluorination as a means of increasing cytosolic arrival of a hydrocarbon moiety [72–73]. Cyclization of **Sat5,** in the form of **Cyc2**, was significantly unfavorable for accumulation and recorded the highest CP_50_ of all those tested. **Brn2** showed the smallest rank difference between the two systems, though it still favored HeLa cells. **Sat2** was the only compound that ranked higher in *E. coli* than in HeLa, supporting the broader pattern that small, polar conjugates are preferred by *E. coli* whereas larger, more hydrophobic conjugates are preferred by mammalian cells.

Having now deciphered the differences in intracellular accumulation between HeLa cells and *E. coli,* we next sought to determine the physicochemical properties which may favor accumulation in one system or the other. These properties encompass several molecular parameters which are commonly impacted by the addition of a lipid moiety (**Fig 3c-3f)**, including lipophilicity (ClogP), aqueous solubility (ClogS), molecular weight (MW), and rotatable bonds [48–49, 76–77]. Upon calculating these properties for the library, we found a significant positive correlation of accumulation in HeLa cells with ClogP, molecular weight, and rotatable bonds, indicating that larger, more hydrophobic molecules are favored to cross the mammalian cell envelope (Spearman’s ρ = 0.71, 0.83, and 0.69, respectively). Consistently, accumulation correlated negatively with ClogS (Spearman’s ρ = −0.84), reinforcing that hydrophobicity and size help guide a molecule through the mammalian membrane. Interestingly, the presence of heteroatoms, such as in **Het1** and **Het2,** regardless of their effects on the physicochemical properties of the molecules, selectively favored permeation in HeLa cells. For *E. coli*, these data support the divergent rules for intracellular accumulation, with small, polar molecules favored to traverse the cell envelope [64–66].

### Evaluating Structural Changes to the Molecule and their Impacts on Accumulation

With a clearer understanding of how lipid structure governs accumulation, we next examined how the lipid group modulates the effect of changes to the small-molecule scaffold. We approached this from two directions: first altering the lipid tail while holding the scaffold constant, then editing the scaffold while holding the lipid tail constant. We began with the straight-chain tail of Sat7, which ranks near the top of the HeLa series and near the bottom of the *E. coli* series. We note that this positioning limits the dynamic range available in each system: in HeLa cells, modifications can readily be scored as detrimental but improvements are difficult to resolve, and the converse holds in *E. coli*. We therefore interpret this dataset as directional rather than quantitative, and treat CP₅₀ determination for the most informative analogues as a priority for follow-up. We reasoned that altering the charge at the end of the tail could add functionality to the lipid which, given the difference in membrane potential between HeLa cells and *E. coli* cells, could achieve selective uptake into cells [78–81]. To accomplish this, we used commercially available octanoic acid derivatives with either a terminal carboxylic acid (**7b**) or amine (**7c**) to assess both positive and negative charges (**Fig. 4a**). We also incorporated an 8- azidooctanoic acid derivative (**7d**) to examine whether accumulation depends on the position of the azide moiety. Azide reactivity was unchanged after swapping the amine and azide positions (**Fig. S11**), so differences in this dataset can be attributed to permeability rather than to ligation efficiency. In *E. coli*, installing a terminal primary amine (**7c**) did not measurably alter accumulation relative to **Sat7**. Although prior work suggests that a primary amine can improve accumulation in diderm bacteria [33], we observed no such effect here, likely because **Sat7** already accumulates poorly in *E. coli*. Both **7b** and **7d** accumulated poorly in E. coli. For the anionic **7b**, this is consistent with the inside- negative inner-membrane potential of *E. coli*, which opposes anion entry [70–71]. **7d** is not explained by this argument, and we therefore do not attribute the shared behavior of **7b** and **7d** to membrane potential alone. In HeLa cells, all modifications to **Sat7** curtailed accumulation, further reaffirming previous findings that hydrophobicity is a guiding principle towards accumulation in mammalian cells.

**Figure 4.**
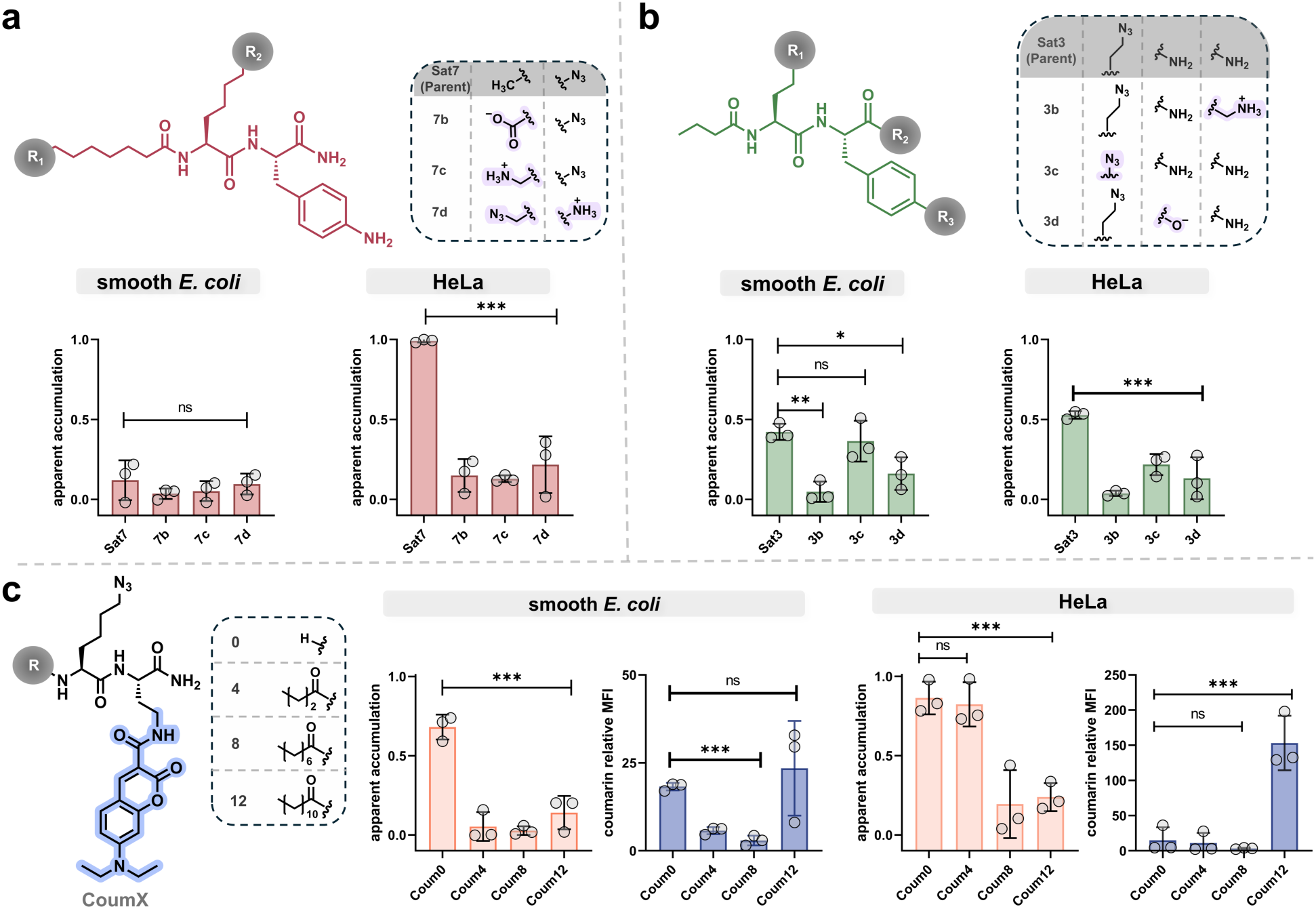
**a**, Chemical structure of **Sat7** and the several modifications to the lipid tail covered in the structure-activity relationship along with CHAMP data on the apparent accumulation in both *E. coli* and HeLa. **b,** Similar to **a,** structure of **Sat3** and the map of modifications to the endogenous portions of the molecule. Similar CHAMP data was acquired for assessing the library. **c,** Structure of the 7-diethylaminocoumarin derivative (**CoumX**) along with the short list of lipid tails covered within this examination. In both *E. coli* and HeLa cells, the apparent accumulation along with relative mean fluorescence intensity (MFI) from the coumarin moiety. Relative mean fluorescence intensity was calculated from normalizing to the background signal of the system. Data represents single-point, apparent accumulation values derived from a treatment of 1 hr at 50 μM of compound. All compounds and their normalized fluorescence intensity can be found in the supplementary data (**Fig. S11**) Data are represented as mean +/- SD (n = 3) of biological replicates. *P*-values were determined by a one-way ANOVA for **a** and **b** while a two-tailed *t*-test was employed for **c** (* denotes a *p*-value < 0.05, ** < 0.01, ***<0.001, ns = not significant).

**Sat3** accumulated better than **Sat1** in both systems while sitting mid-range in each, making it a suitable parent for testing whether a lipid tail can offset changes elsewhere in the molecule. We reasoned that targeted changes to the two-residue scaffold (**Fig. 4b**), namely increasing the basicity of the amine (**3b**), reducing the surface area of the azido- lysine (**3c**), and converting the *C*-terminus to a carboxylic acid (**3d**), would test whether the lipid tail could compensate for these alterations [48–49]. In both systems, almost all modifications led to a significant decrease in accumulation, indicating that the lipid moiety cannot compensate for general changes to the molecular scaffold. The most tolerated change was shortening the azidolysine side chain in **3c**, which partially restored accumulation in *E. coli*, suggesting that a reduction in polar surface area can offset some of the penalty imposed by the tail. Overall, these results indicate that membrane permeation is sensitive to changes in the compound that increased hydrophobicity alone cannot offset.

Previous work used a direct readout based on lipid-conjugated dyes to estimate cytosolic accumulation [16]. A readout of this kind reports total cellular fluorescence, which combines cytosolic and membrane-associated compound; the two contributions cannot be separated without an independent measure of cytosolic arrival. We therefore used solid-phase synthesis to install an intrinsic 7-(diethylamino)coumarin fluorophore in our scaffold as a direct readout for flow cytometry. Using a small subset of saturated chains of varying length (**Coum0, Coum4, Coum8,** and **Coum12)** we aimed to further validate our previous findings with a direct anchoring mechanism (**Fig. 4c**) [48–49]. In *E. coli*, the overall accumulation trends paralleled those in the main library. By the CHAMP readout, the polar, non-lipidated **Coum0** showed the highest apparent accumulation. The direct fluorescence readout behaved in the opposite manner, rising with chain length, a pattern most consistent with membrane partitioning rather than cytosolic arrival [82–84]. In HeLa cells, the strongest accumulators were **Coum0** and **Coum4**, consistent with previous data suggesting that increasing molecular size can shift the optimal lipid tail for permeability [16]. Interestingly, both showed relatively low fluorescence compared with **Coum8** and **Coum12**, which showed much larger fluorescence increases with decreasing permeability. Across this series, total cellular fluorescence and CHAMP-reported cytosolic arrival were inversely related: the compounds with the greatest whole-cell signal were the weakest cytosolic accumulators. This is the central distinction the assay is designed to make: whole-cell readouts, whether fluorescence- or mass spectrometry-based, report the sum of membrane-associated and cytosolic material, whereas CHAMP reports only the fraction that reaches the cytosol and is covalently captured there.

### Perturbing Individual Envelope Layers Identifies the Inner Membrane as a Dominant Barrier

After assessing modifications to the test molecule itself, we sought to understand how lipidation recovers or potentially facilitates accumulation across a variety of exogenous changes to the system. We chose to focus mostly on *E. coli*, given the range of available modifications to alter the cell envelope structure and function [48]. We began by assessing whether the presence of the polar O-antigen moiety on *E. coli* affected accumulation of the library and whether any motifs could overcome it (**Fig. 5a**). Prior work suggests that the O-antigen displayed on smooth *E. coli* can act as an additional barrier to nonpolar molecules entering the cytosol [85–86]. Using a HaloTag expressing strain of rough *E. coli* (BW25113), we compared apparent accumulation to isolate the effects of the O-antigen. No library member showed a substantial change between the smooth and rough strains, apart from small increases for the smallest, most polar conjugates. We note that BW25113 is not an isogenic O-antigen-deficient derivative of the smooth strain, so this comparison also carries background differences; we therefore read it as a negative result (the O-antigen is not the principal barrier to these compounds) rather than as evidence that it is irrelevant.

**Figure 5.**
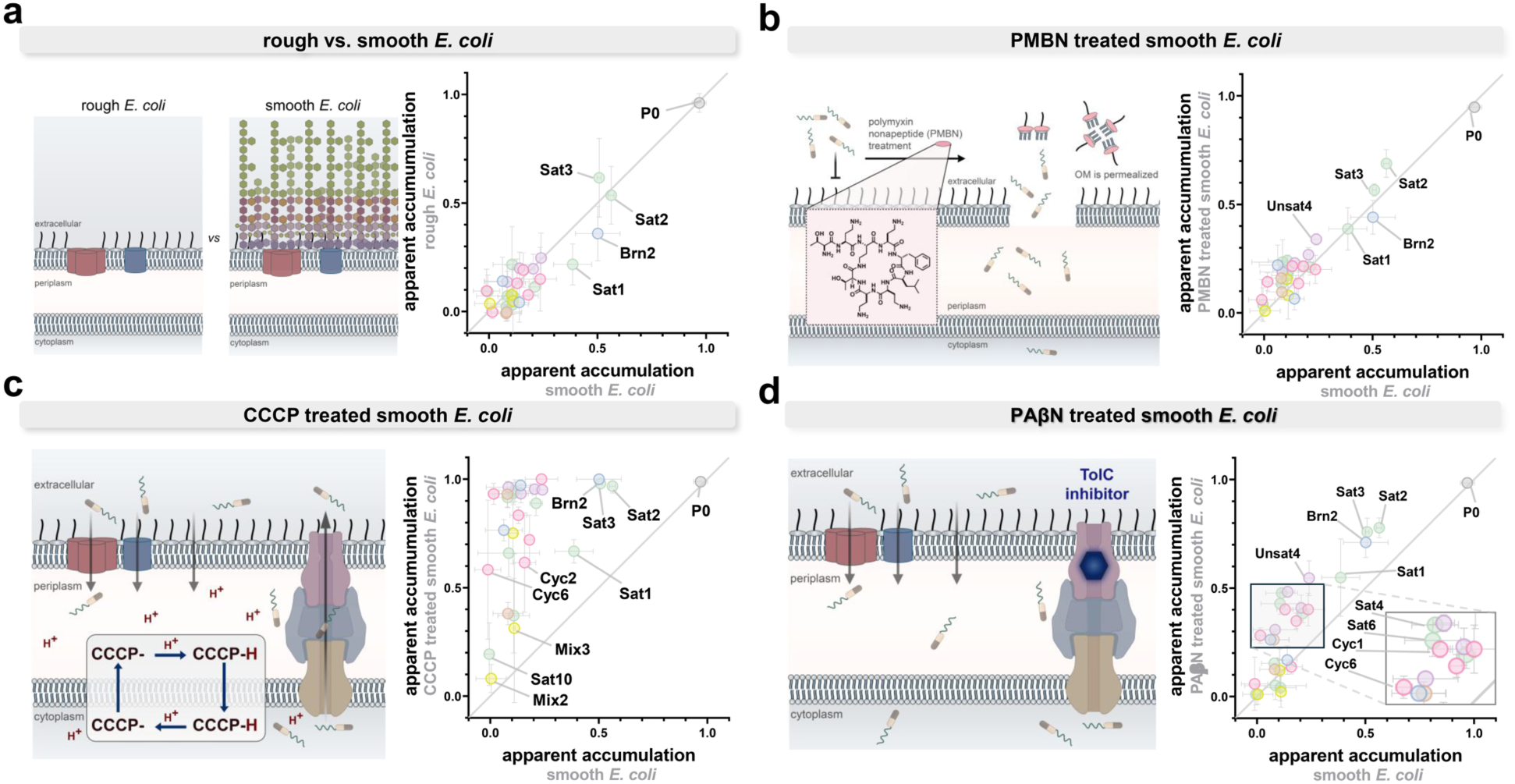
**a**-**d,** Schematic representations of the various exogenous modifications conducted in the *E. coli* CHAMP and graphical data showing differences in accumulation within the library as compared to the smooth *E. coli* control group. Data represents single-point, apparent accumulation values derived from a treatment of 1 hr at 50 μM of compound. **a,** Comparison between the presence or absence of the O-antigen on the lipopolysaccharides (LPS) using HaloTag expressing rough *E. coli* (BW25113). **b,** Co-treatment of smooth *E. coli* with 25 μg/mL polymyxin B nonapeptide (PMBN) to effectively permeabilize the outer membrane to influx of the lipidated molecules. **c,** Pre-treatment of smooth *E. coli* with 100 μM Carbonyl cyanide m-chlorophenylhydrazone (CCCP) for 10 min to disrupt the proton motive force in the inner membrane and prevent efflux of the lipidated series. **d,** Co-treatment of *E. coli* with 16 μg/mL of the AcrAB-TolC inhibitor Phenylalanine Arginine Beta-Naphthylamide (PAβN) to isolate the impact of efflux pumps on cytosolic arrival of the lipidated series. Data are represented as mean +/- SD (n = 3) of biological replicates.

Next, we assessed permeabilization of the outer membrane using polymyxin B nonapeptide (PMBN), a less disruptive derivative of polymyxin B that permeabilizes the outer membrane and has been shown to increase the biological activity of various antibiotics [68,87]. Here, we co-incubated the library with PMBN to isolate the effects of the outer membrane as a potential barrier to cytosolic accumulation (**Fig. 5b**) [88–90]. Interestingly, PMBN showed little to no effect on aiding most conjugates except for small, polar lipid conjugates such as **Sat1** and **Brn2**. Therefore, we infer that outer membrane permeabilization alone is not enough to facilitate cytosolic accumulation of increasingly hydrophobic molecules. In a similar manner, we investigated the effects of hyperporination on accumulation using a strain of rough *E. coli* expressing a mutant FhuA porin protein which acts as a constitutively open pore on the outer membrane. We note that for these experiments, comparisons were made to the parent rough strain [48, 91–93]. CHAMP comparisons made here (**Fig. S12**) only showed **Cyc4, Brn2**, and **Unsat4** as being aided by the presence of the pore. Given the difference in *E. coli* strains used in these outer-membrane modifications, it is difficult to determine why the accumulation patterns diverged. Nonetheless, these results indicate that the outer membrane can still act as a major barrier to permeation of lipidated small molecules.

To probe the inner membrane as a barrier to intracellular accumulation, we first targeted its energy-dependent defenses. We collapsed the inner-membrane proton motive force (PMF) by pretreating smooth *E. coli* with carbonyl cyanide m-chlorophenylhydrazone (CCCP) [66,94]. The effects of CCCP on permeability are debated, but the prevailing view holds that it depolarizes the inner membrane while inactivating efflux pumps [95]. Quantification by CHAMP (**Fig. 5c**) showed that PMF disruption increased apparent accumulation for nearly the entire library. The compounds that responded most strongly were the medium-chain conjugates, suggesting that their exclusion under normal conditions is imposed at the inner membrane, through the potential-dependent barrier, active efflux, or both, rather than at the outer membrane. The differential response of larger, structurally complex compounds like **Sat9** and **Cyc5** indicated that extreme size remains detrimental to permeation even in a severely compromised, depolarized state. While this implicates the inner membrane and its associated active efflux systems as critical bottlenecks, the pleiotropic effects of PMF disruption, including global depolarization and efflux inhibition, preclude isolating the inner membrane’s passive physical barrier alone. While promising, further experiments would be needed to establish the inner membrane as the major contributor to bacterial defense against small-molecule permeation. We next asked whether efflux pumps contribute to the low apparent accumulation of the compounds. To test this, we co-incubated the cells with compounds in the presence or absence of phenylalanine-arginine β-naphthylamide (PAβN) (**Fig. 5d**). PAβN is a known inhibitor of the AcrAB-TolC efflux pump and has been shown to increase the accumulation of several antibiotics such as fluoroquinolones [96–98]. From the library scan, we once again observed short to medium lipid conjugates such as **Sat6, Brn2,** and **Unsat4** experiencing increased cytosolic accumulation, suggesting that these compounds do reach the cytosol but are effluxed by the AcrAB–TolC system [99–100]. Taken together, these envelope perturbations identify which structural classes are limited by which barrier: small polar conjugates by the outer membrane, and medium-chain conjugates by the inner membrane and efflux.

## DISCUSSION

Lipidation is a recurring solution to the problem of membrane permeation, but its use is markedly uneven across biology: lipidated natural products with cytosolic targets are abundant in eukaryotes and monoderm bacteria, and conspicuously rare in Gram-negative pathogens. The lipopeptide antibiotics are the instructive exception (polymyxin B and colistin) are lipidated, potent against Gram-negatives, and act at the outer membrane rather than in the cytosol. What has not been established is whether the rarity of cytosolically acting lipidated antibacterials reflects a permeability constraint, and if so which layer of the envelope imposes it. Prior systematic studies of accumulation in Gram-negative bacteria have examined lipophilicity as a variable, but have relied on whole-cell methods (principally LC-MS/MS of lysates) that cannot separate cytosolic material from compound associated with the envelope [101–102]. Methods such as the CHAMP assay allow us to parse out the differences in cytosolic exposure and membrane localization across mammalian cells and Gram-negative bacteria [48–49]. Here we used that capability to survey lipid modifications spanning chain length, unsaturation, ring size, branching, and heteroatom content in two cell types with the same readout. On the bacterial side, previous studies that utilized mass-spectrometry based readouts have documented that increasing lipophilicity tended to cause increasing accumulation of small-molecule scaffolds, such as sulfoadenosines [65, 101]. Our data point in the opposite direction for cytosolic arrival: across the **P0**–**Sat3** series in *E. coli*, added chain length reduced apparent accumulation. The coumarin series (**Fig. 4c**) shows why these observations need not conflict: in those compounds, total cellular fluorescence rose with chain length while CHAMP-reported cytosolic arrival fell, indicating that added lipophilicity increases envelope association even as it decreases cytosolic delivery. In HeLa cells, our results agree with prior reports: medium-chain lipids (**Sat6**–**Sat8**), cyclic moieties (**Cyc4**, **Cyc5**), and fluorinated alkyl chains (**Het2**) increased cytosolic accumulation [16, 72–73]. The coumarin series makes this distinction concrete: within a single set of compounds, a whole-cell fluorescence readout and a cytosolic-arrival readout ranked the same molecules in opposite orders. Such knowledge is increasingly important as drug candidates grow more lipophilic to engage “undruggable” targets, often violating Lipinski’s rule of five [103]. Our results therefore both corroborate prior findings in mammalian cells and identify where the prevailing lipophilicity metric breaks down in the diderm envelope.

Varying the lipid on a single small parent molecule allowed us to attribute differences in accumulation to the tail rather than to the scaffold, and to ask which physicochemical properties track with accumulation in each system. By incorporating the lipids into a small, polar parent moiety we aimed to corroborate previous results which functionalized lipid moieties to increase membrane permeability of otherwise membrane-impermeant molecules [16, 102]. Given the nature of using small molecules, it is unclear whether the rules defined here will apply to drug scaffolds such as peptide macrocycles, but some evidence points to similarities between the findings here and previous studies [104]. Furthermore, the endogenous modifications made in this study reveal a key limitation of lipidation: its inability to overcome the permeability barriers imposed by charged groups in either system. It is well noted that other factors, such as basicity of amines and H-bond donors, factor significantly into the accumulation of molecules into the cytosol [102]. However, the results here suggest lipidation, and the subsequent increase of hydrophobicity, may not be enough to overcome those modifications to the small molecules across both systems. Perturbing individual envelope layers identified which structural classes are limited by which barrier: outer-membrane permeabilization rescued only the smallest, most polar conjugates, whereas disrupting the proton motive force or inhibiting AcrAB–TolC increased accumulation of medium-chain conjugates. These results implicate the inner membrane and its associated efflux systems, rather than the outer membrane, as the principal constraint on this compound class, though, as noted above, the perturbations used here cannot separate the passive barrier from the active systems that depend on it. Isolating these barriers through systematic lipidation strategies, potentially combined with adjuvants, can provide enhancements towards the accumulation of desired drugs in Gram-negative bacteria. By filling gaps where other approaches fall short, this analysis adds to the toolkit for combating antimicrobial resistance.

Several limitations bound these conclusions. All primary screening data are single-concentration, single-time-point measurements, and while CP₅₀ determinations for a representative subset support the ordering, the full-library comparisons should be read as rank-order rather than as quantitative differences. Because HaloTag expression, cell volume, and assay dynamic range differ between HeLa cells and E. coli, we compare compounds within each system and compare only the resulting trends across systems. The chemical space sampled here is deliberately narrow (a small, polar dipeptide scaffold with high polar surface area and multiple hydrogen-bond donors relative to its size) and the trends we report may not extend to larger or more lipophilic parent scaffolds. For practitioners, the operational conclusion is principally: lipidation should not be treated as a general permeability-enhancing strategy. In mammalian cells, medium-chain, cyclic, and fluorinated tails improved cytosolic delivery of a small polar scaffold; in *E. coli*, only the shortest acyl groups did, and every larger modification was counterproductive. Programs targeting the Gram-negative cytosol are therefore unlikely to benefit from the lipophilicity-driven optimization that works in mammalian systems, and are better served by adjuvant strategies matched to the specific barrier limiting a given scaffold.

## ACKNOWLEDGEMENT

This study was supported by the NIH grant 1R01AI178975-01 (M.M.P.), R35GM124893 (M.M.P.), R01AI179080-01 (M.M.P.).

## SUPPORTING INFORMATION

Additional Figures, tables, and materials/methods are included in the supporting information file

## Notes

### Competing Interest Statement

The authors have declared no competing interest.

### Summary of Updates

We have added additional experiments (in biological triplicates) that enrich and strengthen the text.

