## Supplementary Information for "Structure-Dependent Effects of Lipid Conjugation on Cytosolic Accumulation in *Escherichia coli* and Mammalian Cells"

### Table of Contents


| | |
|---|---|
| **Supplementary Figures** | **S4-S15** |
| **Figure S1.** DBCO-polystyrene bead assay assessing the reactivity of azide-tagged lipid conjugates | **S4** |
| **Figure S2.** ANS assay measurements for the lipid conjugate library | **S5** |
| **Figure S3.** CHAMP assay workflow schematics | **S6** |
| **Figure S4.** One-step labeling assay for detection of IPTG induced HaloTag expression in smooth *E. coli* | **S7** |
| **Figure S5.** Three-step assay with **P0** and **Sat1** visualized with Fl-az and TAMRA-az | **S8** |
| **Figure S6.** Mean fluorescence intensity statistics of the library in smooth *E. coli* | **S19** |
| **Figure S7.** NPN assay of entire library measuring outer membrane permeabilization | **S10** |
| **Figure S8.** SYTOX Green assay measuring complete membrane permeabilization | **S11** |
| **Figure S9.** Mean fluorescence intensity statistics of the library in HeLa cells | **S12** |
| **Figure S10.** DBCO-polystyrene bead assay assessing the reactivity of azide-tagged lipid conjugates in sub-libraries | **S13** |
| **Figure S11.** Mean fluorescence intensity statistics of the sub-libraries in both *E. coli* and HeLa | **S14** |
| **Figure S12.** CHAMP assay with library utilizing hyperporinated rough *E. coli* compared to WT rough *E. coli* | **S15** |
| **Materials and Methods** | **S16-S22** |
| **Materials** | **S16-S17** |
| **Experimental Methods** | **S17-S22** |
| HaloTag protein expression and bacterial cell culture | **S17** |
| One-step bacterial assay with rhodamine chloroalkane (R110cl) | **S17** |
| Two-step bacterial assay with TAMRA-az (before fixation) | **S18** |
| Two-step assay with Fl-az (after fixation) | **S18** |
| Bacterial CHAMP assay protocol (three-step assay) | **S18** |

| | |
|---|---|
| Three-step bacterial assay with PMBN | **S19** |
| Three-step bacterial assay with PAβN | **S19** |
| Three-step bacterial Assay with CCCP | **S19** |
| Mammalian cell culture | **S19** |
| One-step labeling in HeLa | **S19** |
| Two-step labeling in HeLa | **S20** |
| Mammalian CHAMP assay protocol (three-step assay) | **S20** |
| Three-step assay with varying bovine serum albumin (BSA) | **S20** |
| Measuring reactivity of azides | **S21** |
| Outer membrane permeability by NPN uptake | **S21** |
| Inner membrane permeability by SYTOX Green | **S21** |
| Critical aggregation concentration assay determination with ANS | **S21** |
| Calculation of physiochemical properties of the test molecules | **S22** |
| **Synthesis and Characterization of Test Molecules** | **S22-S63** |
| General procedure for solid-phase peptide synthesis of target molecules via solid-phase peptide synthesis | **S22** |
| Solid-phase peptide synthesis of **3d** | **S23** |
| Solid-phase peptide synthesis of **CoumX** series | **S23** |
| **Characterization of Test Molecules** | **S23-S63** |
| General library members | **S23-S53** |
| **Sat7** series | **S54-S56** |
| **Sat3** Series | **S57-S59** |
| **CoumX** series | **S60-S63** |
| **References** | **S65** |

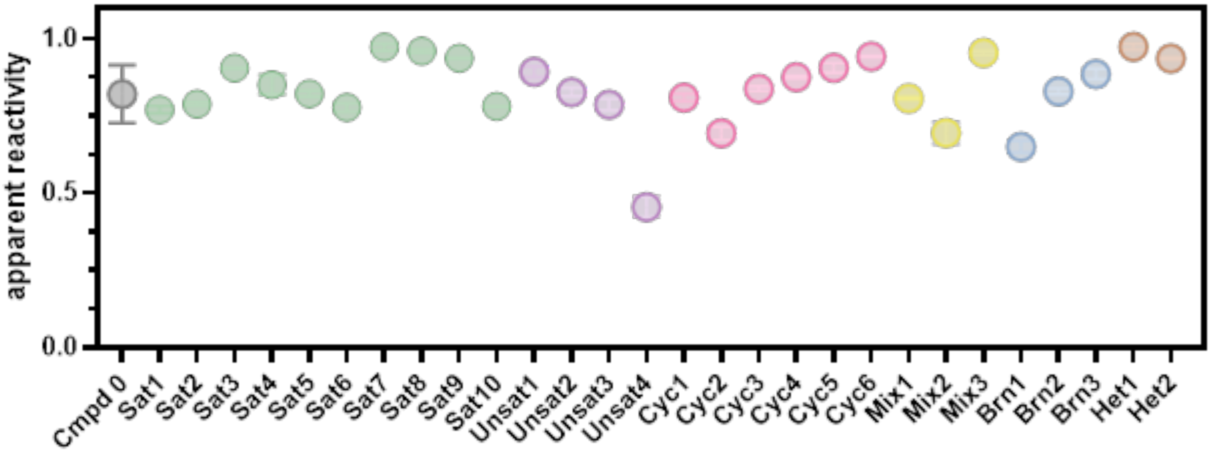


**Figure S1.** Apparent reactivity data of the entire library using the DBCO-modified polystyrene bead assay with the entire library. Data are represented as mean ± SD (n= 3) of technical replicates.

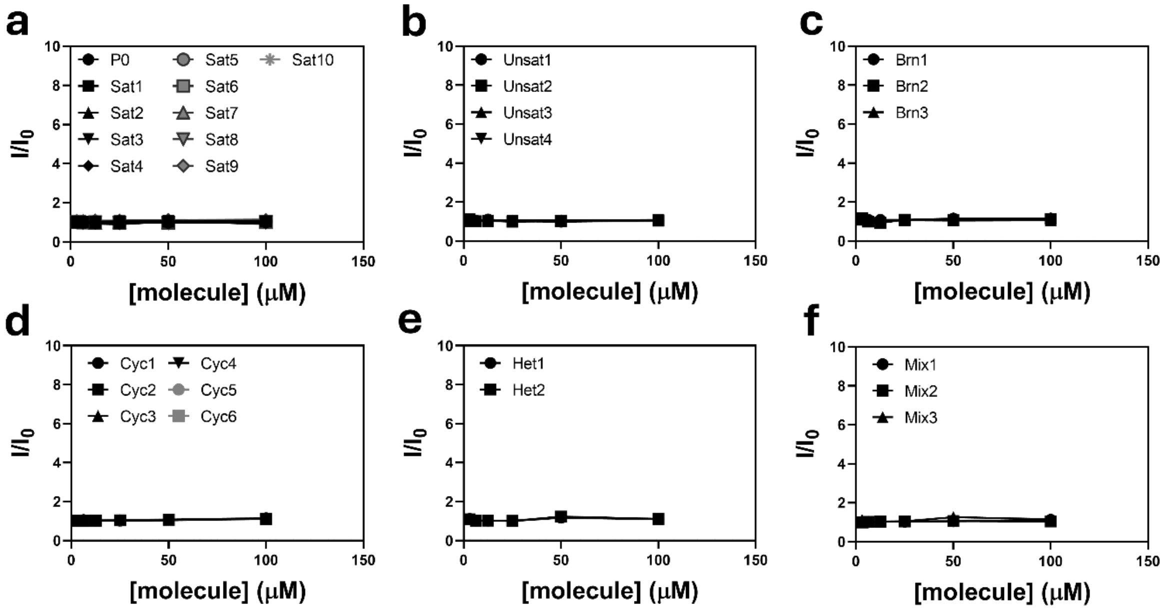


**Figure S2.** Results of the ANS assay for potential aggregation in the different subcategories of lipids: (**a**) saturated chains, (**b**) unsaturated lipids, (**c**) branched chains, (**d**) cyclized lipids, (**e**) heteroatom series and (**f**) mixed modifications. Data are represented as the intensity of ANS signal at 530 nm of treated groups over the untreated group. Data are represented as mean ± SD (n= 3) of technical replicates.

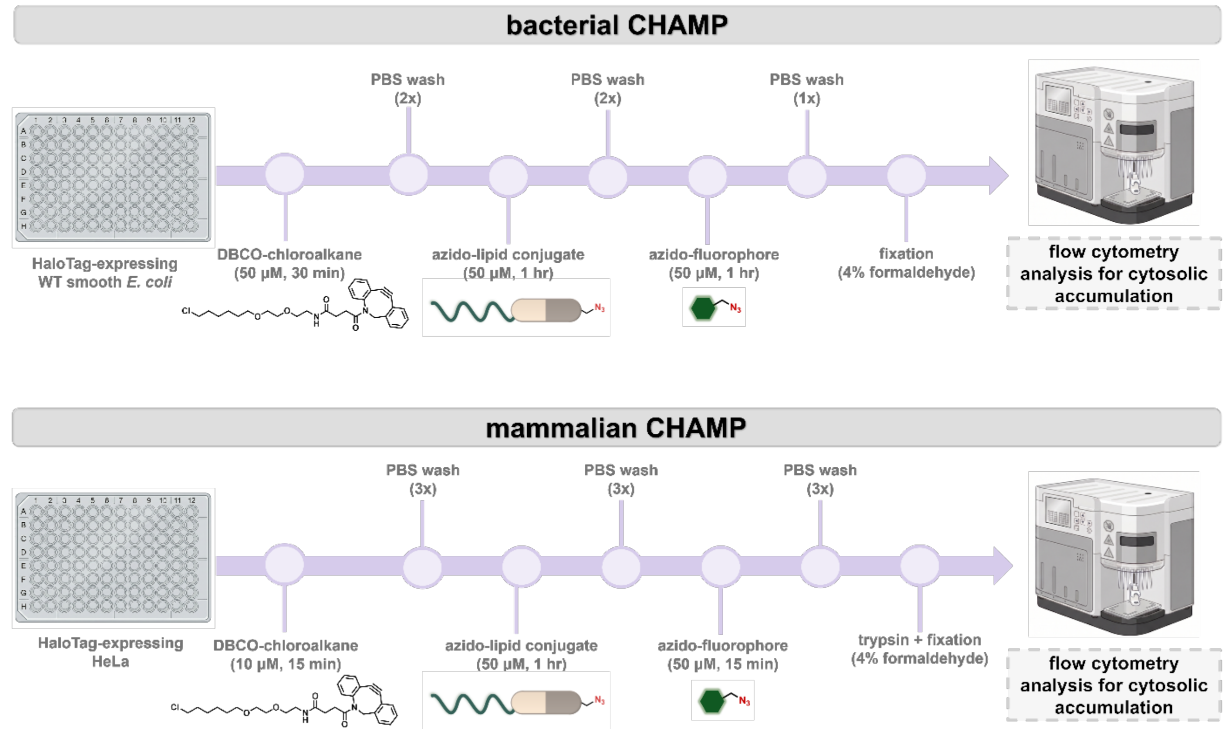


**Figure S3.** Schematic representations of the workflow for CHAMP in both *E. coli* and mammalian cells.

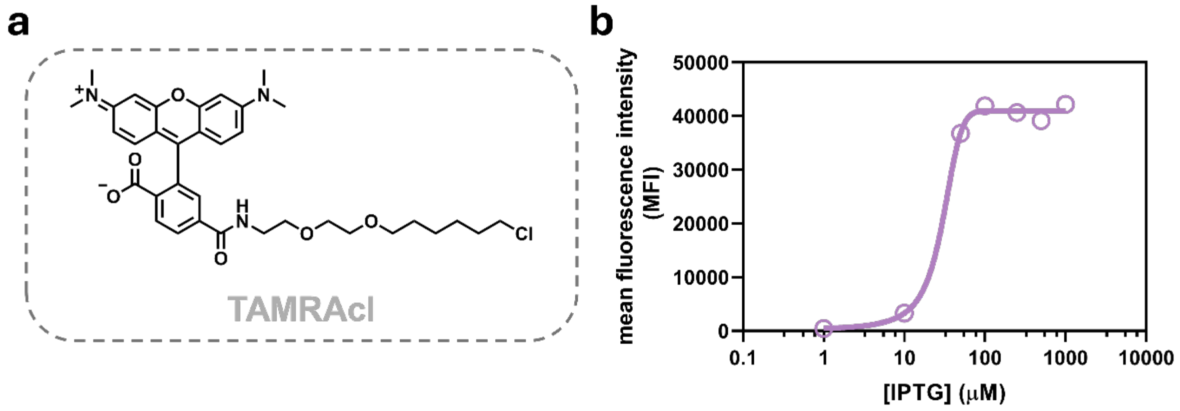


**Figure S4. a,** Structure of **TAMRAcl**, the direct readout for the one step assay. **b** Results of the one-step assay assessing optimization of IPTG concentration for expression of HaloTag protein in smooth *E. coli*. Data are represented as mean ± SD (n= 3) of technical replicates.

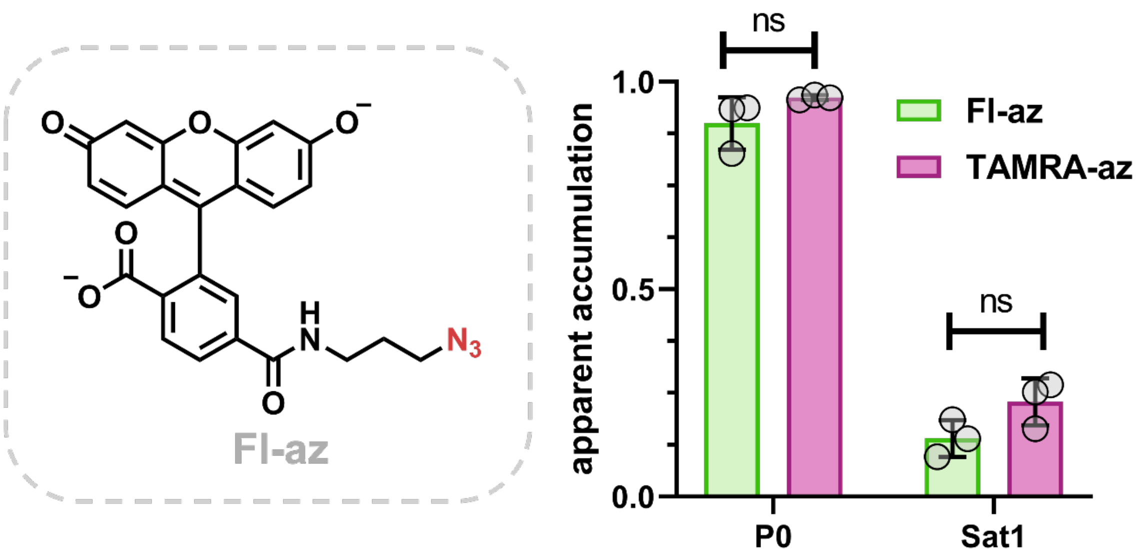


**Figure S5.** Structure of azide modified fluorescein (**Fl-**az) and CHAMP data in smooth *E. coli* using two different readouts **Fl-az** and **TAMRA-az** using **P0** and **Sat1.** Cells visualized with **Fl-az** were fixed with 4% formaldehyde before incubation with the fluorophore, Data are represented as mean ± SD (n= 3) of technical replicates.

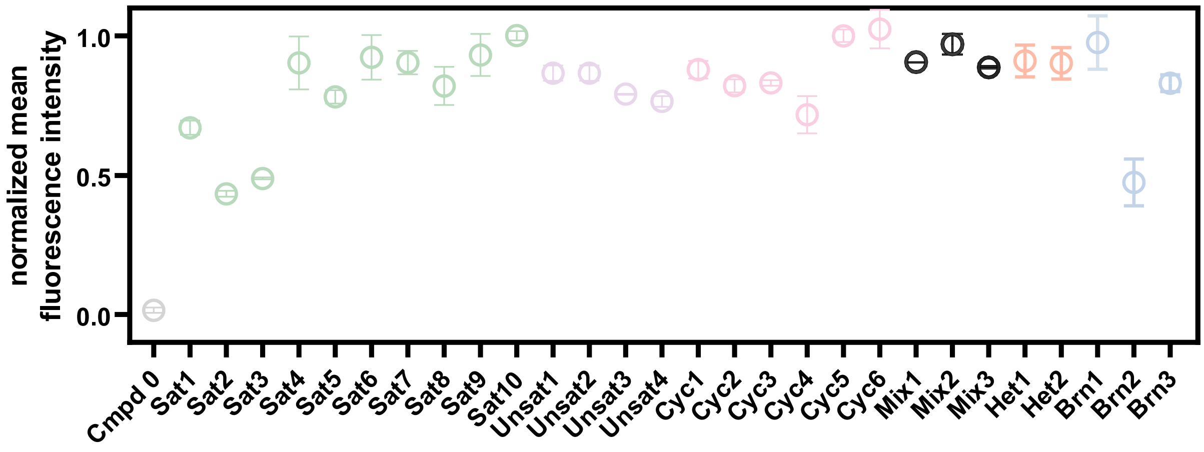


**Figure S6.** Graph showing the normalized mean fluorescence intensity of the entire library in smooth *E. coli* cells. Fluorescence values are normalized to the maximum signal achieved from **DBCOcl**-treated cells with **TAMRA-az** compared to the cells treated with just **TAMRA-az.** Data are represented as mean +/- SD (n = 3) of biological replicates.

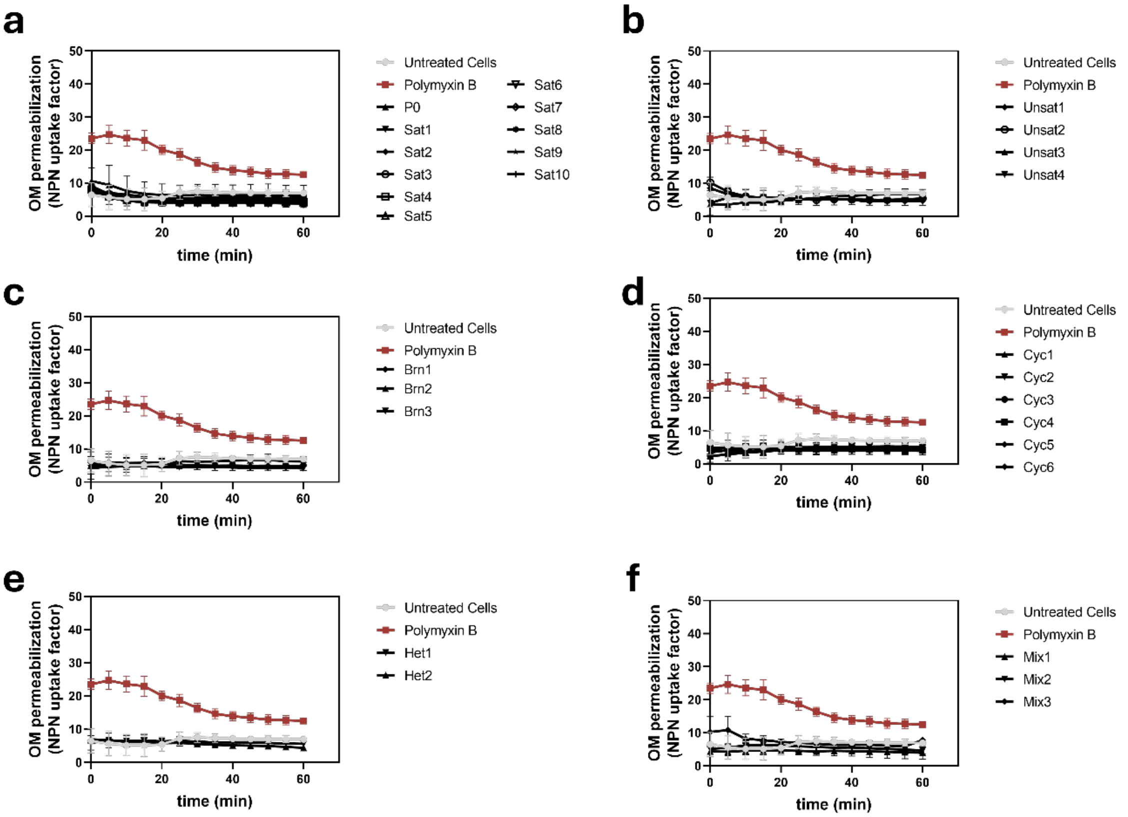


**Figure S7.** Graphics representing the results of NPN coincubation to measure *E. coli* outer membrane integrity: (**a**) saturated chains, (**b**) unsaturated lipids, (**c**) branched chains, (**d**) cyclized lipids, (**e**) heteroatom series and (**f**) mixed modifications. Results show the calculated “NPN uptake factor” over the 1 hr incubation period used for the groups. 16 µg/mL Polymyxin B was used as a positive control for NPN uptake in this experiment. Data are represented as mean +/- SD (n = 3) of technical replicates. Raw NPN fluorescence values were normalized to $OD_{600nm}$.

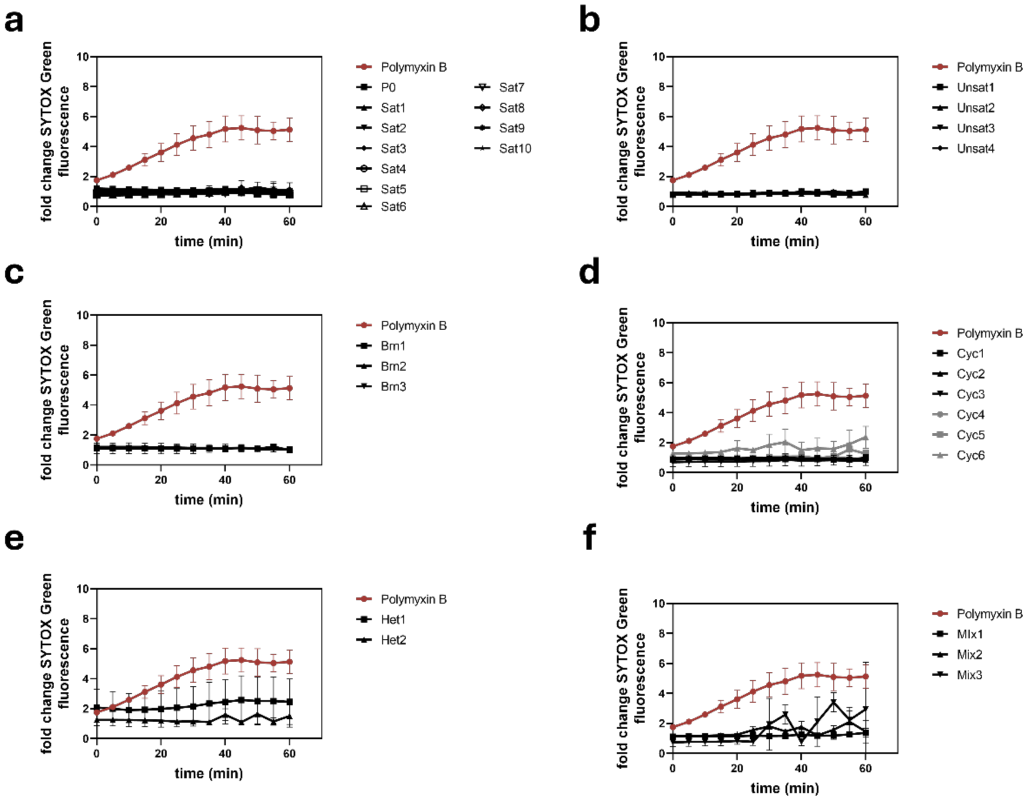


**Figure S8.** Graphics represent the results of SYTOX Green coincubation to measure *E. coli* complete cell envelope integrity: (**a**) saturated chains, (**b**) unsaturated lipids, (**c**) branched chains, (**d**) cyclized lipids, (**e**) heteroatom series and (**f**) mixed modifications. Results show the calculated fold change of SYTOX Green fluorescence compared to the untreated cells over the 1 hr incubation period used for the groups. 32 µg/mL Polymyxin B was used as a positive control for SYTOX Green uptake in this experiment. Data are represented as mean +/- SD (n = 3) of technical replicates. Raw NPN fluorescence values were normalized to $OD_{600nm}$.

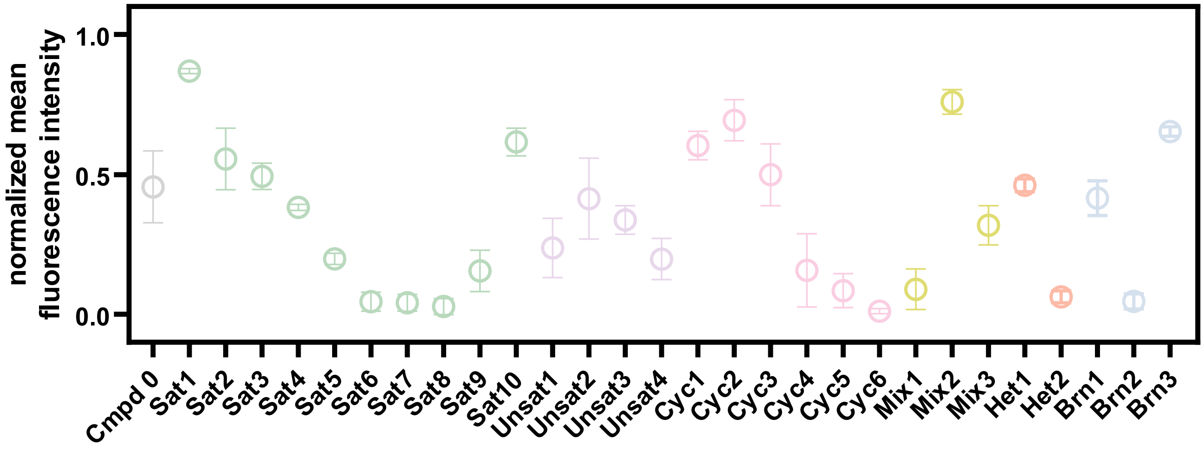


**Figure S9.** Graph showing the normalized mean fluorescence intensity of the entire library in HeLa cells. Fluorescence values are normalized to the maximum signal achieved from **DBCOcl**-treated cells with **TAMRA-az** compared to the cells treated with just **TAMRA-az.** Data are represented as mean +/- SD (n = 3) of biological replicates.

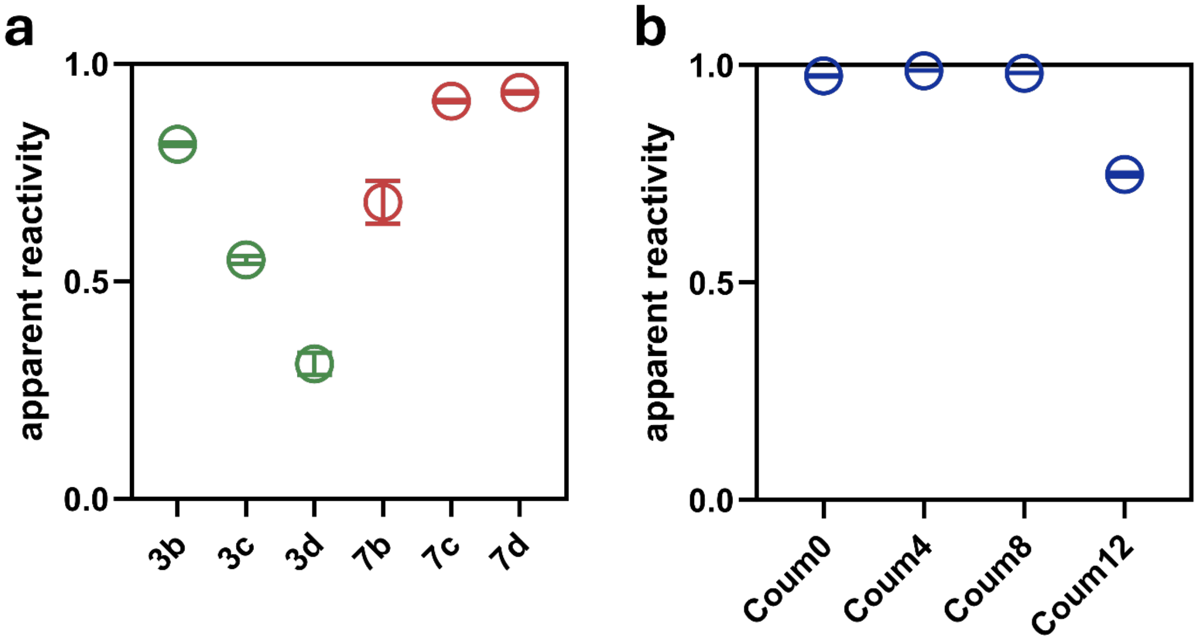


**Figure S10.** Apparent reactivity data of the **a, Sat3** and **Sat7** derivatives, as well as **b, CoumX** derivatives using the DBCO-modified polystyrene bead assay. Data are represented as mean ± SD (n= 3) of technical replicates.

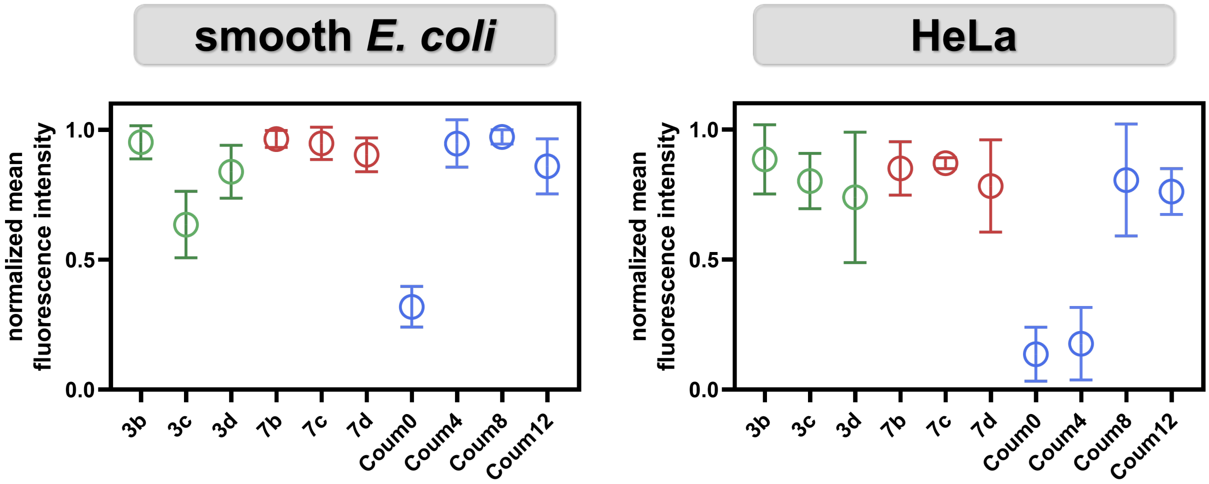


**Figure S11.** Graph showing the normalized mean fluorescence intensity of the library of endogenous changes in both *E. coli* and HeLa cells. Fluorescence values are normalized to the maximum signal achieved from **DBCOcl**-treated cells with **TAMRA-az** compared to the cells treated with just **TAMRA-az.** Data are represented as mean +/- SD (n = 3) of biological replicates.

#### *hyperporinated E. coli*

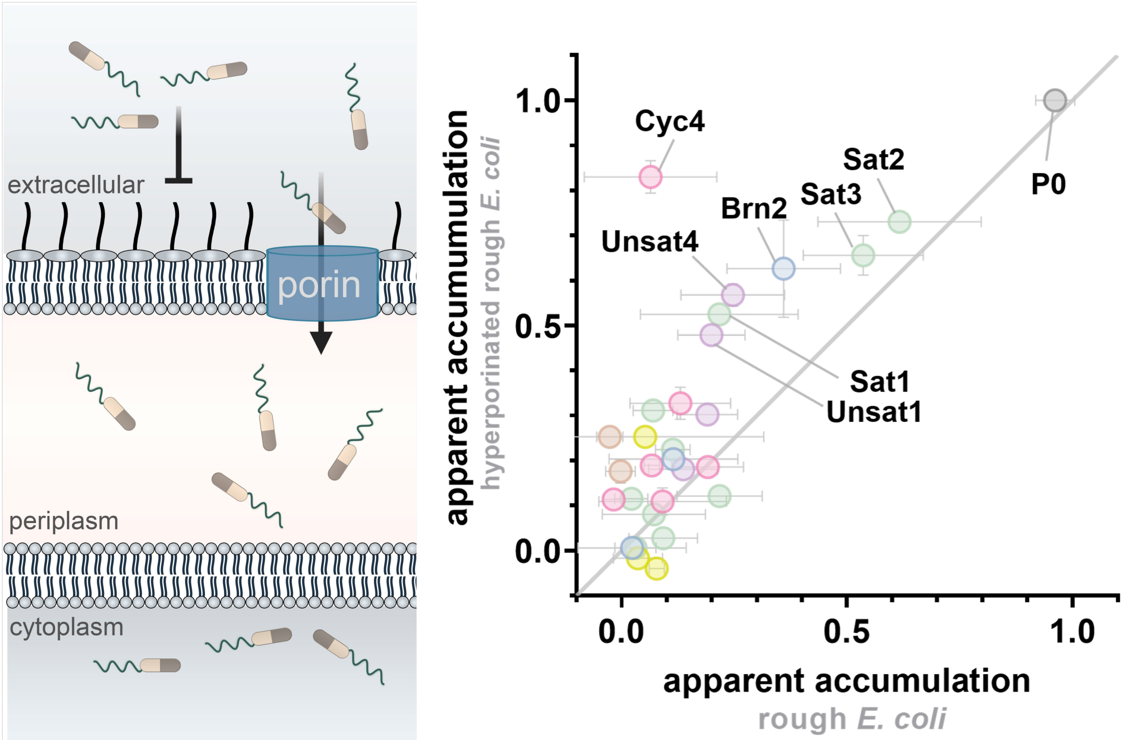


**Figure S12.** Schematic representation of the hyperporinated strain of rough *E. coli* expressing the mutant fhuA membrane protein. CHAMP analysis revealing apparent accumulation differences between the hyperporinated strain (y-axis) and WT rough *E. coli* (x-axis). Data are represented as mean +/- SD (n = 3) of technical replicates.

#### Materials and Methods
#### Materials

| Reagent | Vendor Source | Catalog # |
|---|---|---|
| **Reagents for biological methods** | | |
| DMEM, high glucose, pyrvuate | Sigma-Aldrich | 11995065 |
| Fetal Bovine Serum | Sigma-Aldrich | A52568-01 |
| Penicillin-Streptomycin | Sigma-Aldrich | P4333 |
| Puromycin Dihydrochloride | Sigma-Aldrich | P4512 |
| TrypLETM Express Enzyme (1X), phenol red | Fisher Scientific | 12605028 |
| TAMRA-azide, 6-isomer | Lumiprobe | C8130 |
| N-phenyl-napthylamine (NPN) | Sigma-Aldrich | 104043 |
| SYTOX Green | Fisher Sceintific | S7020 |
| 8-Anilinonaphthalene-1-sulfonic acid (ANS) | Ambeed | A666405 |
| Formaldehyde solution | Sigma-Aldrich | 8187081000 |
| Fluorescein azide, 6-isomer | Lumiprobe | D5130 |
| Phosphate buffered saline | Sigma-Aldrich | P3813 |
| Bovine Serum Albumin (BSA) | Sigma-Aldrich | A9418 |
| LB broth, miller | Sigma-Aldrich | L3152-1K |
| Polymyxin B nonapeptide | Sigma-Aldrich | P2076 |
| Phenylalanine Arginine Beta-Naphthylamide (PAβN) dihydrochloride | Sigma-Aldrich | P4157 |
| Carbonyl cyanide *m*-chlorophenylhydrazone (CCCP) | Ambeed | A806566 |
| Isopropyl-β-D-thiogalactopyranoside | ChemImpex | 00194 |
| Polymyxin B Sulfate | AK Scientific | J90069 |
| Ampicillin Sodium Salt | Sigma-Aldrich | A9518 |
| **Reagents for the synthesis and characterization of lipid-conjugate library** | | |
| Rink amide resin (200-400 mesh, 0.3-0.8 meq/g) | ChemImpex | 10619 |
| 2-Chlorotrityl chloride resin (100-200 mesh, 1.0-2.0 meq/g) | ChemImpex | 03498 |
| α-N-Fmoc-amino acids | Ambeed | Various |
| Carboxylic Acid Lipids | Ambeed, ChemImpex, Fisher Scientific | Various |
| 7-(Diethylamino)coumarin-3-carboxylic acid | BroadPharm | BP-41783 |
| N,N'-Diisopropylcarbodiimide (DIC) | ChemImpex | 00100 |

| | | |
|---|---|---|
| Ethyl Cyano(hydroxyamino)acetate (Oxyma) | TCI Chemicals | E0847 |
| N,N-dimethylformamide (DMF) | Sigma-Aldrich | 319937 |
| Dichloromethane (DCM) ACS-grade | Sigma-Aldrich | D65100 |
| Methanol ACS-grade | Sigma-Aldrich | 179337 |
| Methanol HPLC-grade | Sigma-Aldrich | 34860 |
| O-(Benzotriazol-1-yl)-N,N,N',N'-tetramethyluronium hexafluorophosphate (HBTU) | ChemImpex | 02011 |
| Piperidine | ChemImpex | 02351 |
| N,N'-diisopropylethylamine (DIEA) | ChemImpex | 00141 |
| Trifluoroacetic acid (TFA) reagent-grade | ChemImpex | 00289 |
| Trifluoroacetic acid (TFA)HPLC-grade | Sigma-Aldrich | 302031 |
| Triisopropylsilane (TIPS) | ChemImpex | 01966 |
| **General materials and equipment** | | |
| Greiner Bio-One CELLSTAR TC Treated Cell Culture Flasks | Fisher Scientific | 07-000-225 |
| Corning™ Costar™ 96-Well, Cell Culture Treated, Flat-Bottom Microplate | Fisher Scientific | 09-761-145 |
| 96-well clear untreated round bottom plates | VWR | 82050-622 |

#### Experimental Procedures

##### HaloTag protein expression and bacterial cell culture

For expression of the HaloTag protein in WT *E. coli*, (smooth, ATCC25922) and in different variants of *E. coli* K12 (rough, BW25113), the expression plasmid Halo_pQE-30 was transformed into *Escherichia coli* and grown on an LB/agar plate with ampicillin (100 µg/mL) at 37 °C overnight. Colonies were picked and grown first in 4 mL LB broth with ampicillin at 37 °C overnight. A growth tube with 4mL of LB media was then inoculated with the overnight culture at a ratio of 1:100 in the presence of 100 µg/mL ampicillin and cells were grown at 37 °C for 1.5 hours, or until the optical density at 600 nm reached 0.2 when cultures were induced with 1 mM Isopropyl β-d-1-thiogalactopyranoside (IPTG) at 37 °C for 2 h in a shaker incubator. Cells were then pelleted at 4000 rpm for 3 min. The pellet was washed three times with 1X PBS pH 7.4, following which cells were resuspended in 35 mL of 1X PBS for the assay.

##### One-step bacterial assay with rhodamine chloroalkane (R110cl)

HaloTag expressing *E. coli* cells were washed 2X with 4 mL of 1x PBS and finally re-suspended in 4 mL of 1x PBS. The cells were pipetted into wells in a 96-well plate in triplicate, pelleted down by centrifuging for 3 min at 2700 x *g*, and resuspended using 50 µM R110cl. The cells were incubated for 30 min at 37°C, centrifuged for 3 min, washed 1X with PBS, and fixed with 4% formaldehyde in PBS for 30 min at 37°C . The cells were subjected to analysis by flow cytometry on the Attune NxT Acoustic Focusing Cytometer (Invitrogen).

**Two-step bacteria assay with TAMRA-az (before fixation)**

DBCO-labeled *E. coli* cells were prepared as described earlier and 100 µL of the cell culture was added into each well of a 96-well plate to be washed 2x with PBS. After washing, cells were resuspended in 50 µL of PBS and 50 µL of a 100 µM TAMRA-az solution in PBS was added to the wells. The plates were covered with plastic film, and incubated for 1 h at 37 °C in a shaker incubator. Each azido-tagged molecule or dye was replicated in three wells. The plate was then spun at 4000 rpm for 3 min using a plate holder centrifuge, the supernatant was discarded, and the cells were washed 1X with PBS. Next, cells were fixed with 4% formaldehyde in PBS for 30 min at 37 °C in a shaker incubator. Following formaldehyde fixation, the cells were washed 1X with PBS and resuspended in 200 µL of PBS for analysis by flow cytometry.

**Two-step assay with Fl-az (after fixation)**

DBCO-labeled *E. coli* cells were prepared as described earlier and 100 µL of the cell culture was added into each well of a 96-well plate to be washed 2x with PBS. After washing, cells were fixed with 4% formaldehyde in PBS for 30 min at 37 °C in a shaker incubator. The cells were washed 2x with PBS and cells were resuspended in 50 µL of PBS and 50 µL of a 100 µM Fl-az solution in PBS was added to the wells. The plates were covered with plastic film and incubated for 1 h at 37 °C in a shaker incubator. Each azido-tagged molecule or dye was replicated in three wells. The plate was then spun at 4000 rpm for 3 min using a plate holder centrifuge, the supernatant was discarded, and the cells were washed 1X with PBS. Following the wash with PBS and resuspended in 200 µL of PBS for analysis by flow cytometry on the Attune NxT Acoustic Focusing Cytometer (Invitrogen).

**Bacterial CHAMP Assay Protocol (Three-Step Assay)**

Following IPTG-expression, 4 mL of 50 µM **DBCOcl** would be added to the *E. coli* cells and were incubated for 30 min at 37 °C. 100 µL of the cell suspension was plated in triplicate on a 96 well plates and subsequently washed 2x with PBS. Then, cells were resuspended in 50 µL of PBS and 50 µL of a 100 µM molecule stock, obtained from a 10 mM parent stock, was added to each of the wells, which was then covered and incubated for 1 hr at 37 °C. In the case of time and concentration studies, similar protocol was altered to account for these changes. Control wells, on the same plate, were incubated with 100 µL of PBS for the same duration. Cells were then pelleted in a plate holder centrifuge at 4000 RPM for 3 min and the supernatant was discarded. After washing 2x with PBS, the cells were immediately resuspended in 50 µL of PBS and to that was added 50 µL of 100 µM TAMRA-az, 6-isomer and incubated for 1 hr at 37 °C. Cells were pelleted once again, washed 1x with PBS, and fixed with 4% formaldehyde for 30 min. Following formaldehyde fixation, the cells were washed 1X with PBS and resuspended in 200 µL of PBS for analysis by flow cytometry on the Attune NxT Acoustic Focusing Cytometer (Invitrogen).[1]

#### Three-step bacterial assay with PMBN

DBCO-labeled smooth *E. coli* was prepared as stated earlier and washed 2x with PBS. Cells were resuspended in 50 µL of a 100 µM molecule stock and immediately 50 µL PBS supplemented with 50 µg/mL PMBN and was added to reach a final concentration of 25 µg/mL PMBN. Cells were incubated for 1 hr at 37 °C in the shaker incubator. Cells were then pelleted in a plate holder centrifuge at 4000 RPM for 3 min and the supernatant was discarded. After washing 2x with PBS, the cells were immediately resuspended in 50 µL of PBS and to that was added 50 µL of 100 µM TAMRA-az, 6-isomer and incubated for 1 hr at 37 °C. Cells were pelleted once again, washed 1x with PBS, and fixed with 4% formaldehyde for 30 min. Following formaldehyde fixation, the cells were washed 1X with PBS and resuspended in 200 µL of PBS for analysis by flow cytometry on the Attune NxT Acoustic Focusing Cytometer (Invitrogen).

#### Three-step bacterial assay with PAβN

DBCO-labeled smooth *E. coli* was prepared as stated earlier and washed 2x with PBS. Cells were resuspended in 50 µL of a 100 µM molecule stock and immediately 50 µL PBS supplemented with 32 µg/mL PAβN and was added to reach a final concentration of 16 µg/mL PAβN. Cells were incubated for 1 hr at 37 °C in the shaker incubator. Cells were then pelleted in a plate holder centrifuge at 4000 RPM for 3 min and the supernatant was discarded. After washing 2x with PBS, the cells were immediately resuspended in 50 µL of PBS and to that was added 50 µL of 100 µM TAMRA-az, 6-isomer and incubated for 1 hr at 37 °C. Cells were pelleted once again, washed 1x with PBS, and fixed with 4% formaldehyde for 30 min. Following formaldehyde fixation, the cells were washed 1X with PBS and resuspended in 200 µL of PBS for analysis by flow cytometry on the Attune NxT Acoustic Focusing Cytometer (Invitrogen).

#### Three-step bacterial assay with CCCP

DBCO-labeled smooth *E. coli* was prepared as stated earlier and washed 2x with PBS. Cells were resuspended in 100 µL of a 100 µM CCCP and incubated for 10 min at 37 °C in the shaker incubator. Cells were washed 1x with PBS and resuspended in 50 µL of PBS before addition of 50 µL of a 100 µM stock of each molecule. Cells were incubated for 1 hr at 37 °C in the shaker incubator. Cells were then pelleted in a plate holder centrifuge at 4000 RPM for 3 min and the supernatant was discarded. After washing 2x with PBS, the cells were immediately resuspended in 50 µL of PBS and to that was added 50 µL of 100 µM TAMRA-az, 6-isomer and incubated for 1 hr at 37 °C. Cells were pelleted once again, washed 1x with PBS, and fixed with 4% formaldehyde for 30 min. Following formaldehyde fixation, the cells were washed 1X with PBS and resuspended in 200 µL of PBS for analysis by flow cytometry on the Attune NxT Acoustic Focusing Cytometer (Invitrogen).

#### Mammalian cell culture

HaloTag-expressing HeLa (HT HeLa) cells were cultured in Dulbecco's Modified Eagle Medium (DMEM) supplemented with 10% fetal bovine serum (FBS), 1% penicillin-streptomycin, and 1 µg/mL puromycin. Cells were maintained at 37°C with 5% $CO_2$. HaloTag HeLa cells were passaged using trypsin upon reaching 80–90% confluency and resuspended in DMEM. Cells were seeded in 96-well plates at a density of 3 x $10^4$ cells per well, incubated for 24 hours at 37°C with 5% $CO_2$.

**One-step labeling in HeLa**

3 x1 $0^4$ HT HeLa cells were seeded into 96-well plates one day before the assay, reaching 80–90% confluency. Following overnight incubation at 37°C, cells were washed three times with PBS and treated with 50 µM TAMRAcl or its “dead” control (lacking a terminal chloro group) in media for 15 minutes. Then, the cells were washed three times using PBS, trypsinized for 5 min, and fixed with 4% formaldehyde in PBS for 30 min statically at room temperature. The cells were then subjected to analysis by flow cytometry on the Attune NxT Acoustic Focusing Cytometer (Invitrogen).

**Two-step labeling in HeLa**

3 x $10^4$ HT HeLa cells were seeded into 96-well plates one day before the assay, reaching 80–90% confluency. Following overnight incubation at 37°C, cells were washed three times with PBS and treated with 100 µL of10 µM DBCOcl in media for 15 minutes which is followed by chase with 100 µL of 50 µM of a TAMRA-az in media for 15 minutes. Then, the cells were washed three times using PBS, trypsinized for 5 min, and fixed with 4% formaldehyde in PBS for 30 min statically at room temperature. The cells were then subjected to analysis by flow cytometry on the Attune NxT Acoustic Focusing Cytometer (Invitrogen).

**Mammalian CHAMP assay protocol (three-step assay)**

3 x $10^4$ HeLa cells were seeded into 96-well plates one day before the assay, reaching 80–90% confluency. Following overnight incubation at 37°C, cells were washed three times with PBS and treated with 10 µM DBCOcl in media for 15 minutes. This was followed by a pulse with 100 µM of azide-tagged compounds for 1 hr at 37°C. Concentration and time of incubation were adjusted based on the experiment requirements. Following the pulse step, a chase step with 50 µM TAMRA-az in media for 15 minutes. Cells were washed three times with PBS between each step. After staining, cells were washed again, trypsinized with TrypLETM Express Enzyme, fixed with 4% formaldehyde, and analyzed using flow cytometry on the Attune NxT Acoustic Focusing Cytometer (Invitrogen).[2]

**Three-step assay with varying bovine serum albumin (BSA)**

80–90% confluency. Following overnight incubation at 37°C, cells were washed three times with PBS and treated with 10 µM DBCOcl in the normal media described above for 15 minutes. This was followed by a pulse with 100 µM of azide-tagged compounds in serum-free DMEM media, supplemented with either 0, 5, or 10 mg/mL BSA for 1 hr at 37°C. Following the pulse step, a chase step with 50 µM TAMRA-az in media for 15 minutes. Cells were washed three times with PBS between each step. After staining, cells were washed again, trypsinized with TrypLE$^{TM}$ Express Enzyme, fixed with 4% formaldehyde, and analyzed using flow cytometry on the Attune NxT Acoustic Focusing Cytometer (Invitrogen).

**Measuring reactivity of azides**

100 µL of Spherotech amino polystyrene beads were placed in a 1.5 mL Eppendorf tube and centrifuged at 21000 xg for 10 mins. Wash with DI water (21000 x g, 10 mins) and resuspend in pH 9 sodium borate buffer. Add 4 µg/mL DBCO-NHS/BCN-NHS or 20 mM propargyl-peg1-NHS and incubate at 37°C with shaking for 2 hours. Wash with DI water. Resuspend pH 9 sodium borate buffer and add 20 µL acetic anhydride/mL DI water. Incubate at 37°C while shaking for 2 hours. Wash twice with DI water and resuspend in 1 mL PBS. Dilute 40x in PBS before using reactivity assays. For acetylated beads, follow the same procedure but skip the NHS labeling step. Increase the volume of acetic anhydride to 60 µL/mL. Take a 0.65 nm filter plate and wet the membrane with PBS. Remove the PBS with vacuum manifold. Add 100 µL of diluted beads to each well of the plate. Remove PBS with vacuum manifold. Add 100 µL of azide solution in PBS to alkyne beads at a concentration of 50 µM. Control wells receive 100 µL of PBS. Cover the plate and incubate for 1 hour at 37°C with shaking. Wash the plate with 200 µL of PBS using the vacuum manifold. Add 100 µL of 50 µM FAM-az in PBS to each well of the plate. Cover the plate and incubate at 37°C with shaking for 1 hour. Wash the plate with 200 µL of PBS using a vacuum manifold. Resuspend the beads in 200 µL of PBS, pipetting up and down, and transfer to a round-bottomed 96 well plate. Read on the attune flow cytometer.

**Outer membrane permeability by NPN uptake**

For outer membrane permeability, we tested using an NPN uptake assay.[2] 80 µL of NPN (Millipore-Sigma), at a final concentration of 10 µM in PBS, was added to the DBCO-labeled *E. coli* cells at stationary phase. 20 µL of given molecules were added to achieve a final concentration of 50 µM for a total volume of 100 µL per well. For positive control, 20 µL of Polymyxin B, to achieve a final concentration of 16 µg/mL was added to the cells. All samples were completed in triplicate. Fluorescence intensity, at 37 °C with shaking, was measured using an Agilent Biotek Synergy H1 well-plate reader using an excitation wavelength of 355 nm and an emission wavelength of 405 nm. Measurements were taken every 5 minute for 60 min total, and OM permeabilization was calculated using the NPN uptake factor formula below:

$$\frac{\text{Fluorescence of sample with NPN} - \text{Fluorescence of Sample without NPN}}{\text{Fluorescence of PBS with NPN} - \text{Fluorescence of PBS without NPN}}$$

Raw NPN fluorescence values were normalized using OD600nm.

**Inner membrane permeability by SYTOX Green**

To measure full cell envelope permeabilization, we used the SYTOX green assay.[2] 80 µL of SYTOX green (Invitrogen) in PBS, at a final concentration to 1 µM, was added to DBCO-labeled *E. coli* cells at stationary phase. 20 µL of each peptide, to achieve a final concentration of 50 μM, was added to wells in triplicate. Similar to the NPN assay, 32 μg/ mL polymyxin B was used in the same concentration here as a positive control. Fluorescence was measured in an Agilent Biotek Synergy H1 well-plate reader at an excitation of 504 nm and an emission at 525 nm every 5 min for 60 min at 37 °C with shaking. Fluorescence was normalized by diving each treated sample from the background fluorescence of SYTOX Green. Raw SYTOX Green fluorescence values were normalized with $OD_{600nm}$.

##### Critical aggregation concentration assay determination with ANS

Aggregation concentration of peptides was conducted using the established ANS assay.[3] Here, 250μM 8-anilo-1-naphthalenesulfonic acid (ANS) in PBS was used to dissolve the peptides in a given range of concentrations. Fluorescence spectra was recorded on an Agilent Biotek Synergy H1 well-plate reader with an excitation wavelength of 356 nm and emission of 530 nm. Plates were incubated for 1 hr after the first read at at 37 °C with shaking, where they were read at the endpoint. Results were plotted as $I/I_0$ where I is the maximum fluorescence intensity of ANS containing the peptide samples and $I_0$ being the maximum intensity of signal without any peptide.

##### Calculation of physiochemical properties of the test molecules

DataWarrior software was used to calculate the molecular weight (MW), water/octanol partitioning coefficient (cLogP), aqueous solubility (cLogS) polar surface area (PSA), and number of rotatable bonds for each molecule along with various other values.[4]

#### Synthesis and Characterization of Test Molecules

##### General procedure for solid-phase peptide synthesis of target molecules via solid-phase peptide synthesis

Molecules in the library were prepared by Fmoc-based solid-phase peptide synthesis using rink amide resin (0.5 mmol/g of loading capacity). Amino acids were coupled using four equivalents of amino acid, ethyl (*2Z*)-2-cyano-2-(hydroxyimino) acetate (oxyma), and *N,N'*-diisopropylcarbodiimide in 10 mL of DMF. Deprotection was conducted using a 20% piperidine solution in DMF. For lipid conjugation to the N-terminus, both carboxylic acid-based tails were used. In brief, four equivalents of lipid, ethyl (*2Z*)-2-cyano-2-(hydroxyimino) acetate (oxyma), and *N,N'*-diisopropylcarbodiimide were dissolved 10 mL of DMF and stirred for 2 hours at room temperature. Acetylation with acetic acid was accomplished using a mixture of acetic anhydride, DIEA, and DMF (5:8.5:86.5, v/v/v) for 1 hour at room temperature. All peptides were cleaved from the rink-amide resin using trifluoroacetic acid (TFA), triisopropylsilane (TIPS), and ,1,4-dimethoxybenzene (DMB) (92.5:2.5:5,v/v/v) shaking at room temperature for 2 hours. The solution was filtered and concentrated before precipitation via cold diethyl ether. The resultant peptide was purified using reverse-phased high-performance liquid chromatography (RP-HPLC) equipped with Waters 1525 with 2489 UV/Visible Detector on a Phenomenex Luna 10 μm C8(2) 100 Å (250 x 21.2 mm) column using gradient elution with either $H_2O$/MeCN or

$H_2O$/MeOH with 0.1% TFA. HPLC fractions were concentrated *in vacuo* and lyophilized to dryness using a Labconco Freezone 4.5L (-84°C) lyophilizer. Peptides were further analyzed for purity using a Phenomenex Luna 5 μm C8(2) on the same RP-HPLC; gradient elution in $H_2O$/MeCN with 0.01% TFA at 1 mL/min. The peptide identities were confirmed using electrospray ionization mass spectroscopy (Advion CMS-SO1 ESI Mass Spectrometer) mass spectroscopy (Shimadzu 8020). Peptides were characterized using UV-Vis absorbance at 284.5 (log ε= 3.23) and analyses were obtained on an Agilent 6545B Q-TOF LC/MS equipped with 1260 infinity II LC system with auto sampler.[5]

**Solid-phase peptide synthesis of 3d**

To prepare **3d** with the carboxylic acid C-terminus, 2-chlorotritylchloride-resin (1.55 mmol/g loading capacity) was employed. Synthesis was performed in accordance with standardized protocols for the resin and similar N-α-Fmoc amino acids were used in the synthesis of **3d. 3d** was characterized using UV-Vis absorbance at 284.5 (log ε= 3.23) [ref] and analyses were obtained on an Agilent 6545B Q-TOF LC/MS equipped with 1260 infinity II LC system with auto sampler.

**Solid-phase peptide synthesis of CoumX:**

Synthesis of the CoumX series followed the same protocol as the general procedure for synthesis of the peptides on rink amide resin (0.5 mmol/g loading capacity). However, the Fmoc-Phe(4-NHBoc)-OH was replaced by an methoxytrityl protective group on Fmoc-Dab(Mtt)-OH. Synthesis involved building the backbone and coupling the appropriate lipid to the N-term before a selective sidechain deprotection of the Mtt group. In brief, Mtt removal was performed based on standardized protocols using 1% TFA, 0.5% TIPS, in DCM. The resin was shaken with the DCM-TFA-TIPS mixture for 15 min increments and washed rigorously with DCM in between each shaking step. After 3 increments, 5 equivalents of 7-(Diethylamino)coumarin-3-carboxylic acid in DMF with 10 equivalents of DIPEA and 5 equivalents of HBTU were added to the resin, covered with aluminum foil, and shaken overnight at room temperature. The CoumX molecules were washed, cleaved from the resin, and purified using the HPLC methods labeled above. Peptides were characterized using UV-Vis absorbance at 409 nm (ε= 34,000) [ref] and analyses were obtained on an Agilent 6545B Q-TOF LC/MS equipped with 1260 infinity II LC system with auto sampler.

**P0**

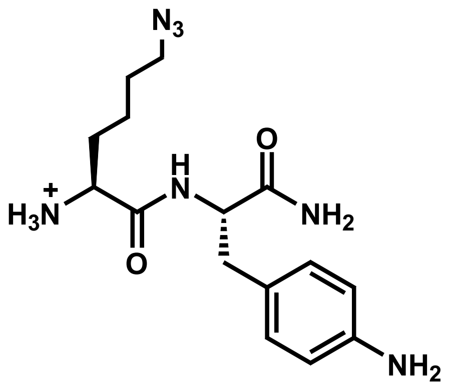


Analytical HPLC Chromatogram of **P0**

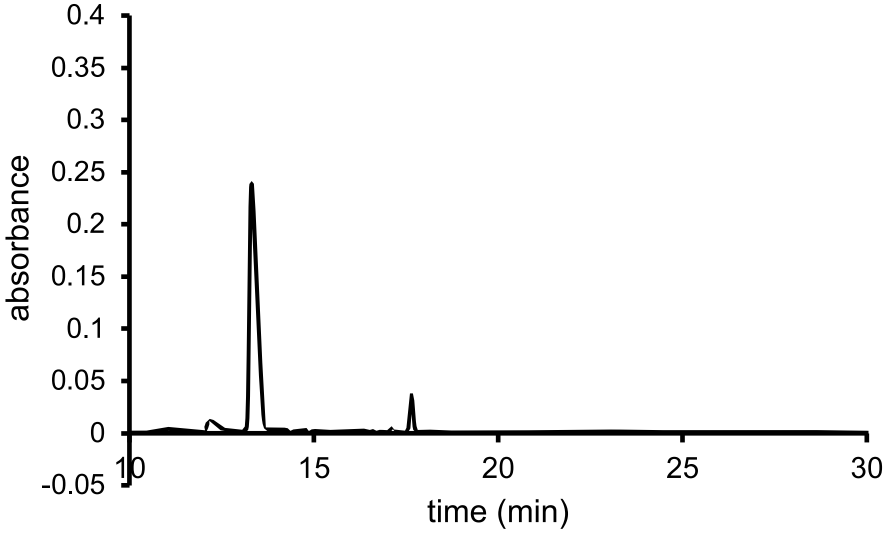


QTOF High Resolution Mass Spectrum for **P0** (m/z 334.1986 for $[M+H]^+$, found 334.2007)

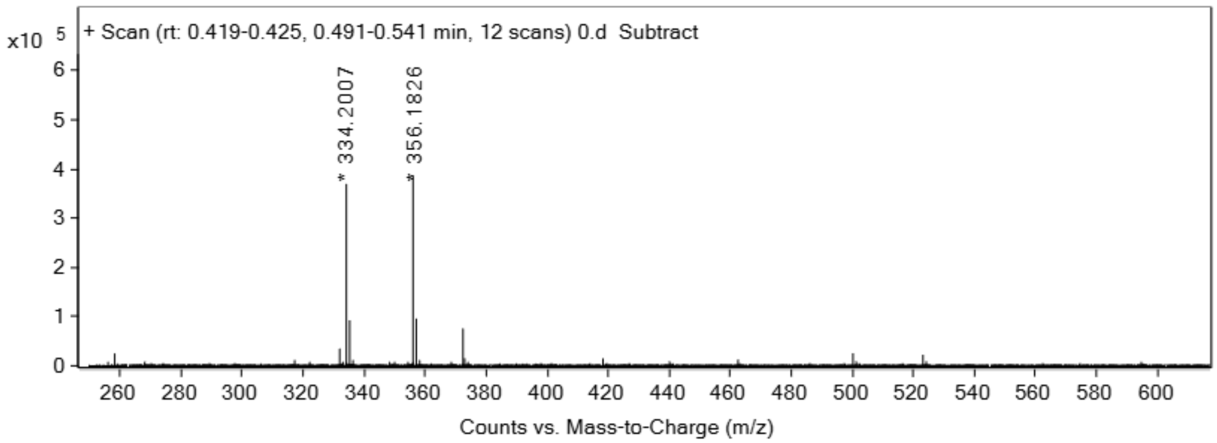

**Sat1**

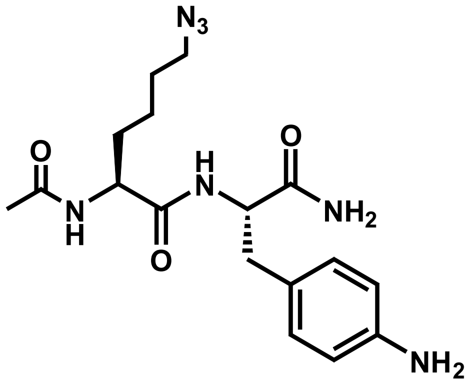


Analytical HPLC Chromatogram of **Sat1**

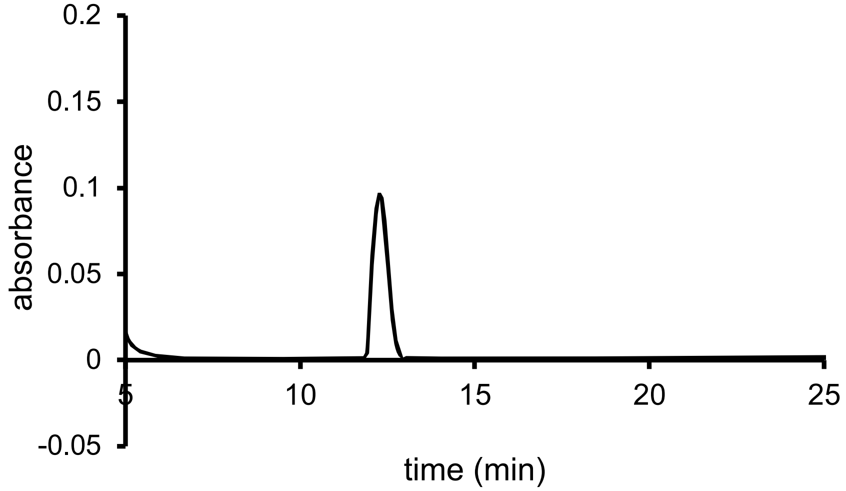


QTOF High Resolution Mass Spectrum for **Sat1** (m/z 376.2092 for $[M+H]^+$, found 376.2096)

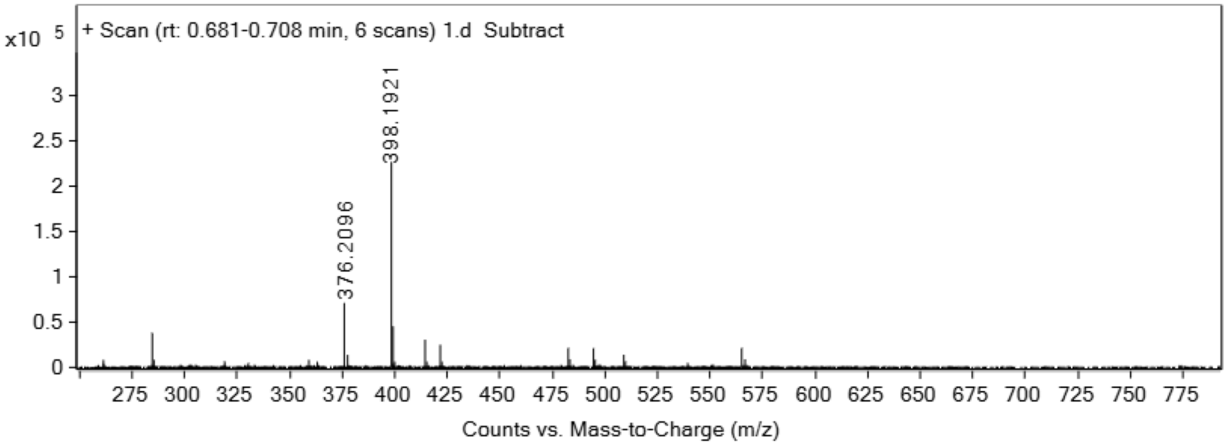

**Sat2**

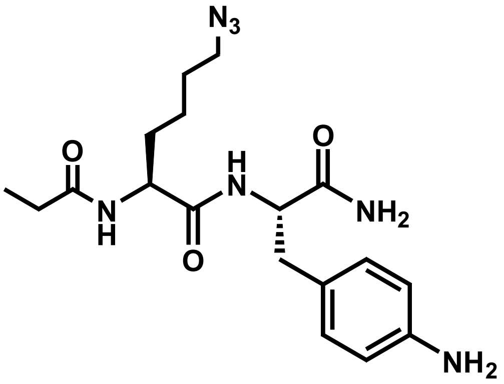


Analytical HPLC Chromatogram of **Sat2**

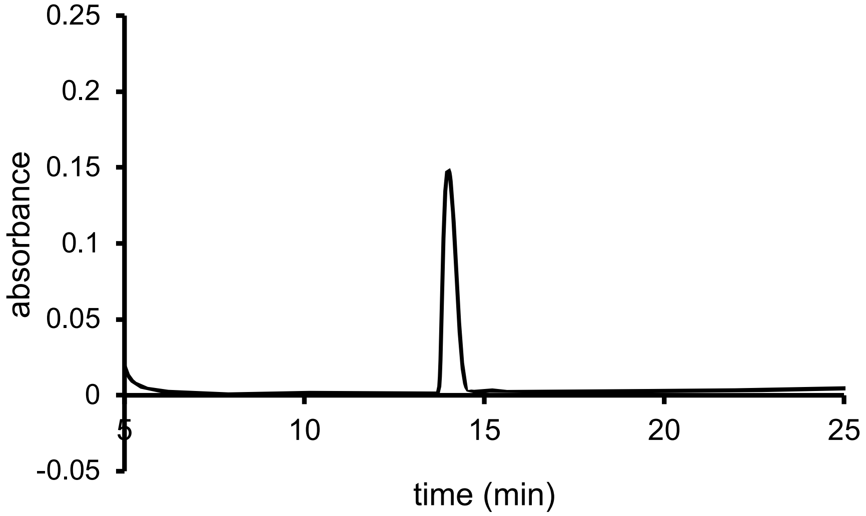


QTOF High Resolution Mass Spectrum for **Sat2** (m/z 390.2249 for $[M+H]^+$, found 390.2251)

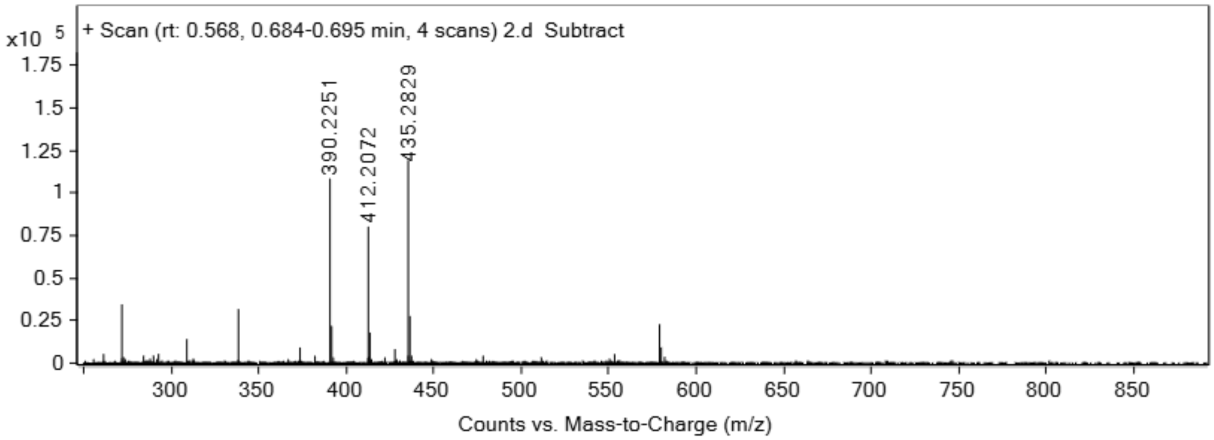

**Sat3**

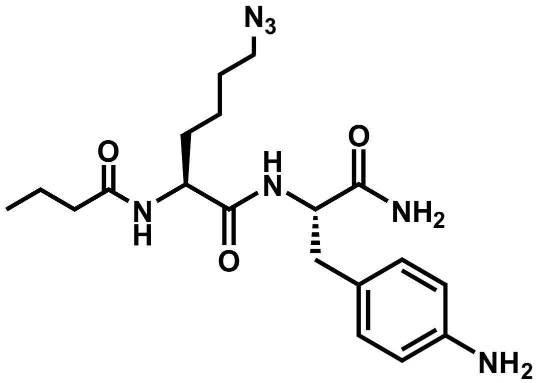


Analytical HPLC Chromatogram of **Sat3**

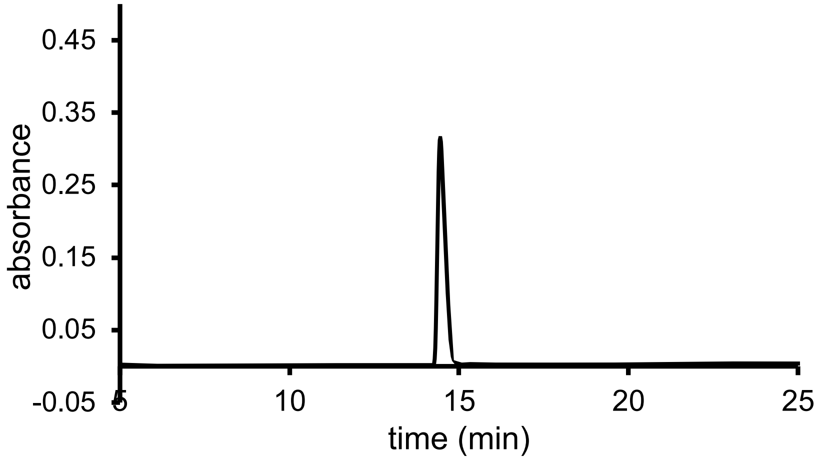


QTOF High Resolution Mass Spectrum for **Sat3** (m/z 404.2405 for $[M+H]^+$, found 404.2410)

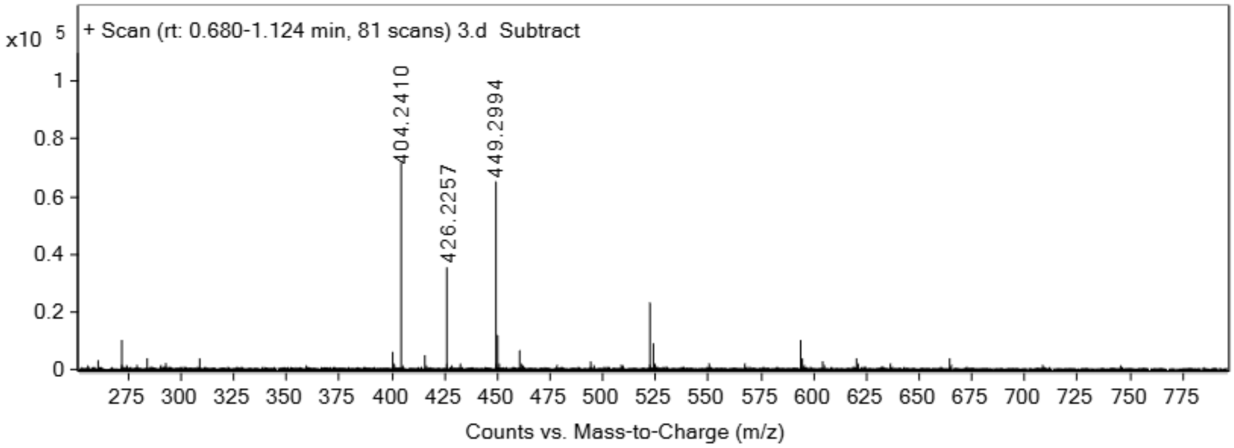

**Sat4**

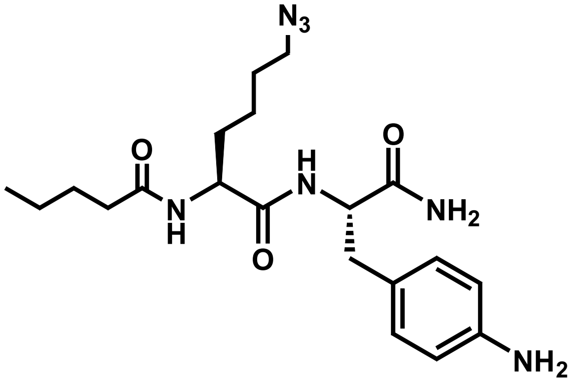

Analytical HPLC Chromatogram of **Sat4**

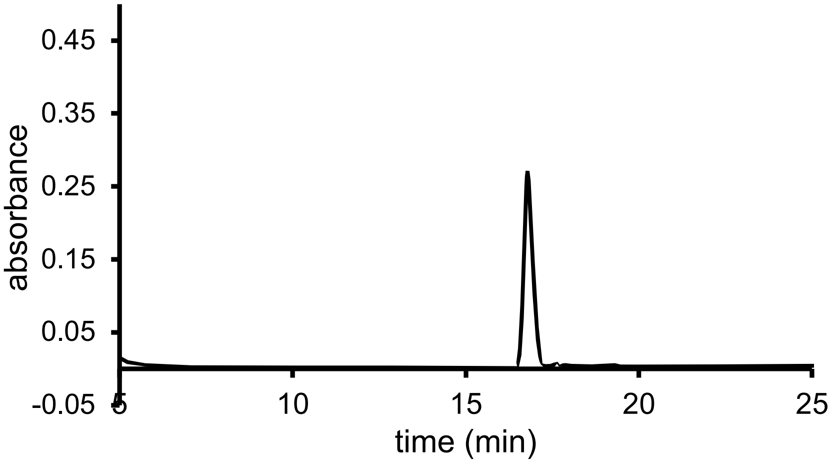


QTOF High Resolution Mass Spectrum for **Sat4** (m/z 418.2562 for $[M+H]^+$, found 418.2568)

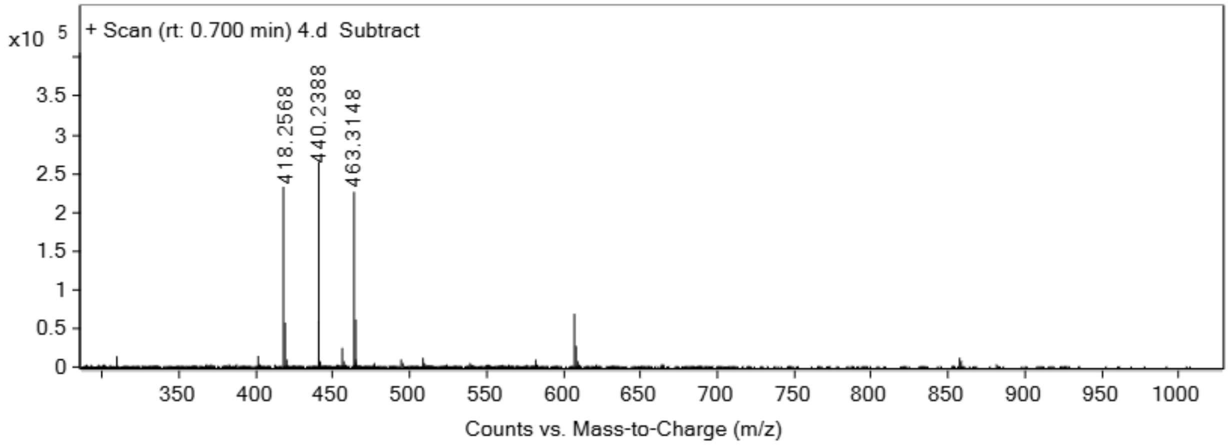

**Sat5**

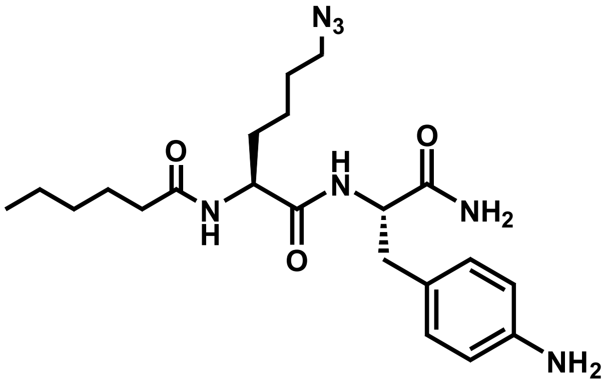

Analytical HPLC Chromatogram of **Sat5**

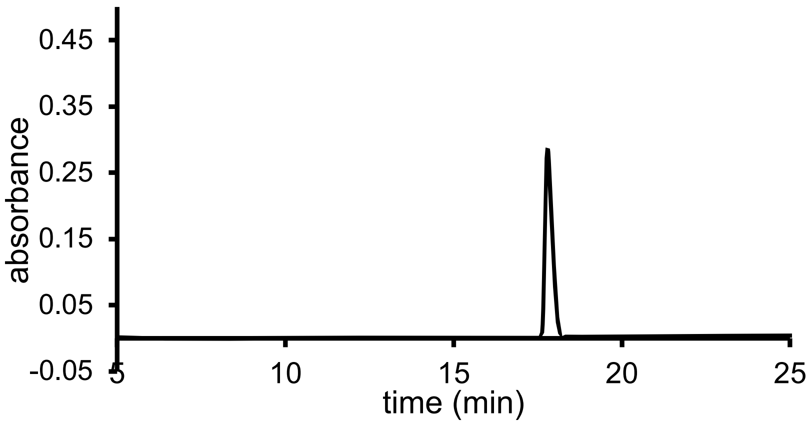


QTOF High Resolution Mass Spectrum for **Sat5** (m/z 432.2718 for $[M+H]^+$, found 432.2727)

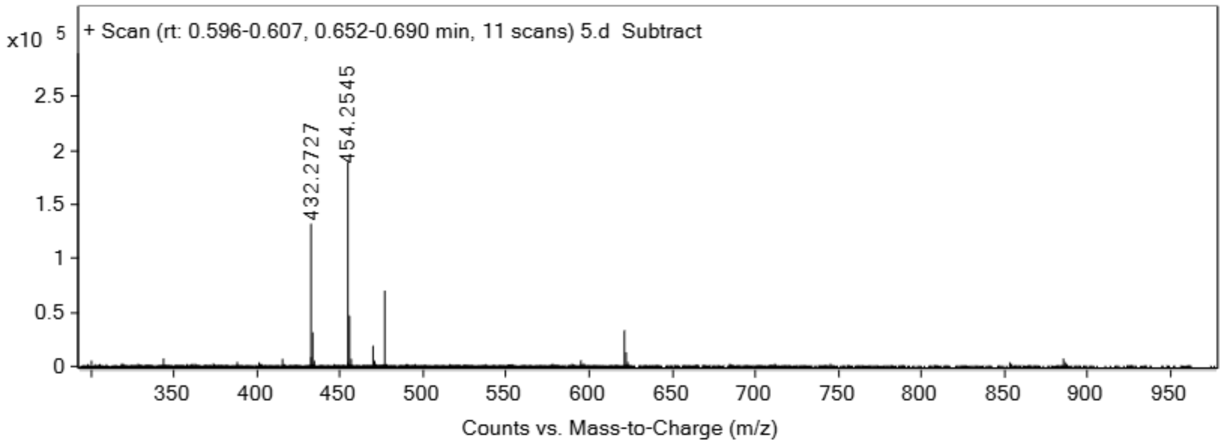

**Sat6**

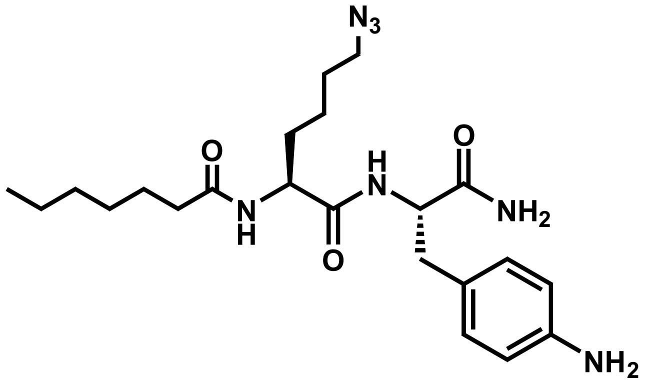


Analytical HPLC Chromatogram of **Sat6**

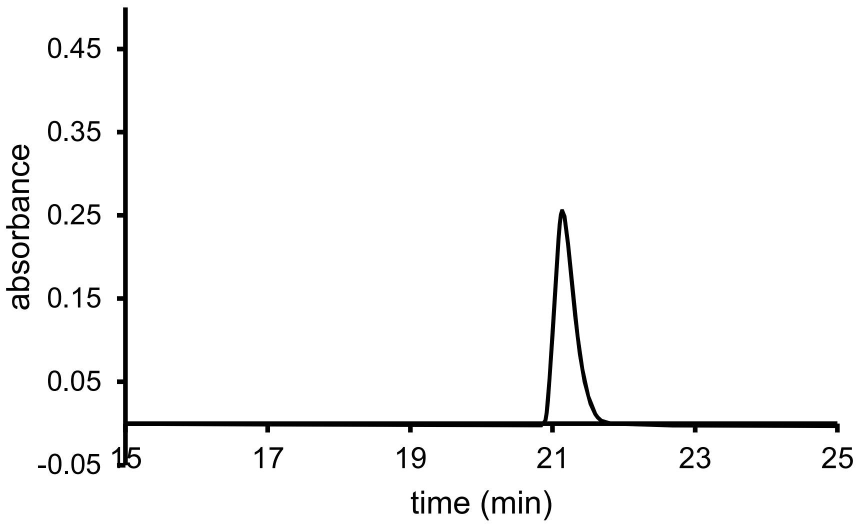


QTOF High Resolution Mass Spectrum for **Sat6** (m/z 446.2875 for $[M+H]^+$, found 446.2880)

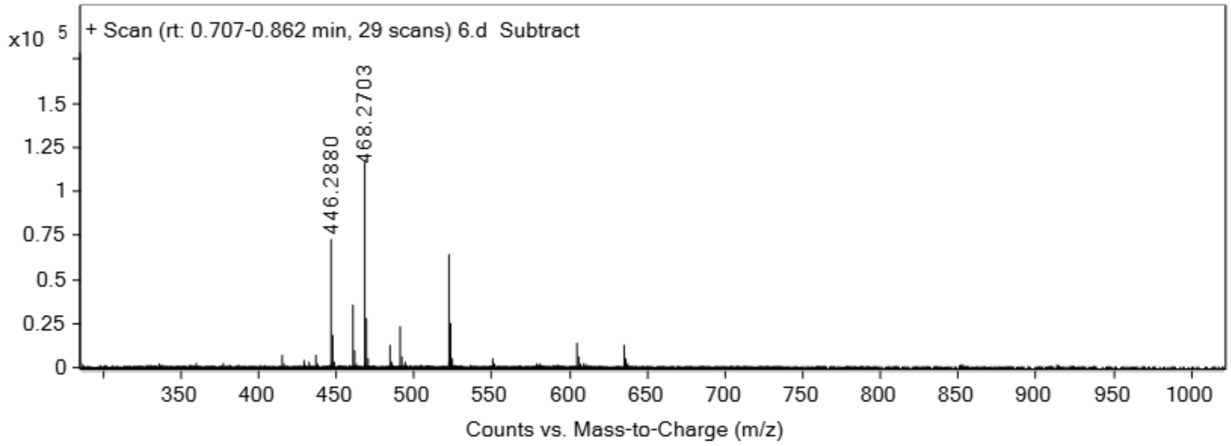


**Sat7**

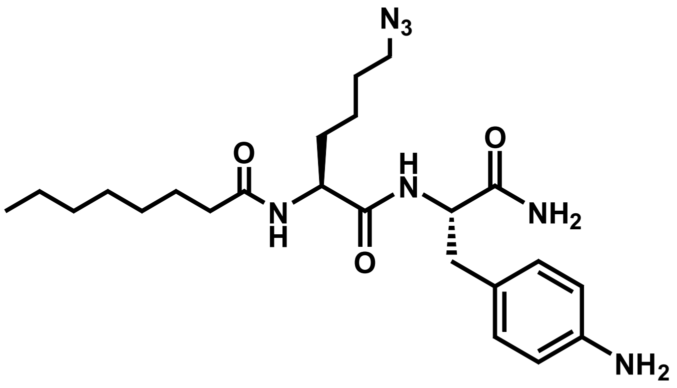


Analytical HPLC Chromatogram of **Sat7**

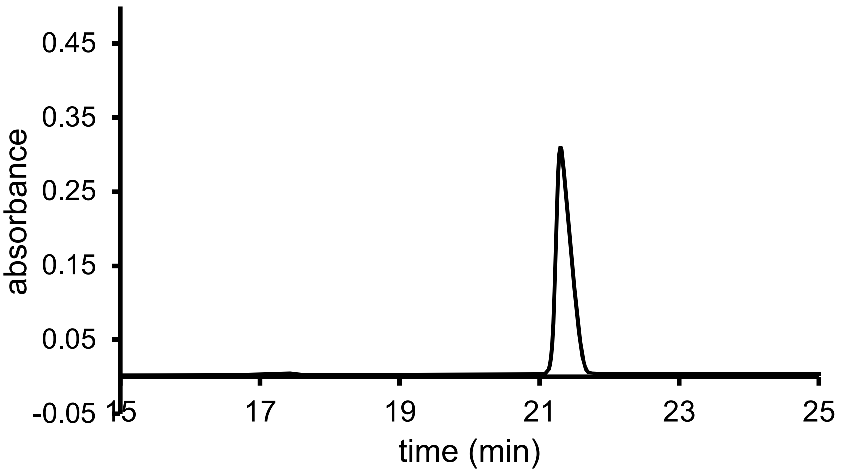


QTOF High Resolution Mass Spectrum for **Sat7** (m/z 460.3031 for $[M+H]^+$, found 460.3029)

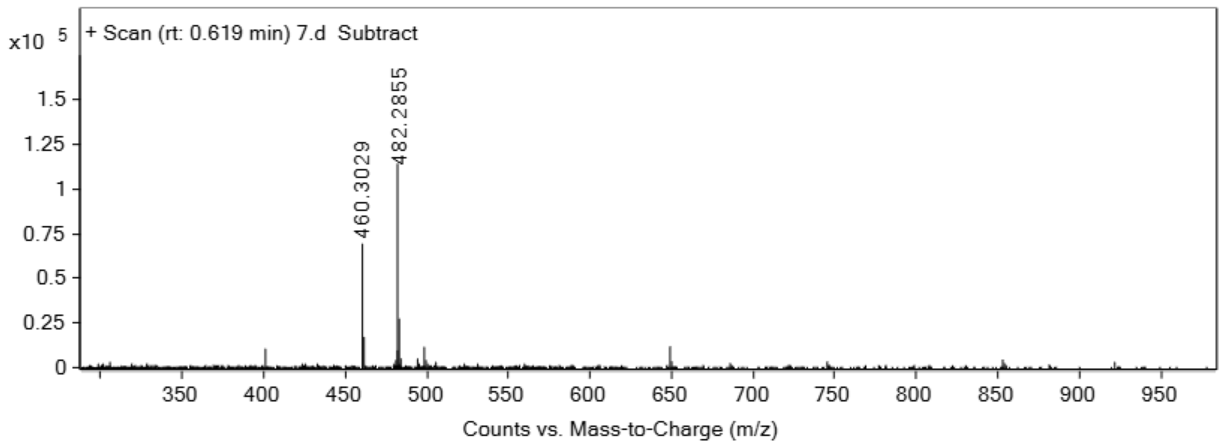


**Sat8**

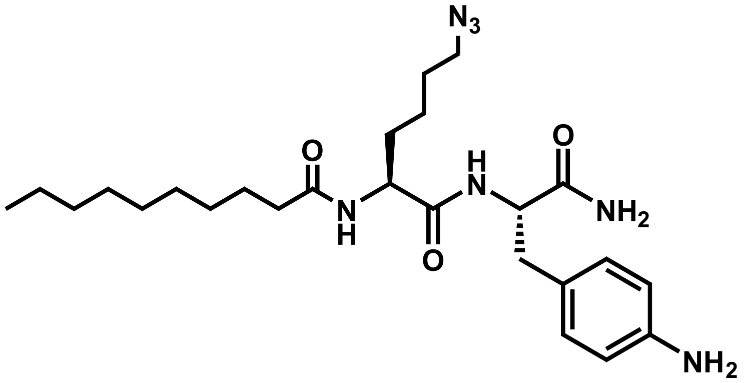


Analytical HPLC Chromatogram of **Sat8**

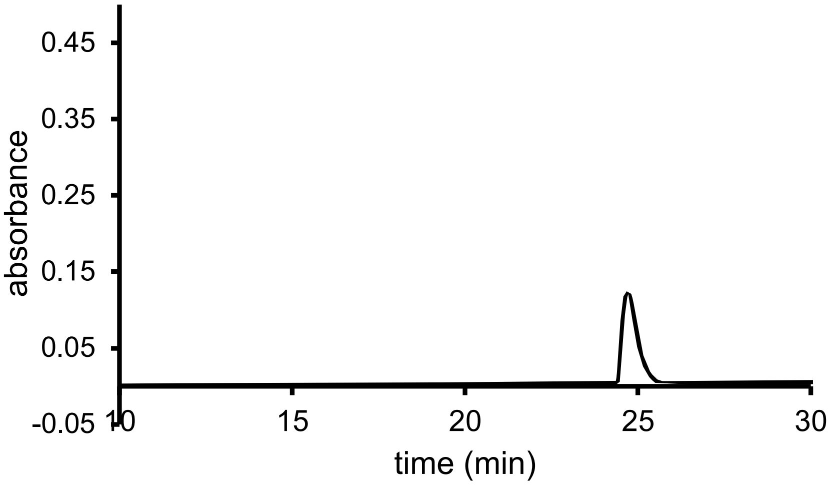


QTOF High Resolution Mass Spectrum for **Sat8** (m/z 510.3164 for [M+Na]$^+$, found 510.3162)

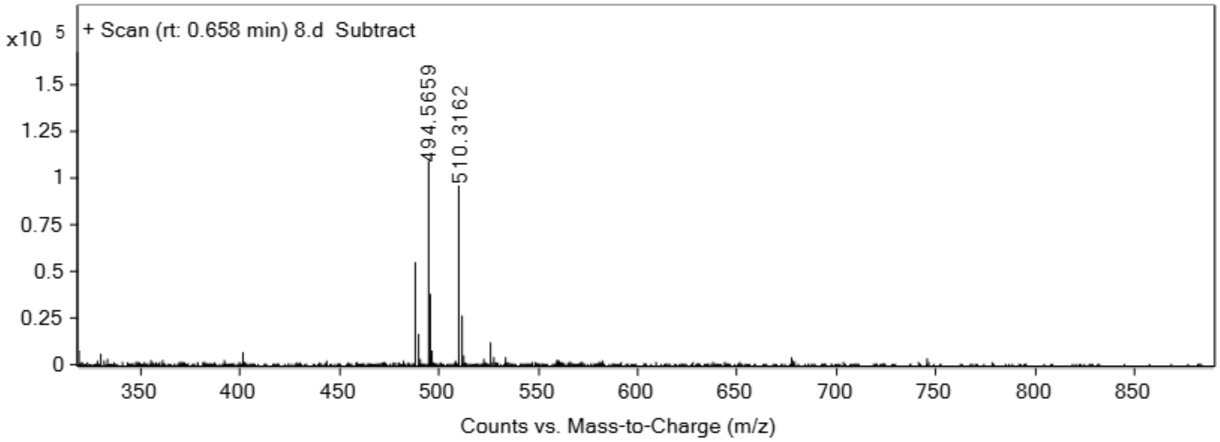


**Sat9**

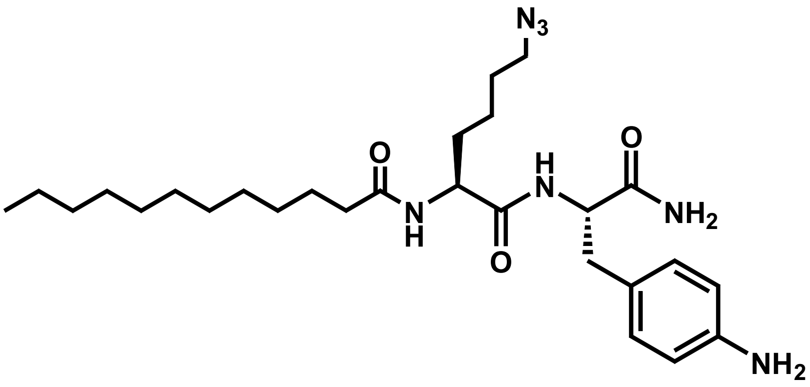


Analytical HPLC Chromatogram of **Sat9**

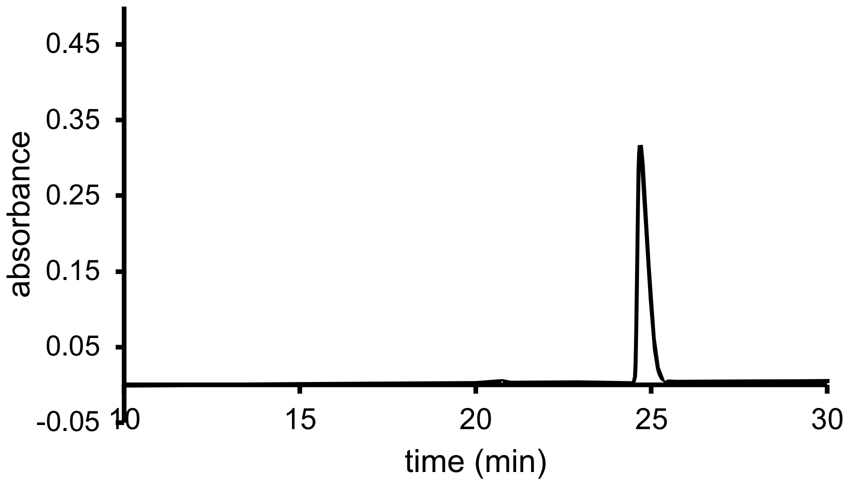


QTOF High Resolution Mass Spectrum for **Sat9** (m/z 516.3657 for $[M+H]^+$, found 516.3653)

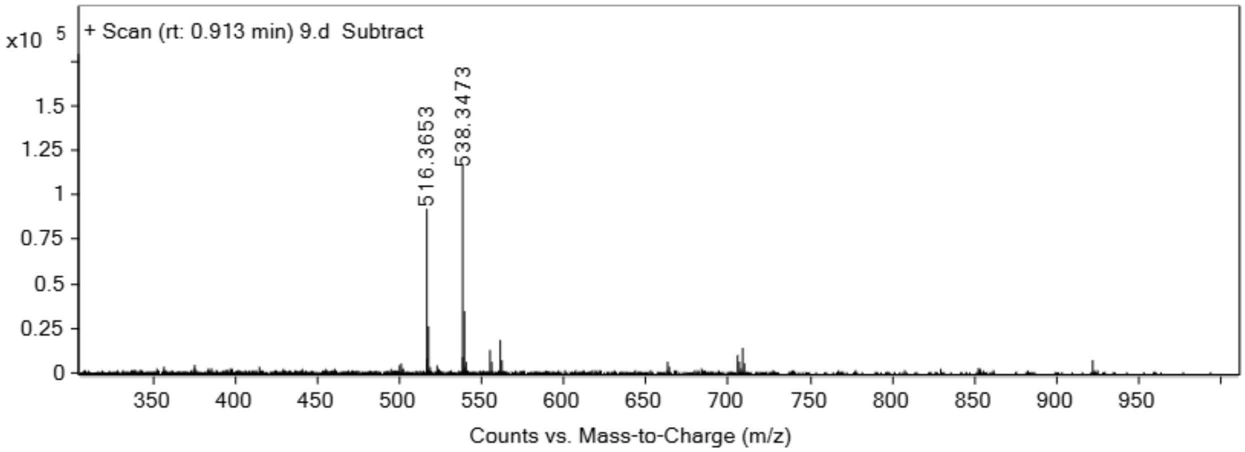


**Sat10**

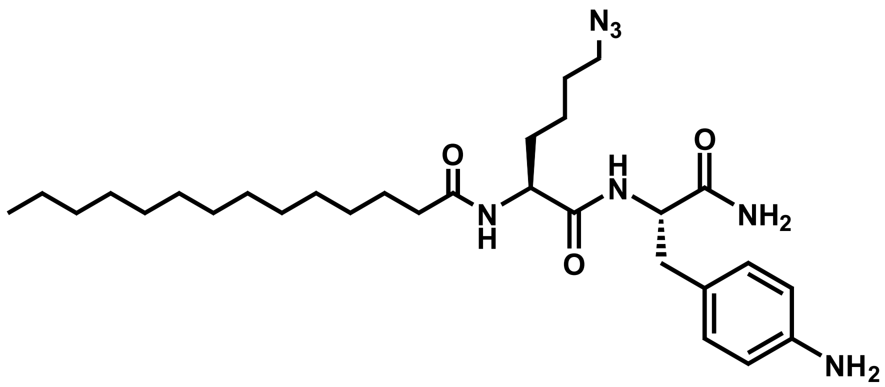


Analytical HPLC Chromatogram of **Sat10**

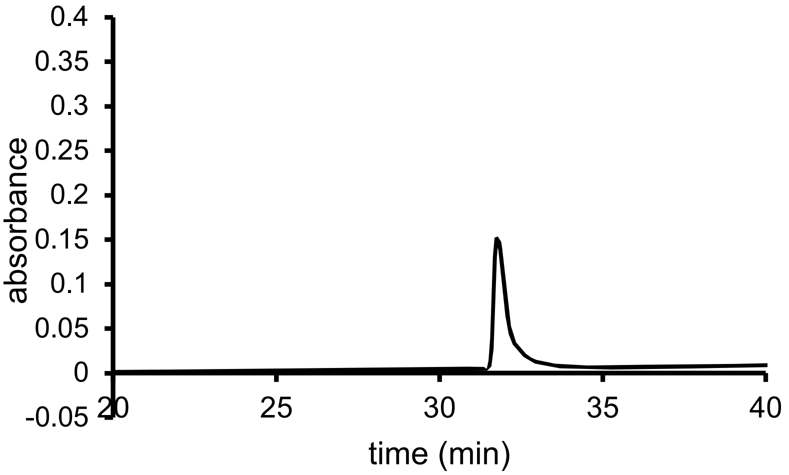


QTOF High Resolution Mass Spectrum for **Sat10** (m/z 544.3970 for $[M+H]^+$, found 544.3956)

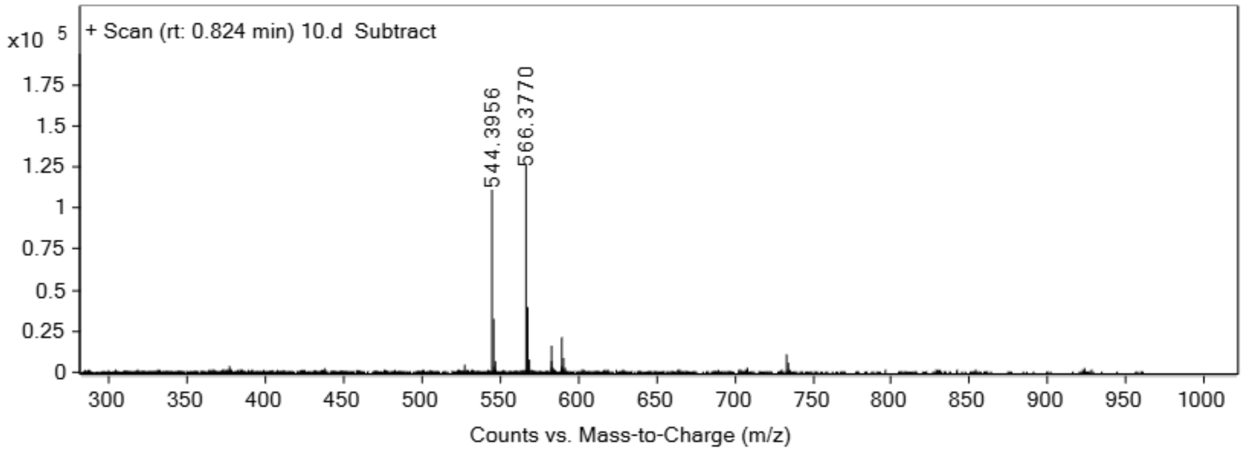


**Unsat1**

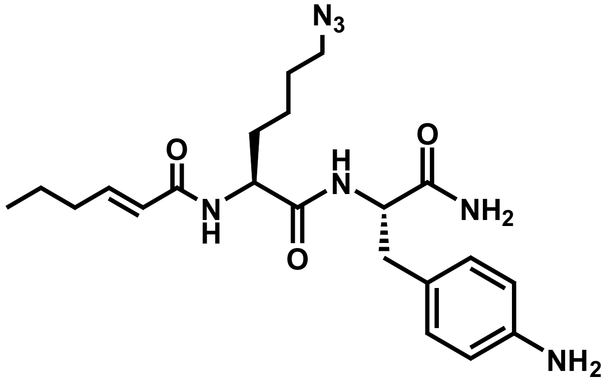


Analytical HPLC Chromatogram of **Unsat1**

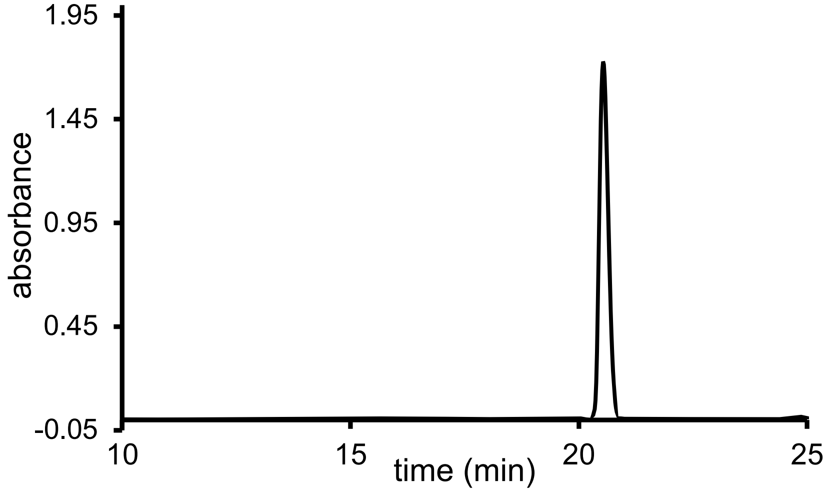

QTOF High Resolution Mass Spectrum for **Unsat1** (m/z 430.2562 for $[M+H]^+$, found 430.2578)

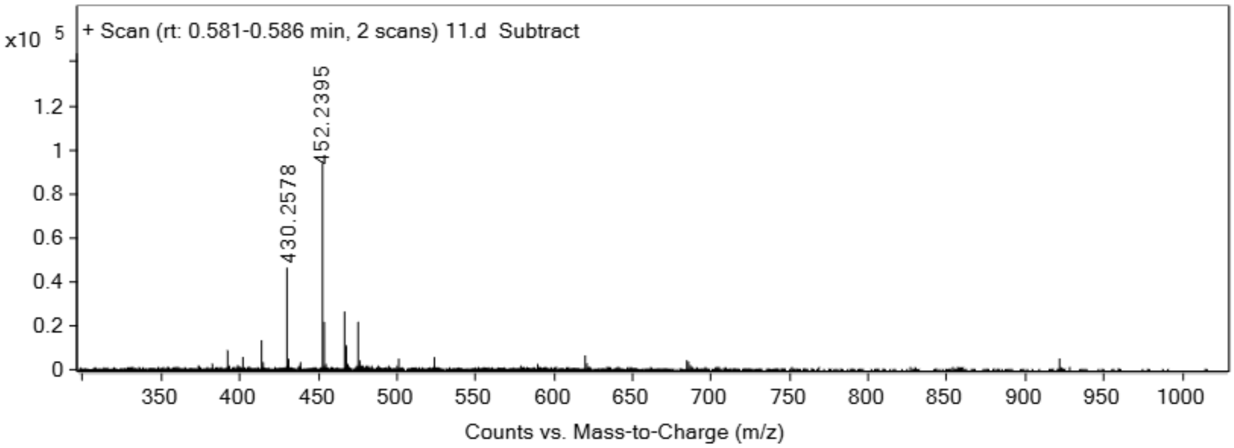


**Unsat2**

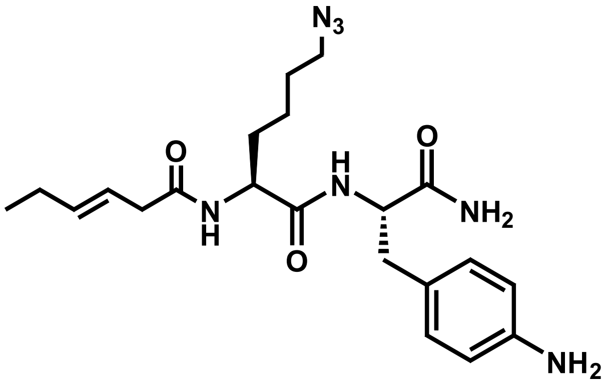


Analytical HPLC Chromatogram of **Unsat2**

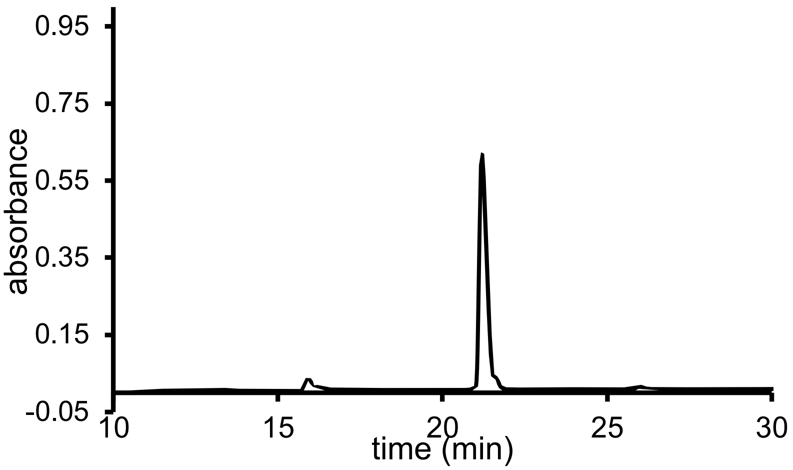

QTOF High Resolution Mass Spectrum for **Unsat2** (m/z 430.2562 for $[M+H]^+$, found 430.2558)

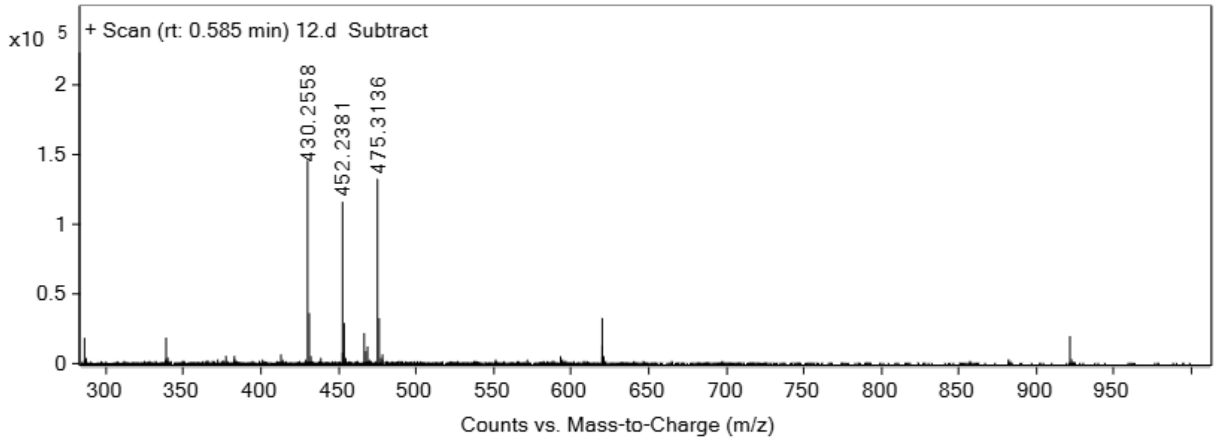


**Unsat3**

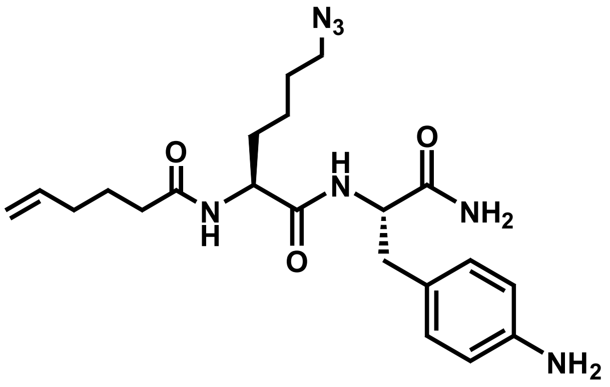


Analytical HPLC Chromatogram of **Unsat3**

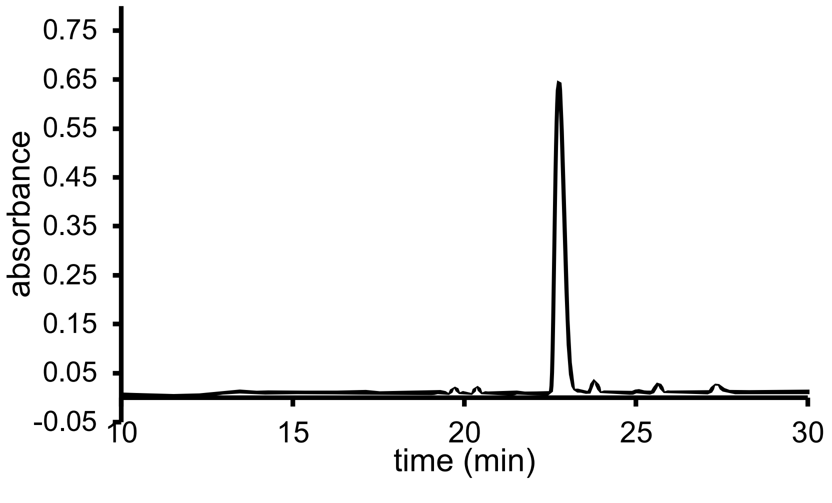

QTOF High Resolution Mass Spectrum for **Unsat3** (m/z 430.2562 for $[M+H]^+$, found 430.2563)

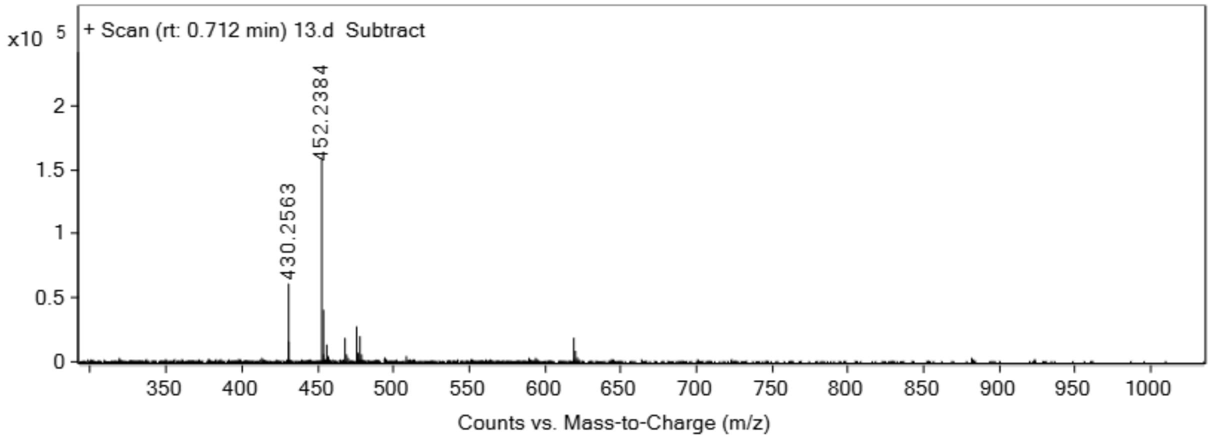


**Unsat4**

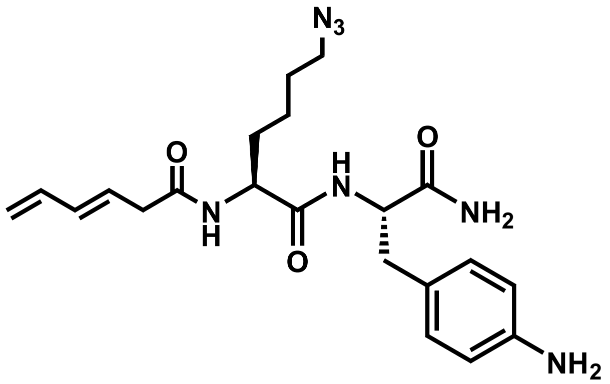


Analytical HPLC Chromatogram of **Unsat4**

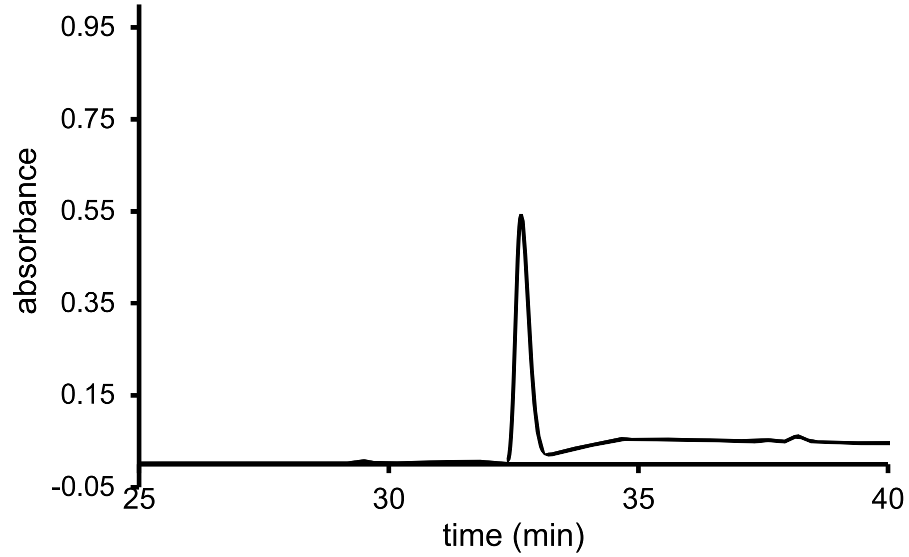


QTOF High Resolution Mass Spectrum for **Unsat4** (m/z 428.2405 for $[M+H]^+$, found 428.2403)

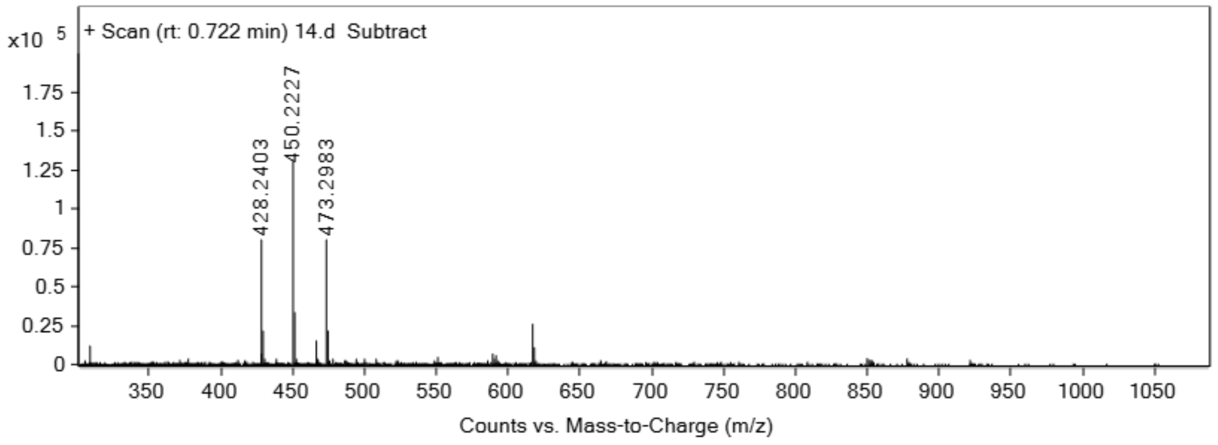


**Cyc1**

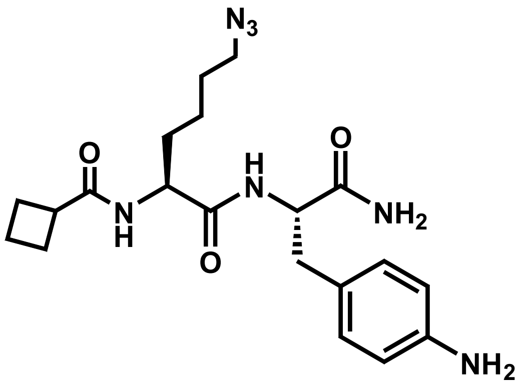


Analytical HPLC Chromatogram of **Cyc1**

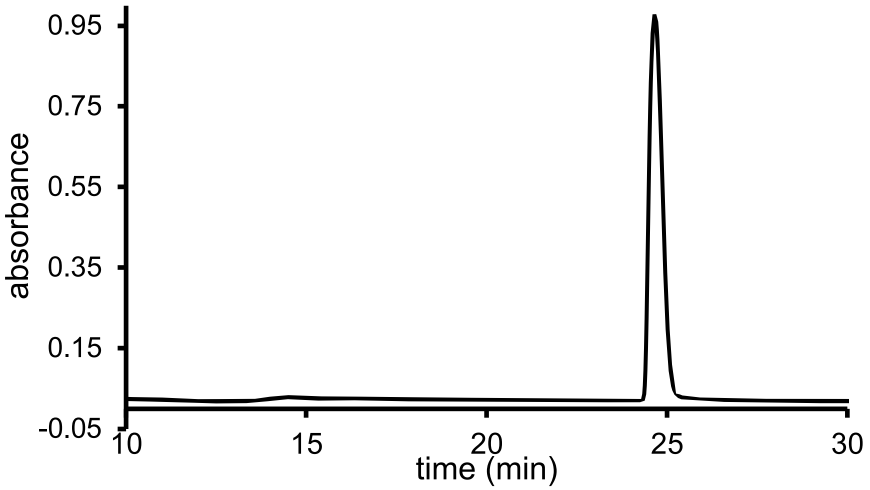


QTOF High Resolution Mass Spectrum for **Cyc1** (m/z 416.2405 for $[M+H]^+$, found 416.2412)

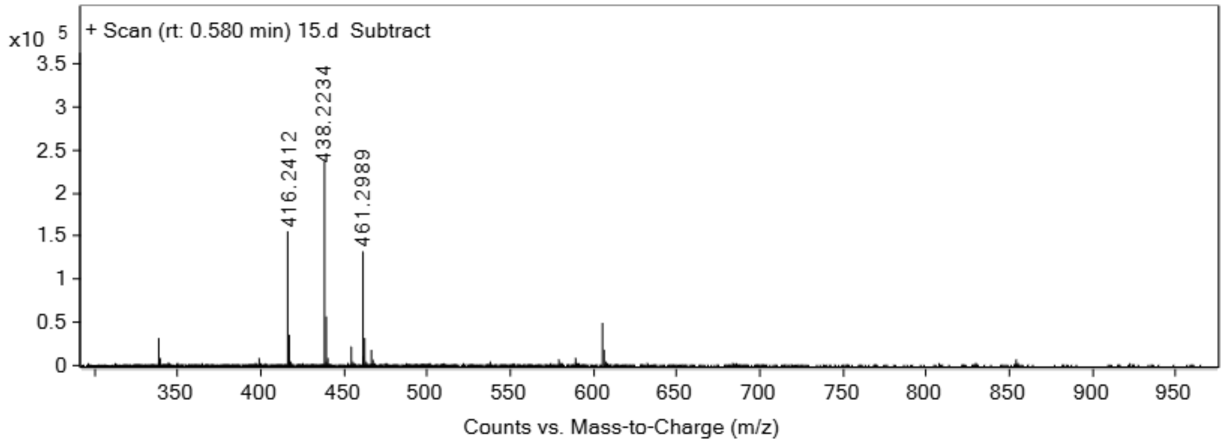


**Cyc2**

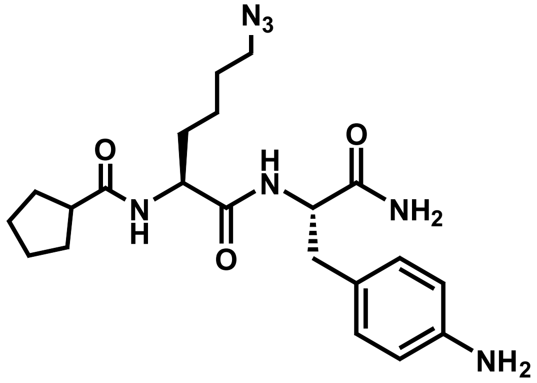


Analytical HPLC Chromatogram of **Cyc2**

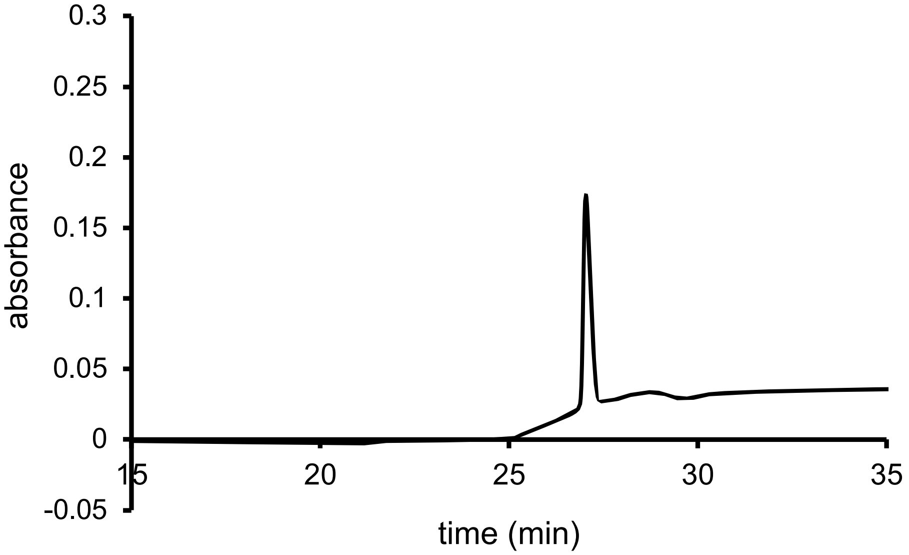


QTOF High Resolution Mass Spectrum for **Cyc2** (m/z 430.2562 for $[M+H]^+$, found 416.2578)

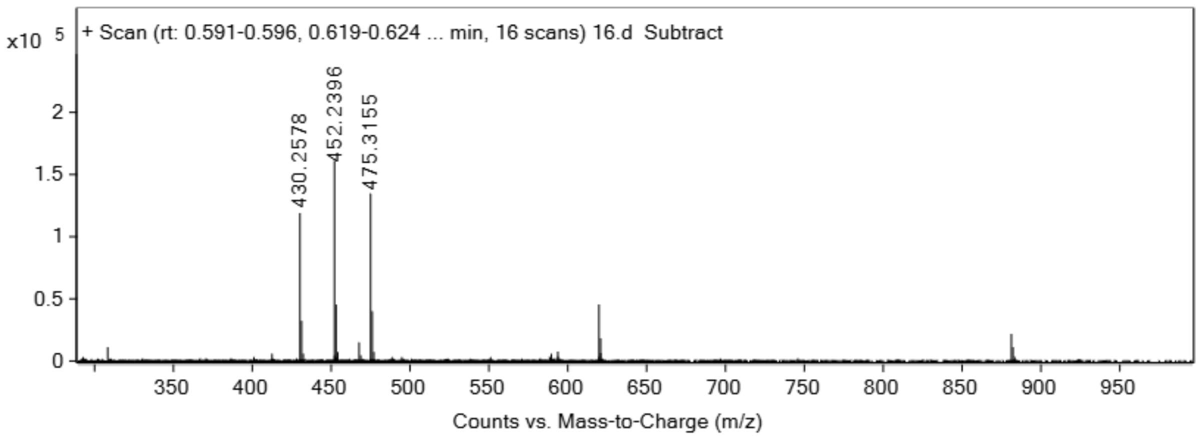


**Cyc3**

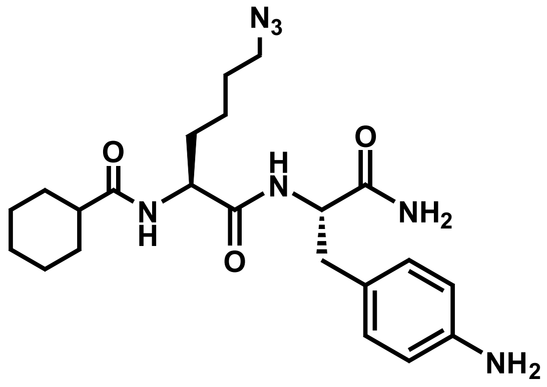

Analytical HPLC Chromatogram of **Cyc3**

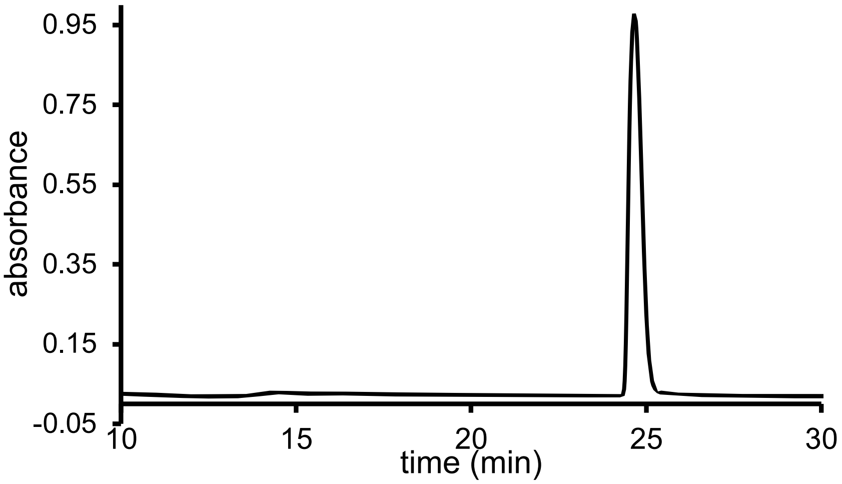


QTOF High Resolution Mass Spectrum for **Cyc3** (m/z 444.2718 for $[M+H]^+$, found 444.2718)

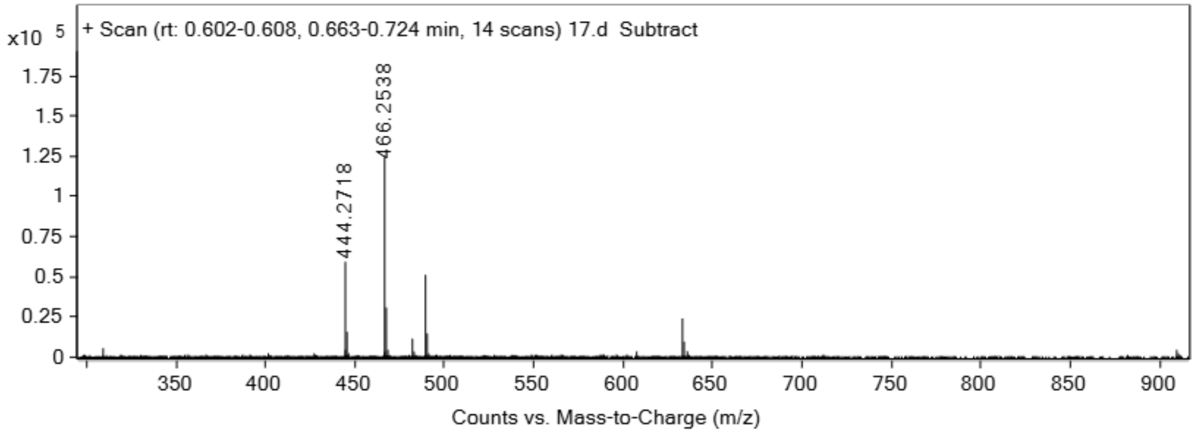


**Cyc4**

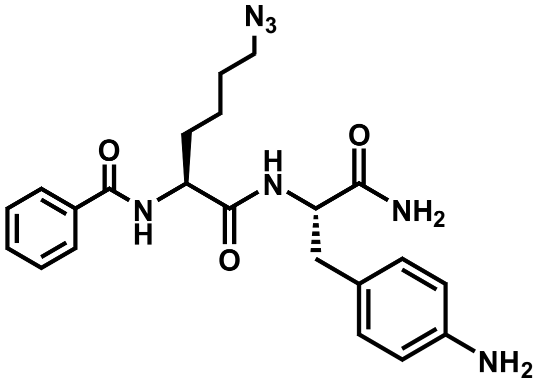

Analytical HPLC Chromatogram of **Cyc4**

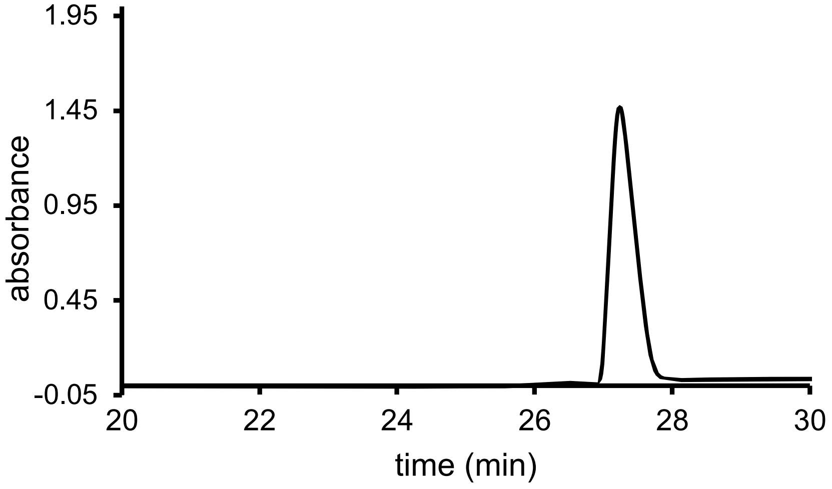


QTOF High Resolution Mass Spectrum for **Cyc4** (m/z 438.2249 for $[M+H]^+$, found 438.2266)

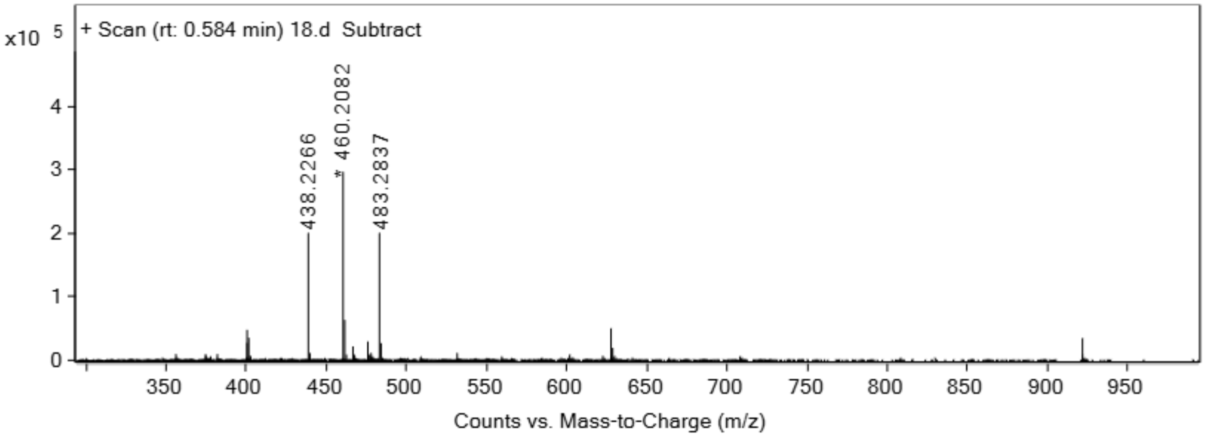


**Cyc5**

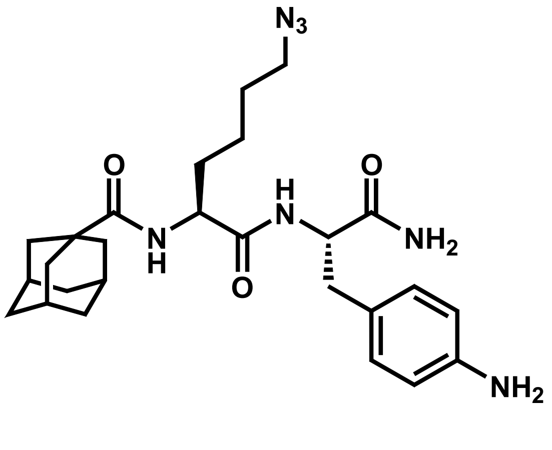

Analytical HPLC Chromatogram of **Cyc5**

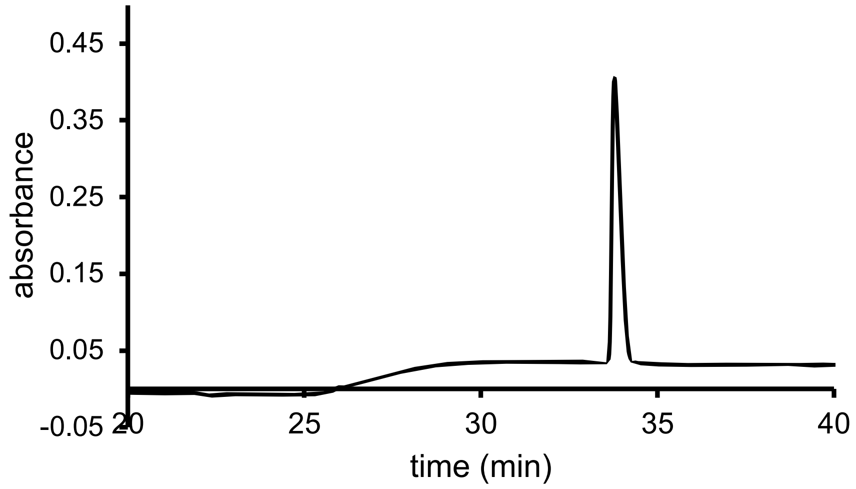


QTOF High Resolution Mass Spectrum for **Cyc5** (m/z 496.3031 for $[M+H]^{+}$, found 496.3051)

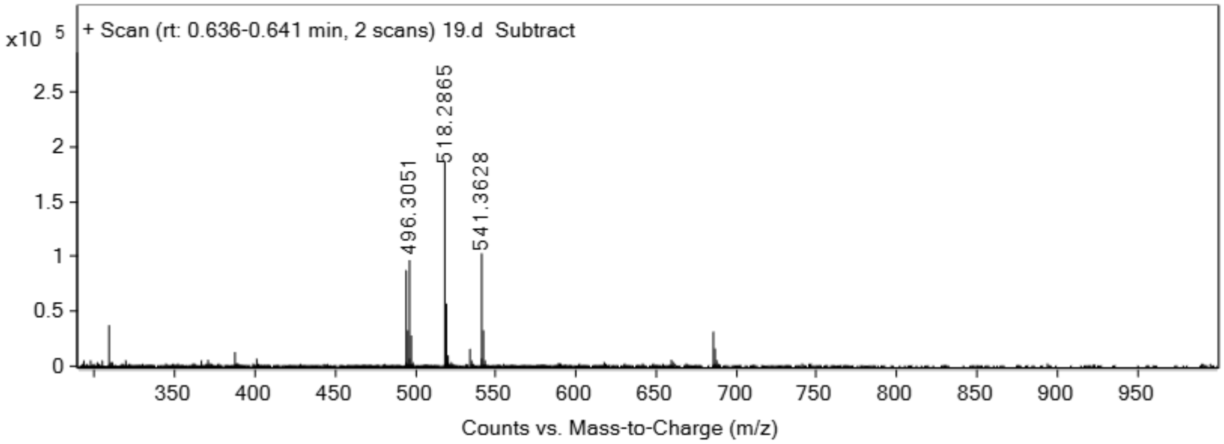


**Cyc6**

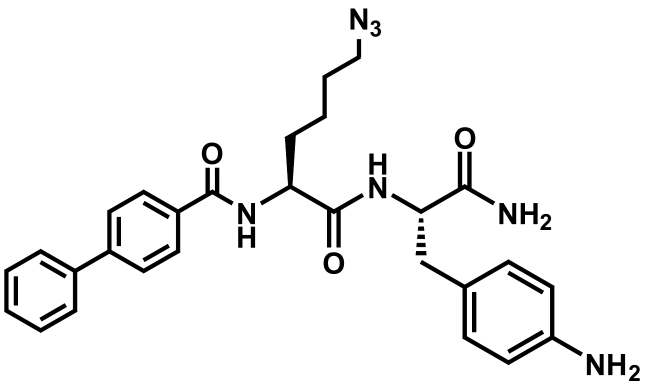


Analytical HPLC Chromatogram of **Cyc6**

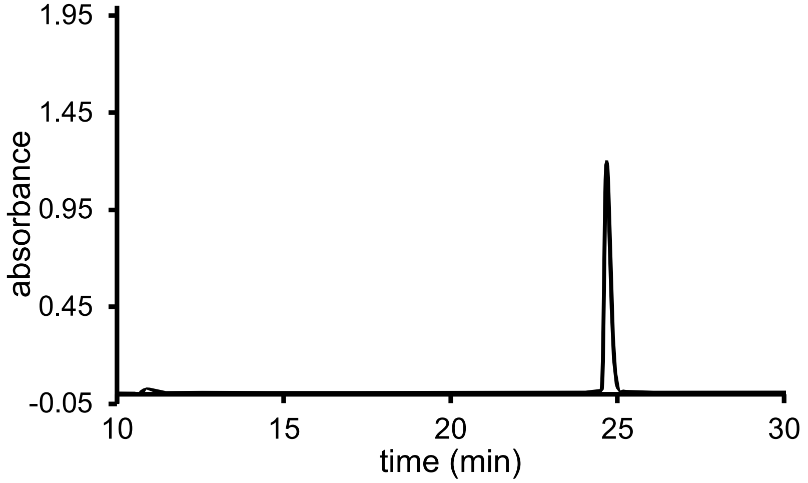


QTOF High Resolution Mass Spectrum for **Cyc6** (m/z 514.2562 for $[M+H]^+$, found 514.2577)

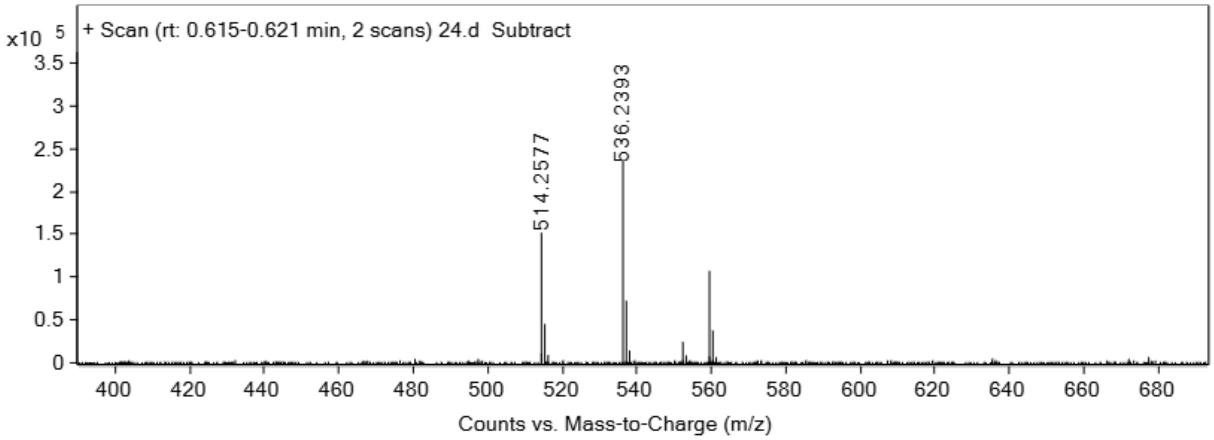


**Brn1**

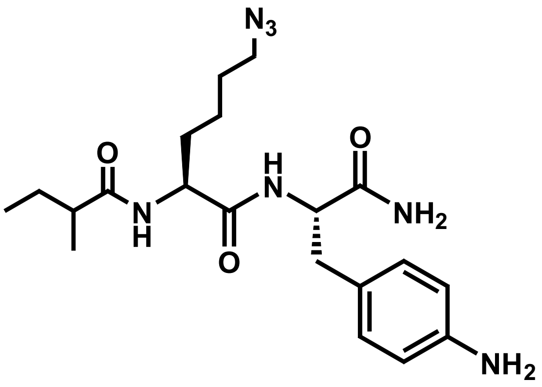

Analytical HPLC Chromatogram of **Brn1**

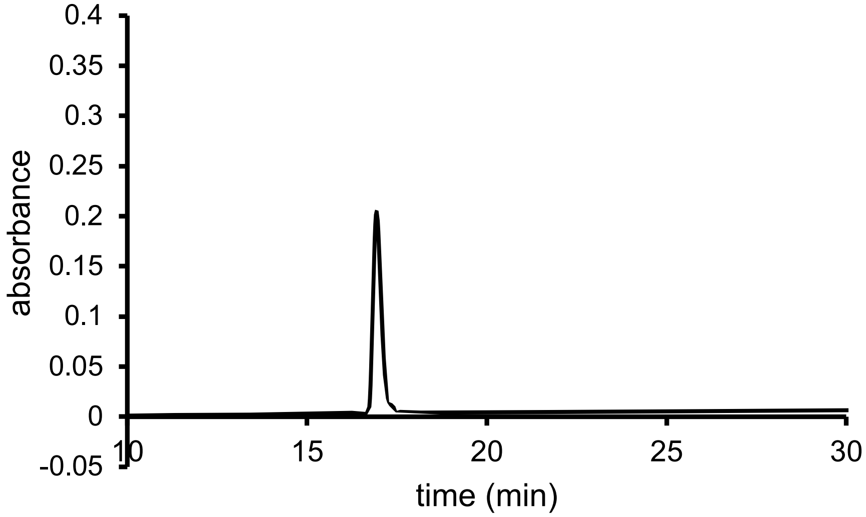


QTOF High Resolution Mass Spectrum for **Brn1** (m/z 418.2562 for $[M+H]^+$, found 418.2580)

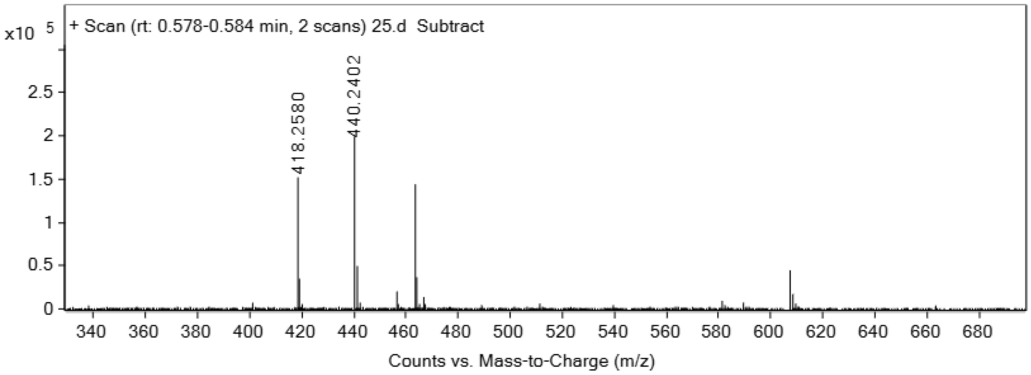


**Brn2**

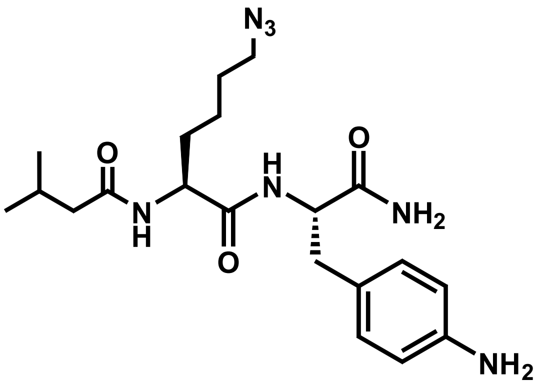


Analytical HPLC Chromatogram of **Brn2**

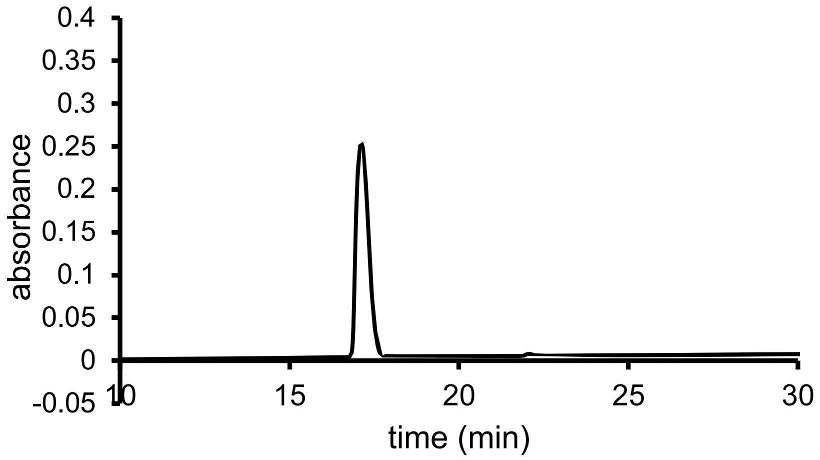


QTOF High Resolution Mass Spectrum for **Brn2** (m/z 418.2562 for $[M+H]^+$, found 418.2593)

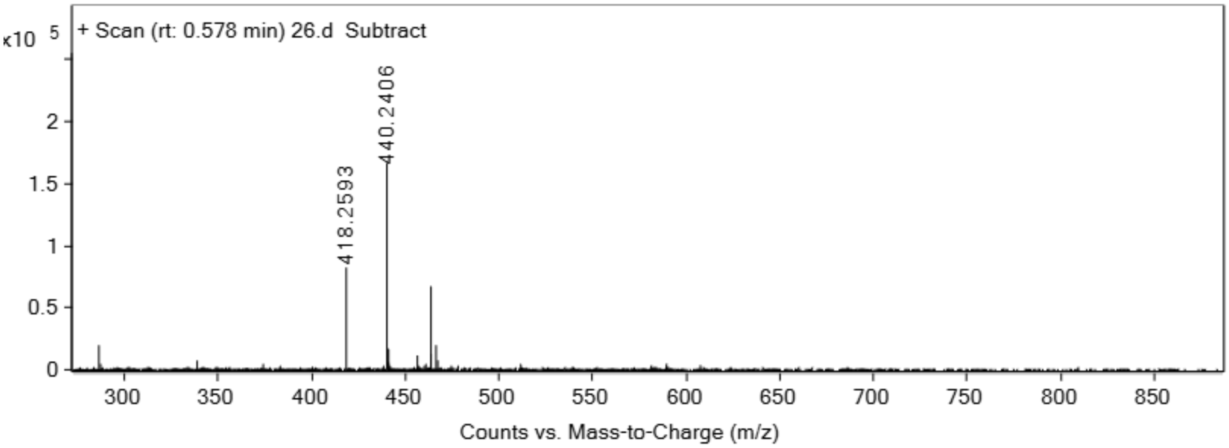


**Brn3**

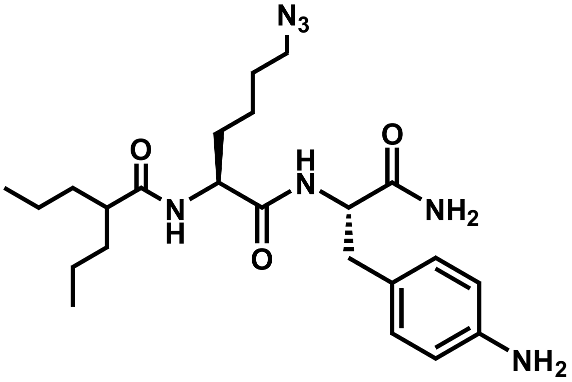


Analytical HPLC Chromatogram of **Brn3**

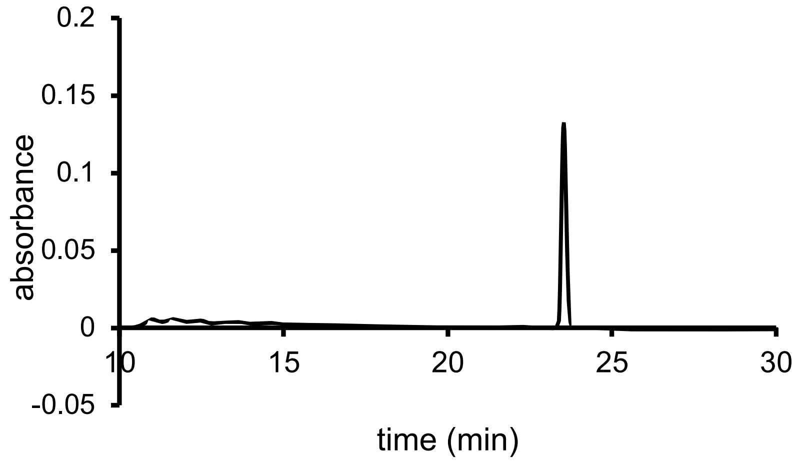


QTOF High Resolution Mass Spectrum for **Brn3** (m/z 460.3032 for $[M+H]^+$, found 460.3031)

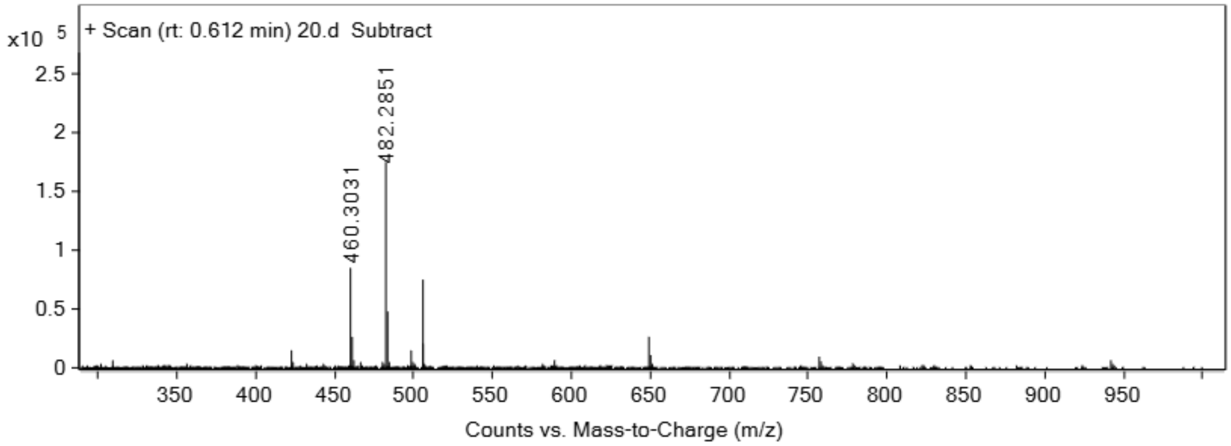


**Het1**

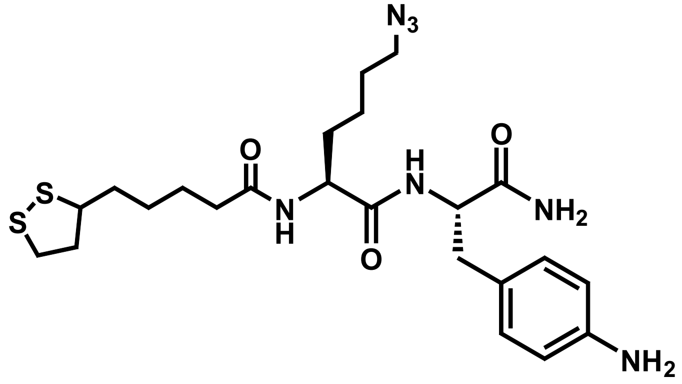


Analytical HPLC Chromatogram of **Het1**

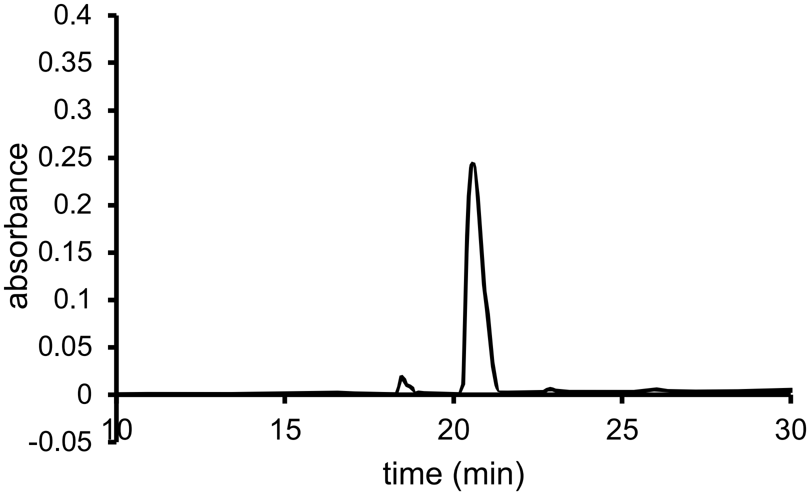


QTOF High Resolution Mass Spectrum for **Het1** (m/z 522.2316 for [M+H]$^{+}$, found 522.2341)

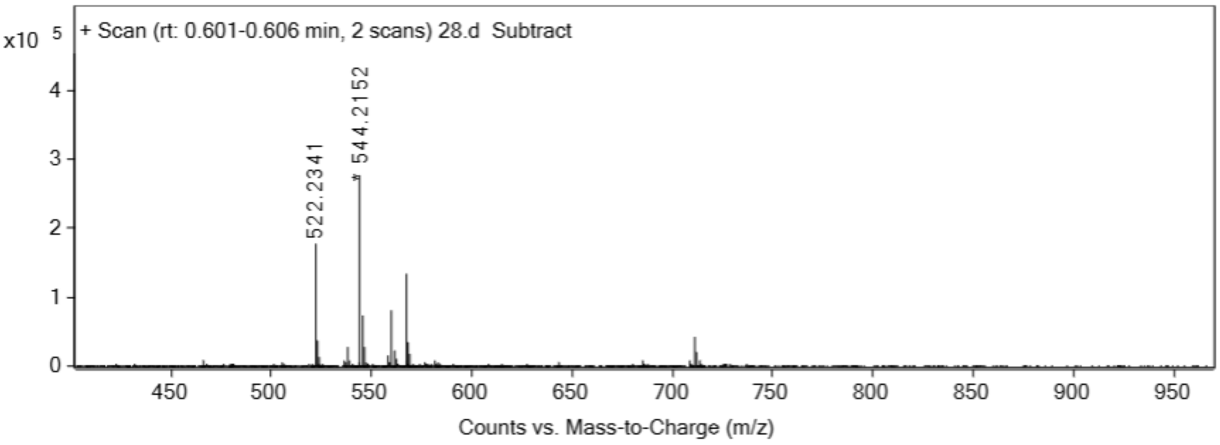


**Het2**

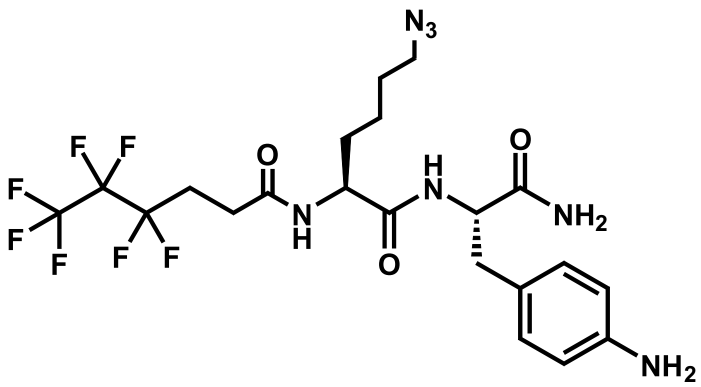


Analytical HPLC Chromatogram of **Het2**

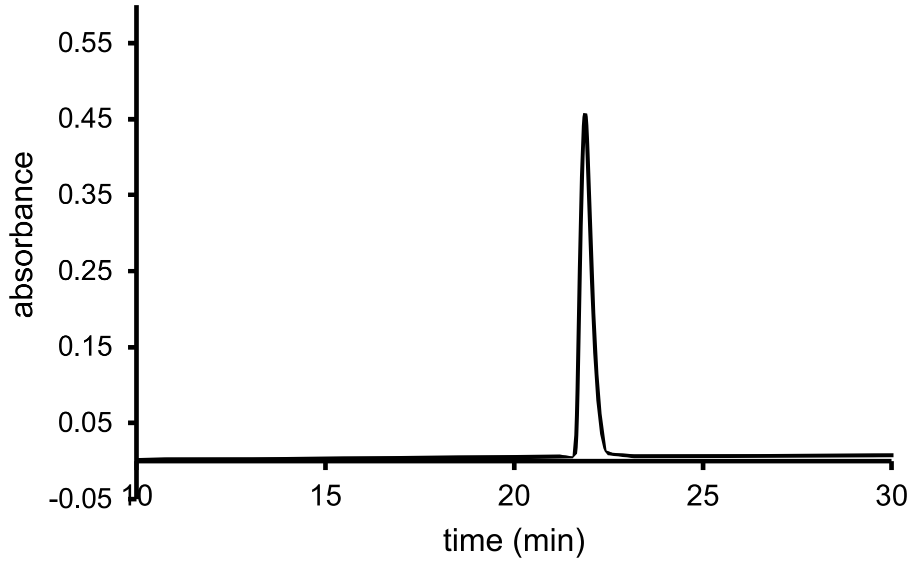


QTOF High Resolution Mass Spectrum for **Het2** (m/z 558. 2058 for $[M+H]^+$, found 558.2053)

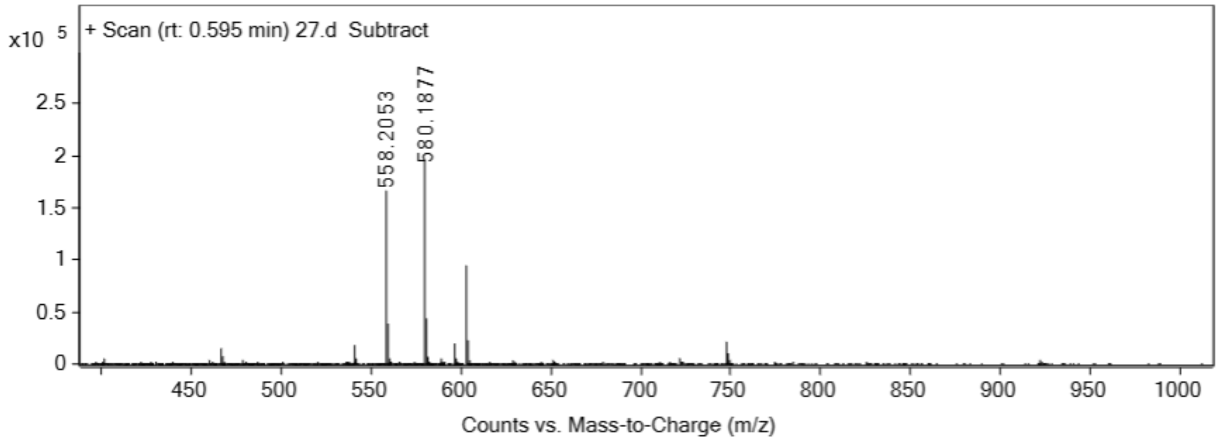


**Mix1**

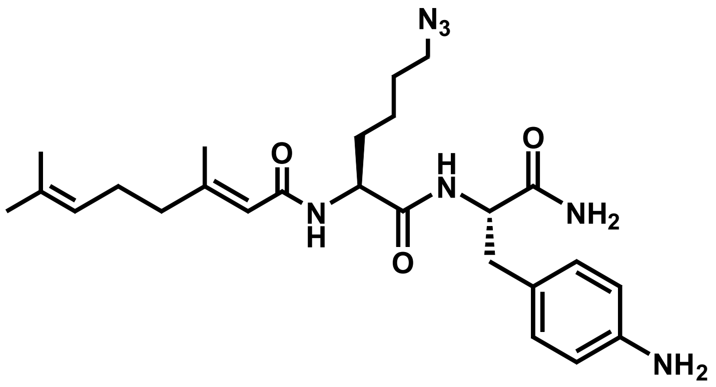


Analytical HPLC Chromatogram of **Mix1**

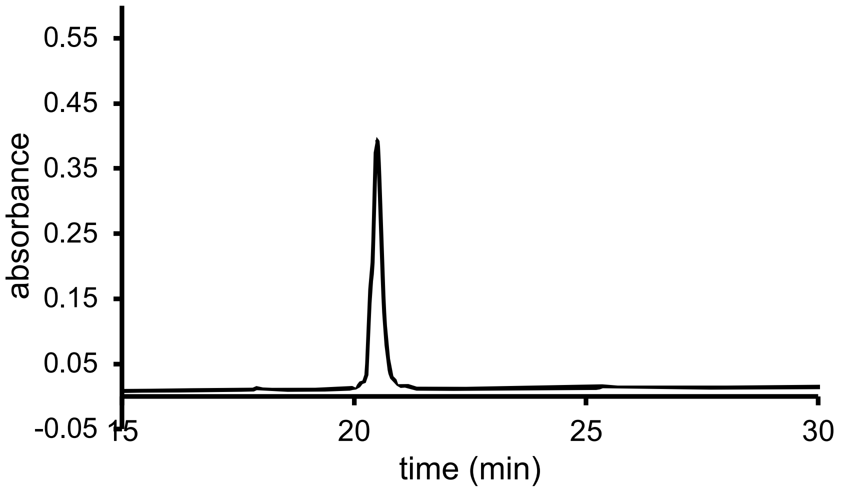


QTOF High Resolution Mass Spectrum for **Mix1** (m/z 484.3031 for [M+H]$^+$, found 484.3024)

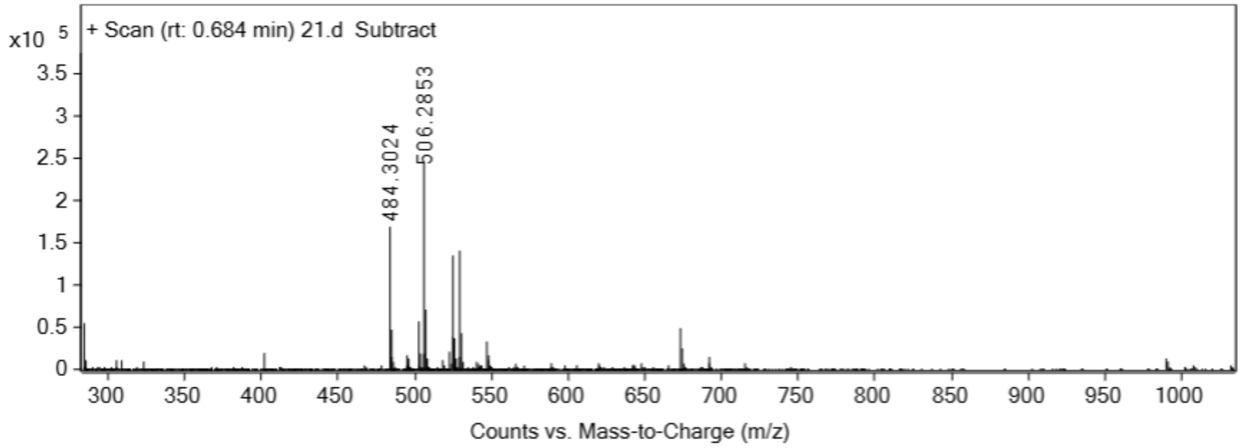


**Mix2**

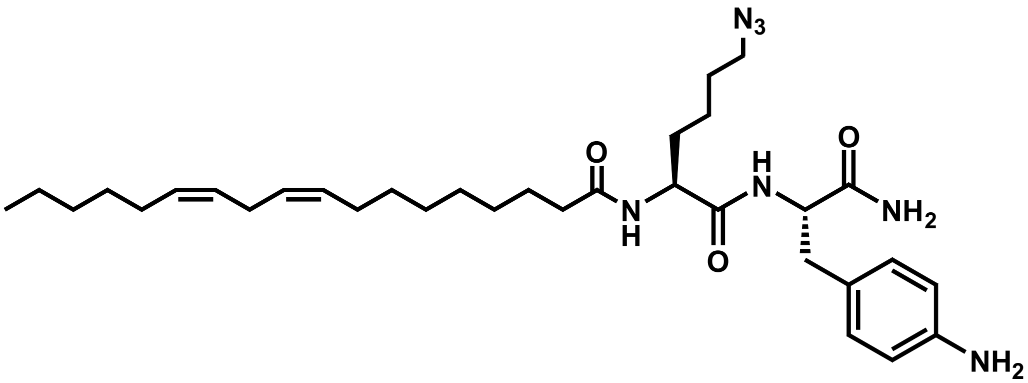


Analytical HPLC Chromatogram of **Mix2**

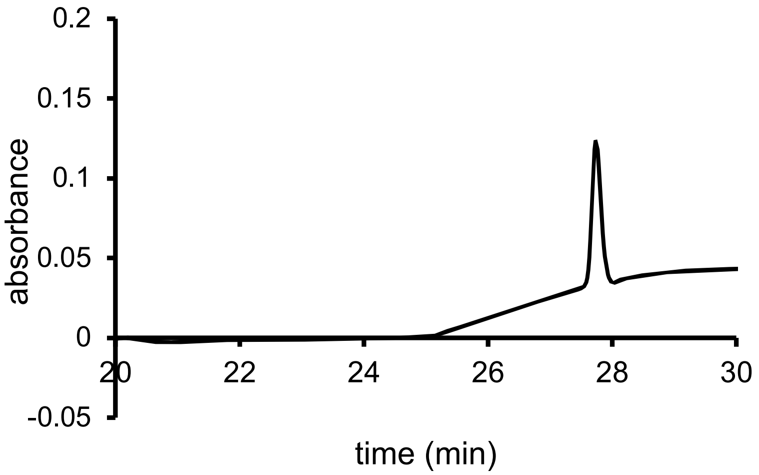


QTOF High Resolution Mass Spectrum for **Mix2** (m/z 596.4283 for $[M+H]^+$, found 596.4301)

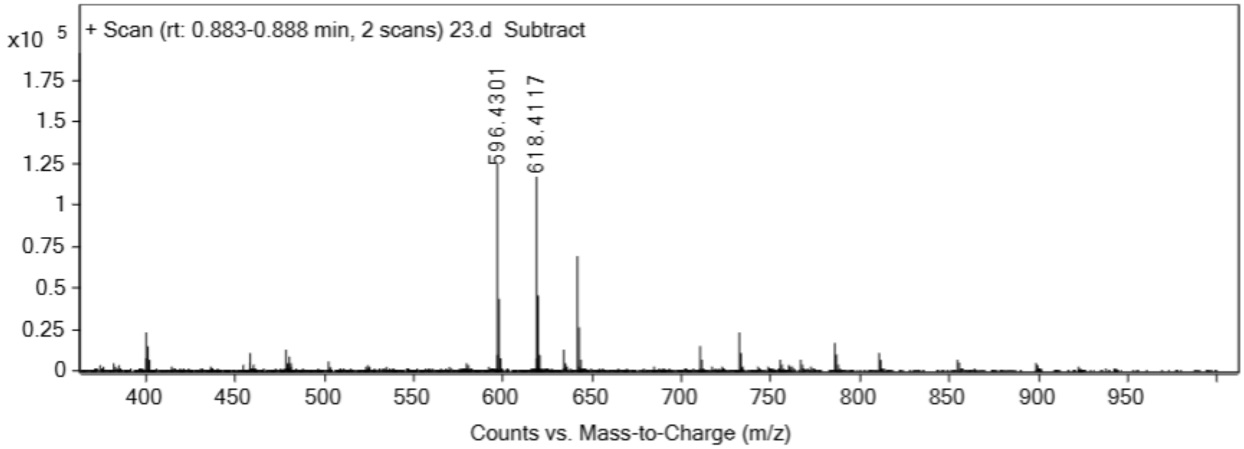


**Mix3**

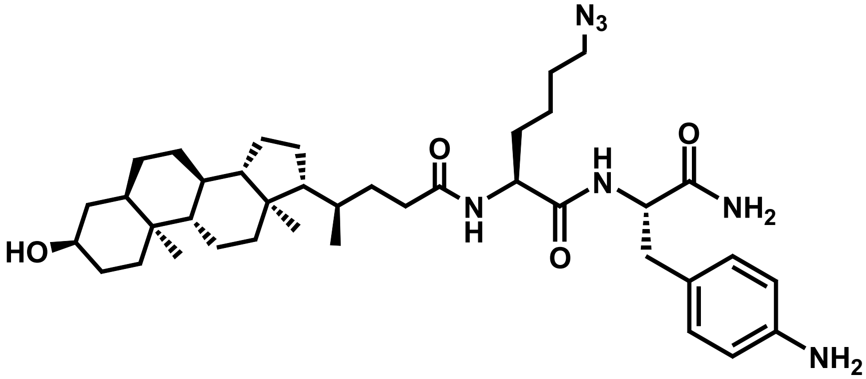


Analytical HPLC Chromatogram of **Mix3**

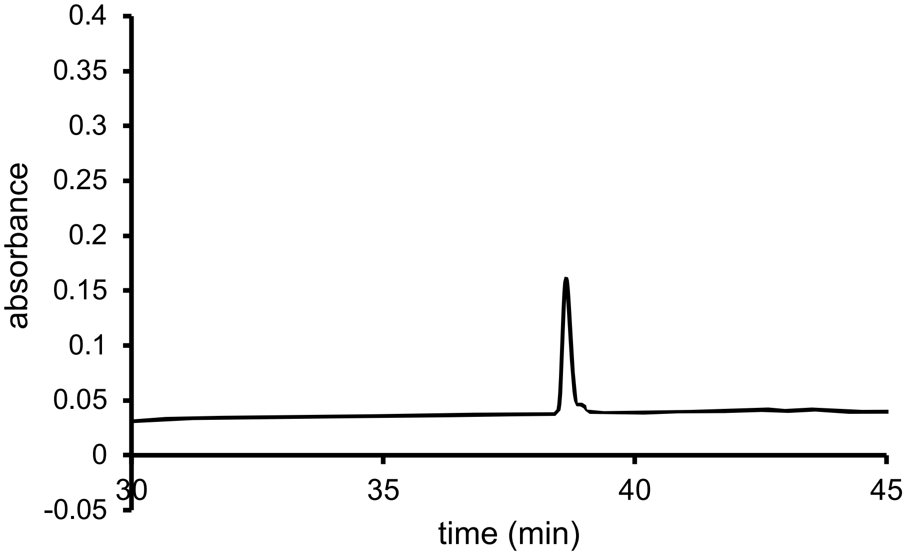


QTOF High Resolution Mass Spectrum for **Mix3** (m/z 692.4858 for $[M+H]^+$, found 692.4869)

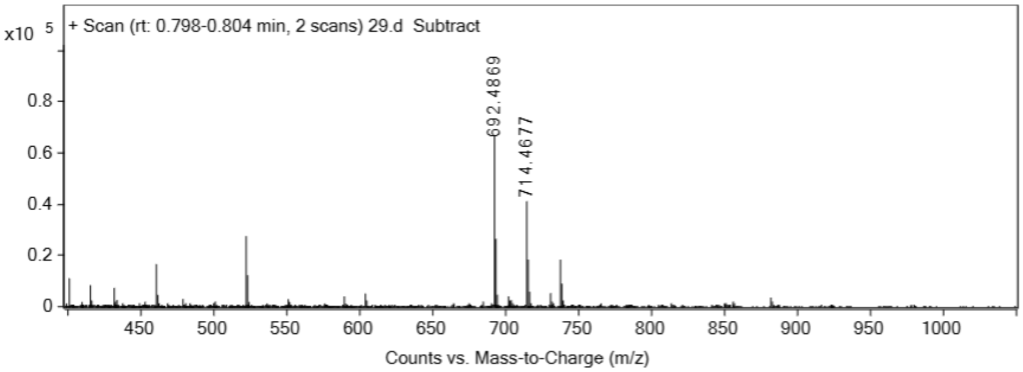


**7b**

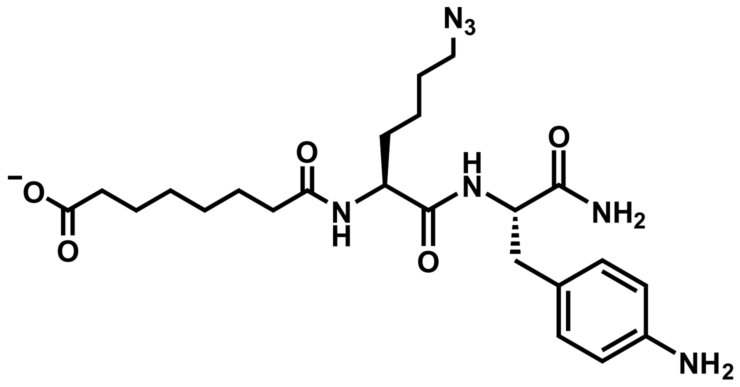


Analytical HPLC Chromatogram of **7b**

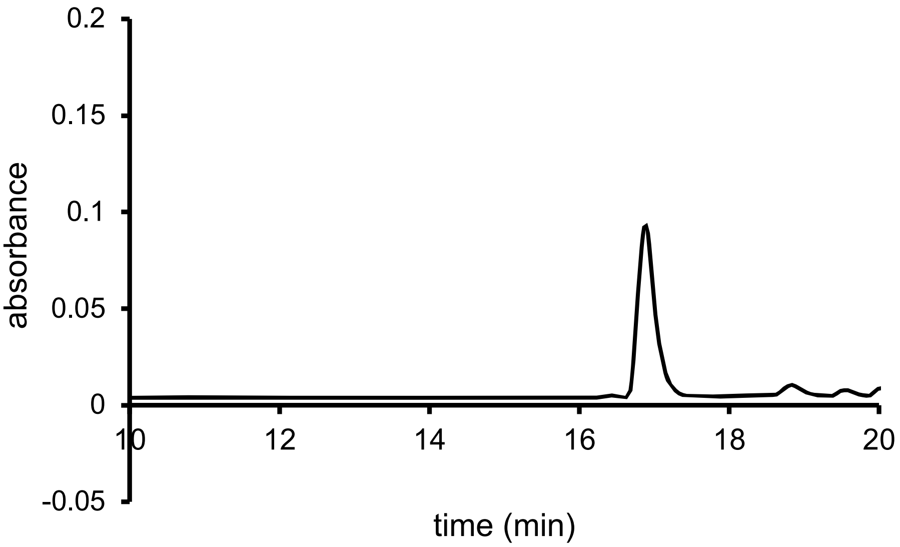


QTOF High Resolution Mass Spectrum for **7b** (m/z 490.2772 for $[M+H]^+$, found 490.2775)

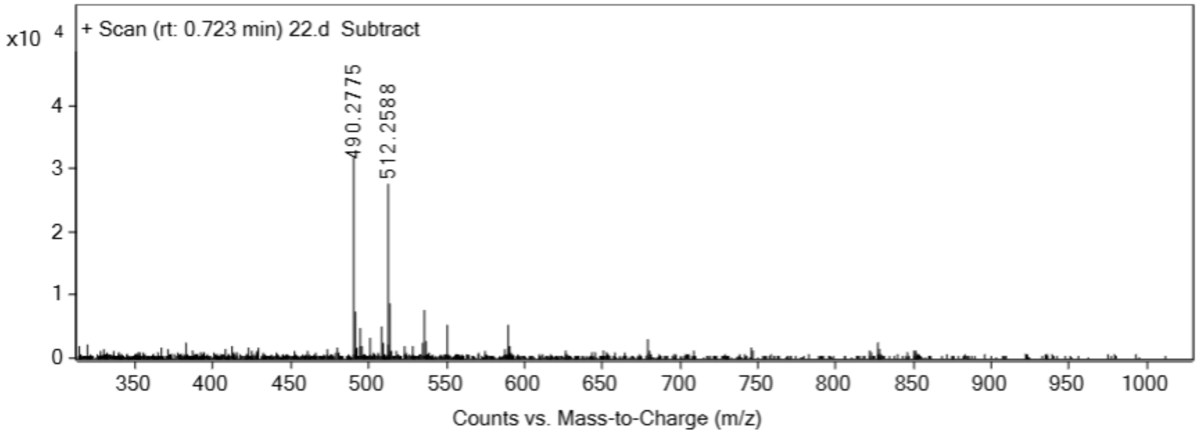


**7c**

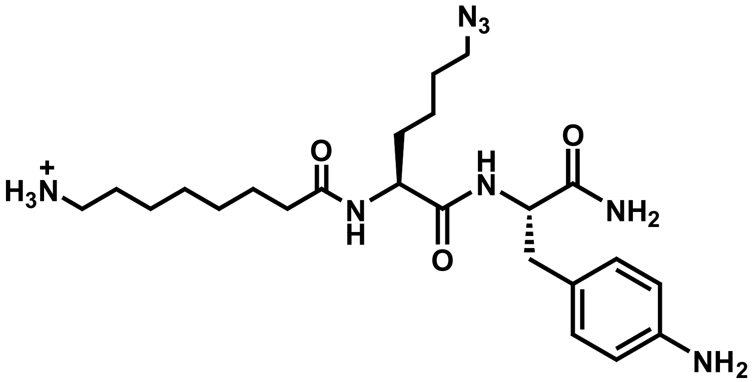


Analytical HPLC Chromatogram of **7c**

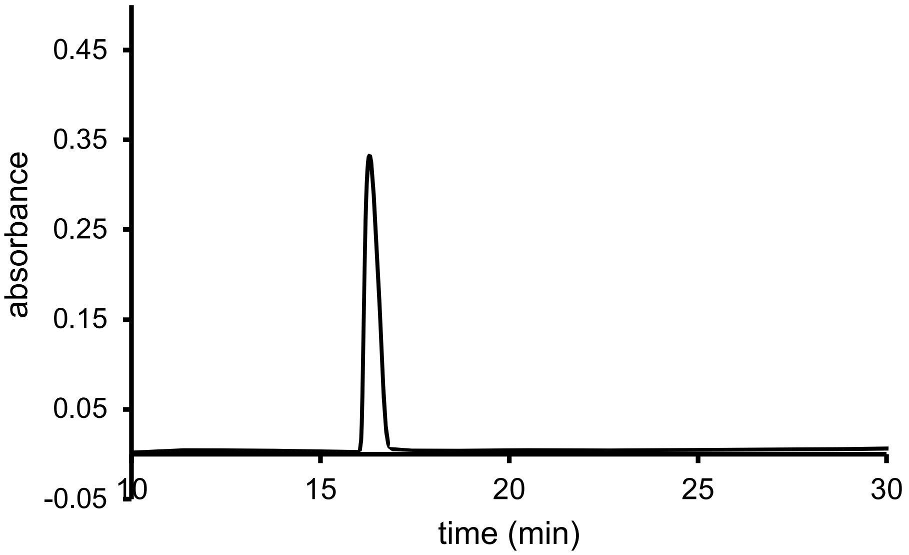


QTOF High Resolution Mass Spectrum for **7c** (m/z 475.3140 for $[M+H]^+$, found 475. 3137)

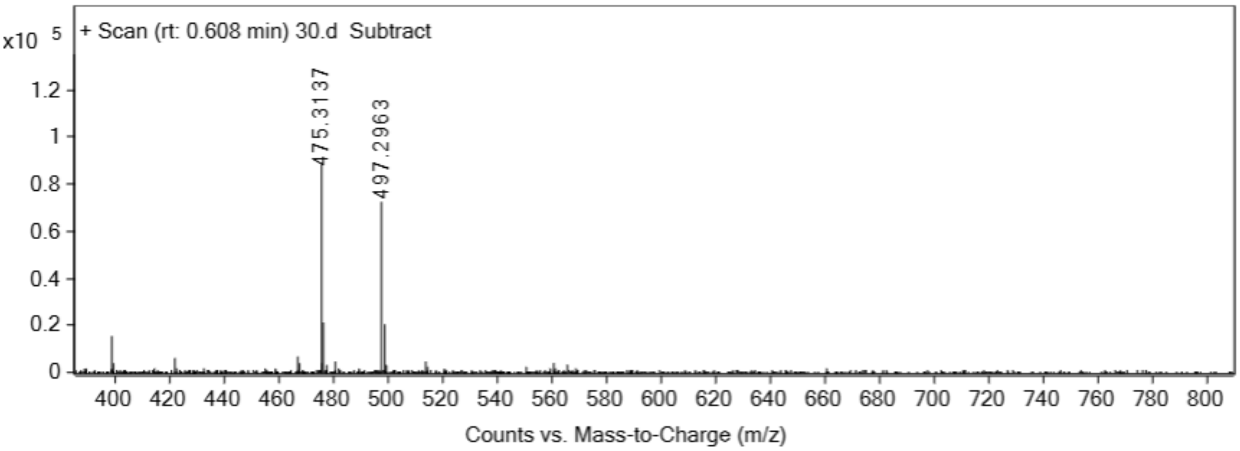


**7d**

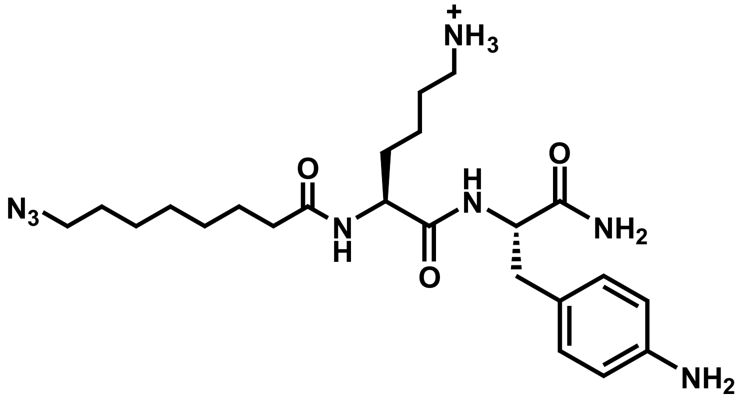

Analytical HPLC Chromatogram of **7d**

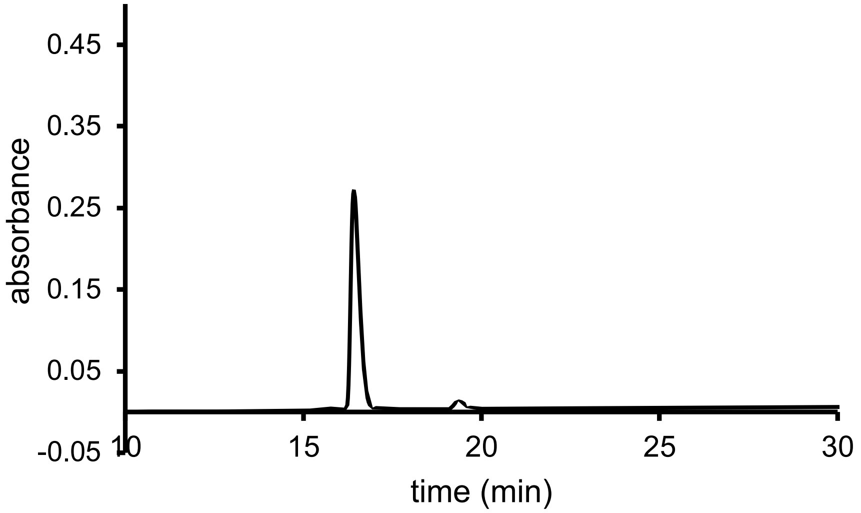


QTOF High Resolution Mass Spectrum for **7c** (m/z 475.3140 for [M+H]$^+$, found 475. 3132)

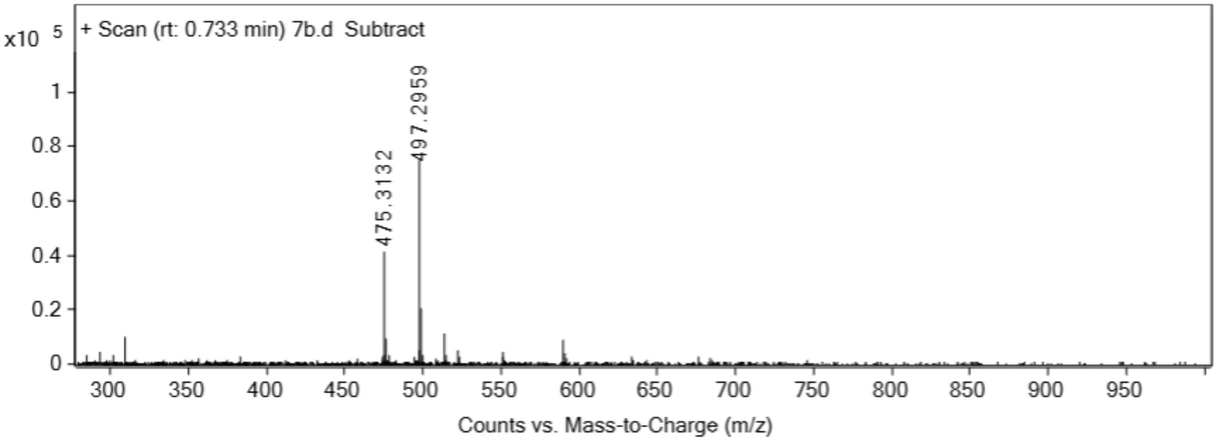


**3b**

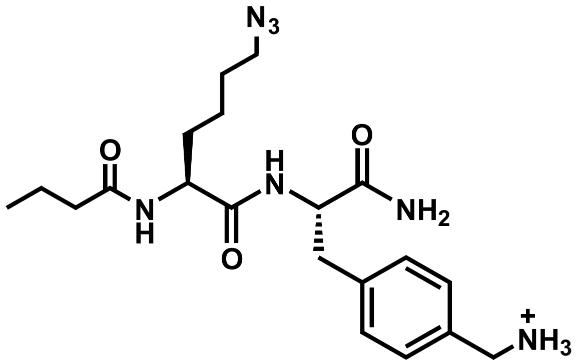


Analytical HPLC Chromatogram of **3b**

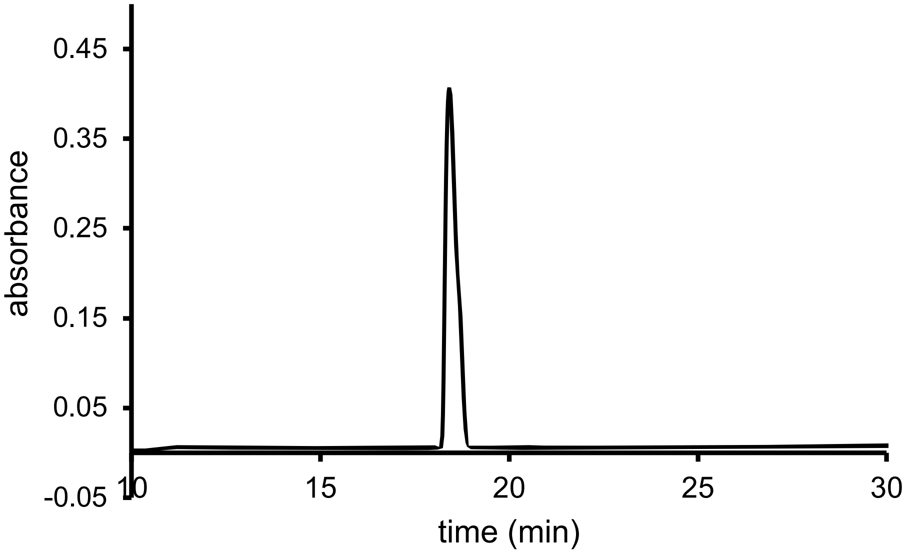


QTOF High Resolution Mass Spectrum for **3b** (m/z 418.2563 for $[M+H]^+$, found 418.2566)

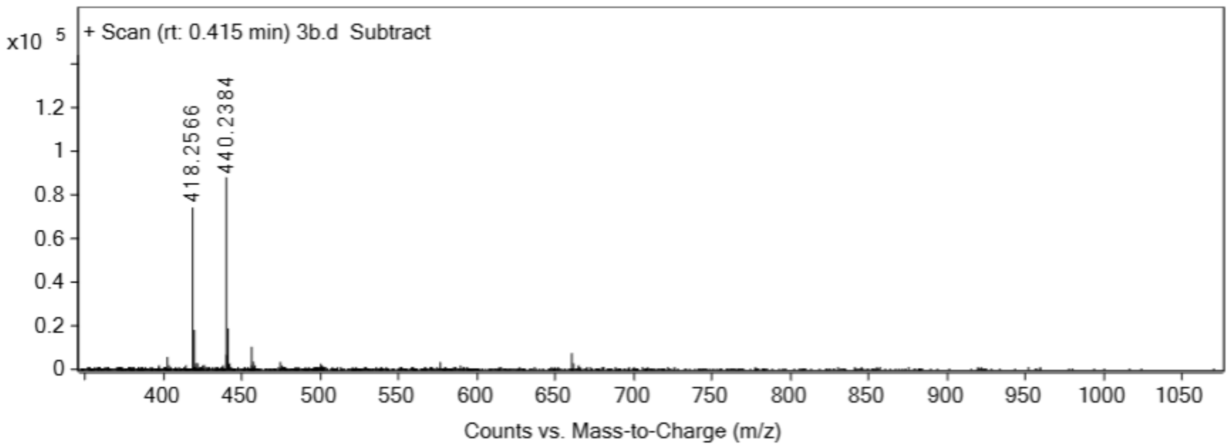


**3c**

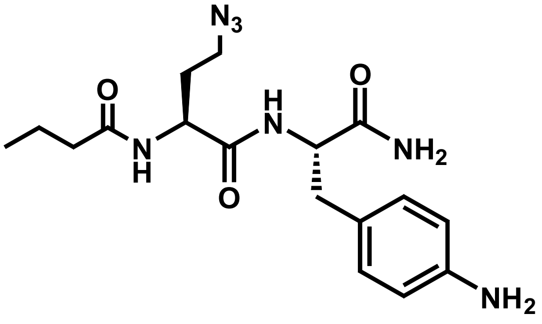


Analytical HPLC Chromatogram of **3c**

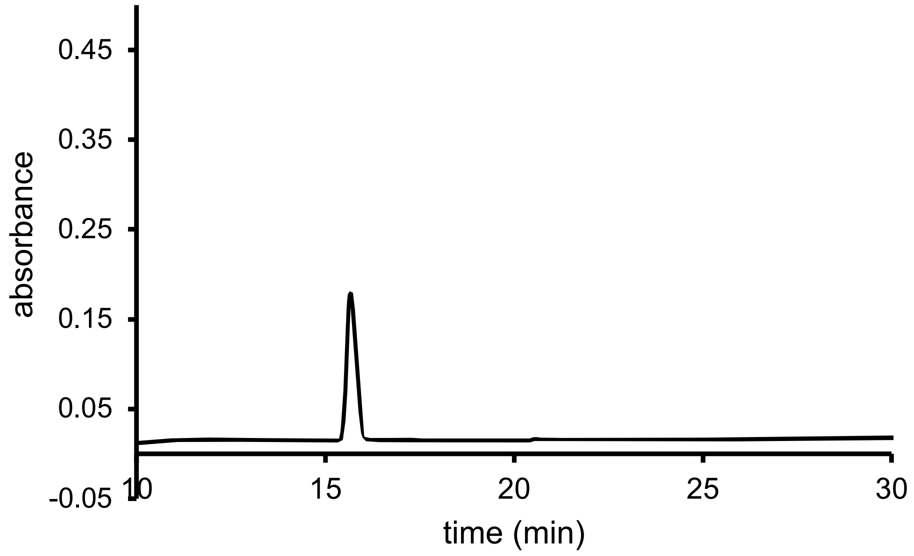


QTOF High Resolution Mass Spectrum for **3c** (m/z 376.2092 for $[M+H]^+$, found 376.2097)

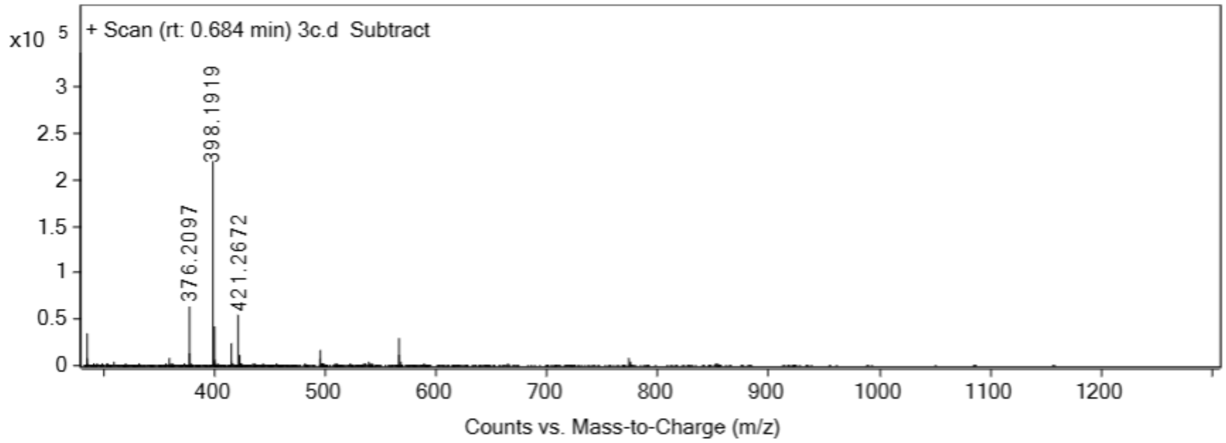


**3d**

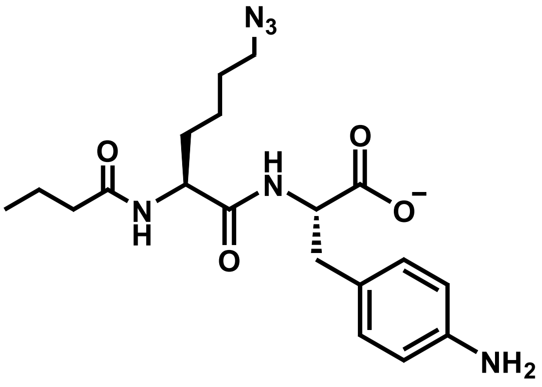


Analytical HPLC Chromatogram of **3d**

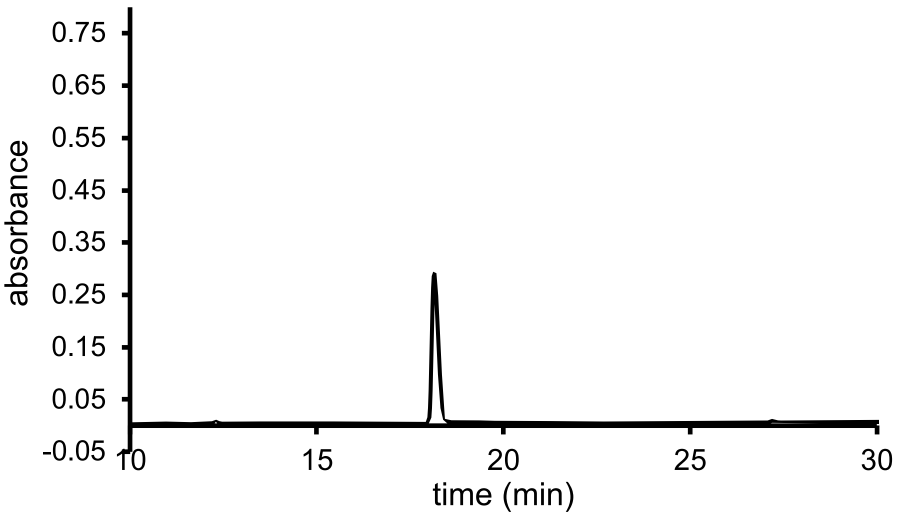


QTOF High Resolution Mass Spectrum for **3d** (m/z 405.2265 for $[M+H]^+$, found 405.2260)

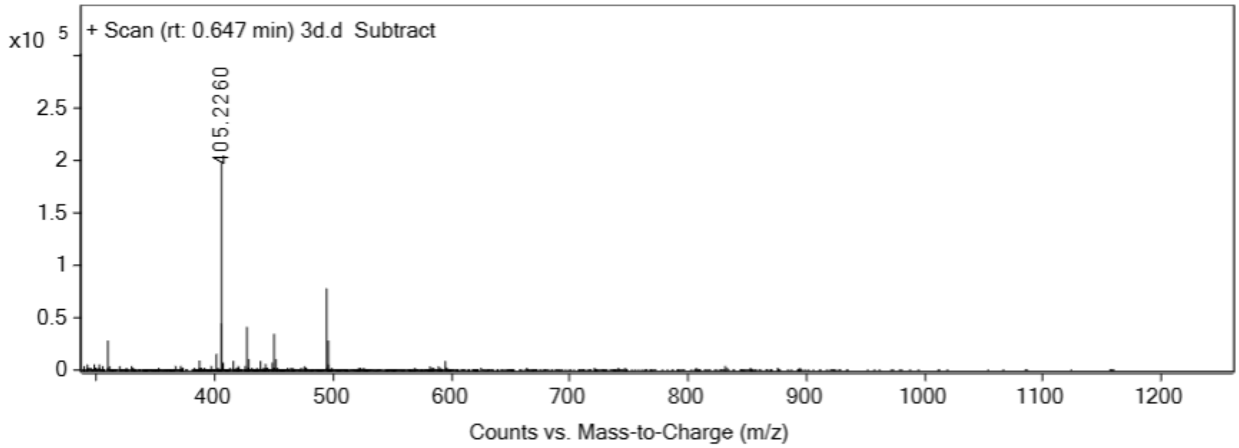


#### Coum0

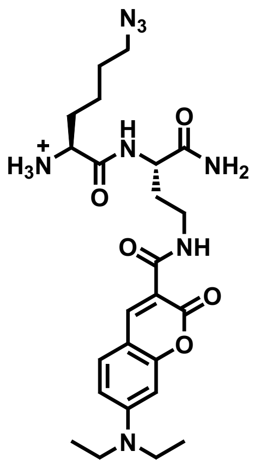


Analytical HPLC Chromatogram of **Coum0**

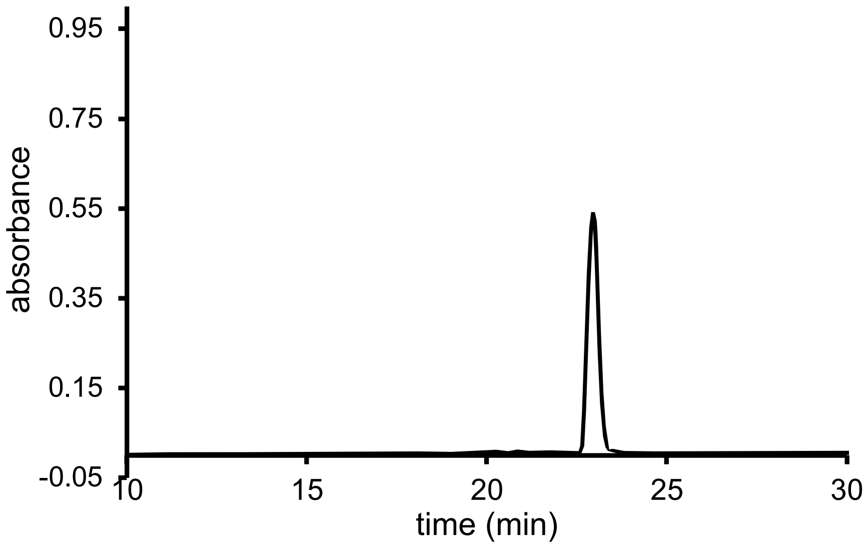


QTOF High Resolution Mass Spectrum for **Coum0** (m/z 515. 2725 for [M+H]$^+$, found 515.2749)

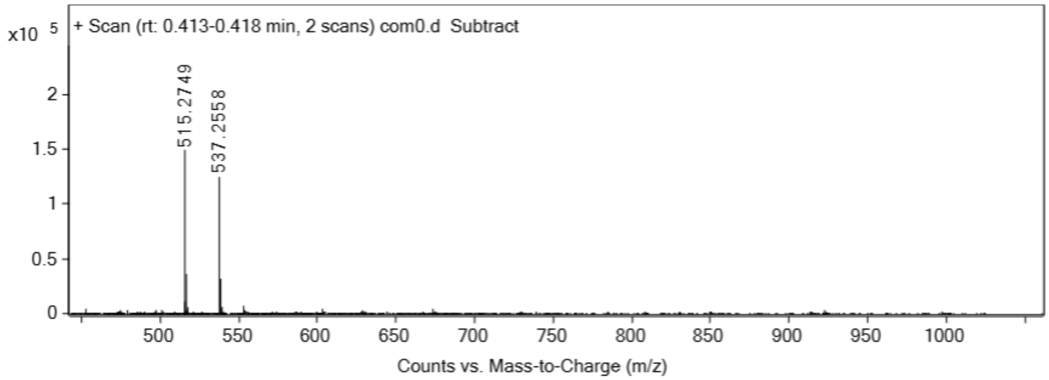


**Coum4**

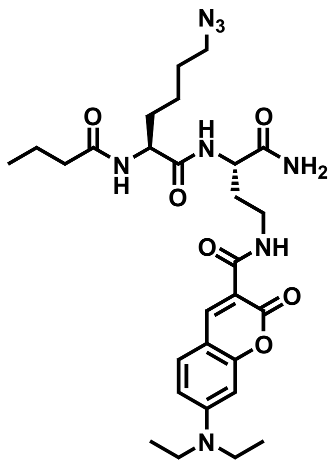


Analytical HPLC Chromatogram of **Coum4**

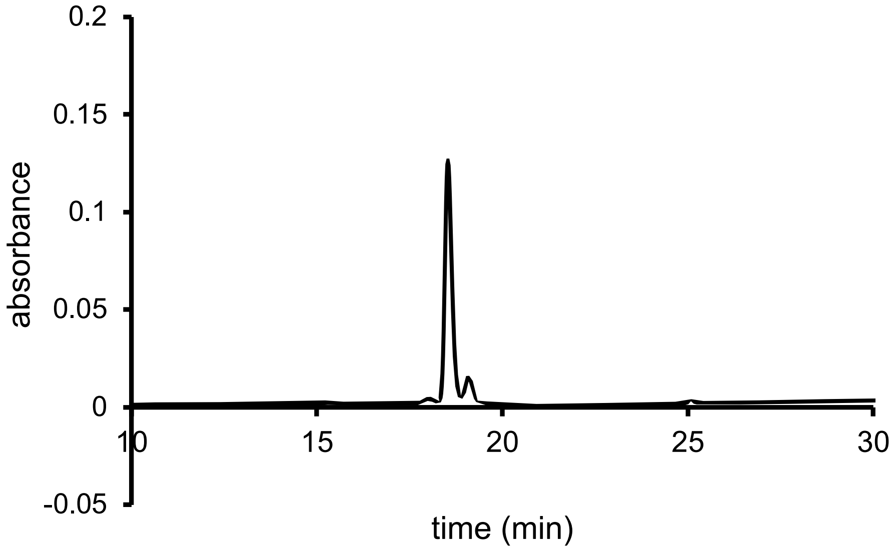


QTOF High Resolution Mass Spectrum for **Coum4** (m/z 585.3144 for $[M+H]^+$, found 585.3156)

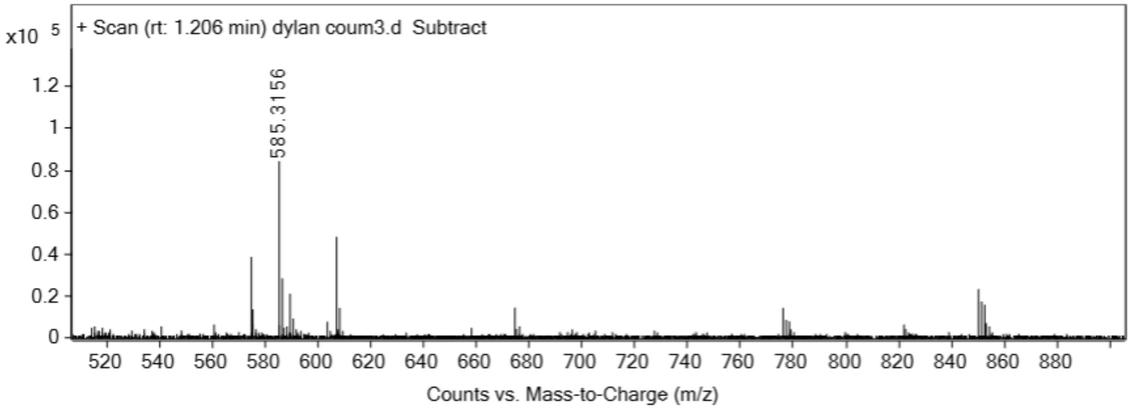


**Coum8**

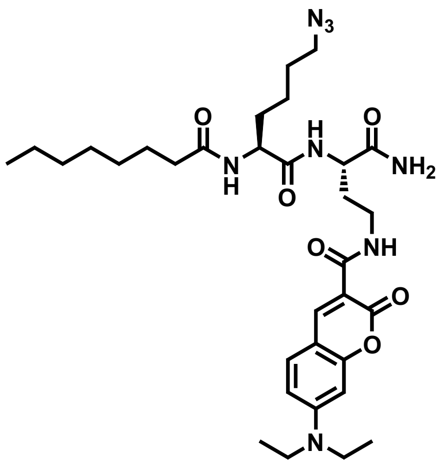


Analytical HPLC Chromatogram of **Coum8**

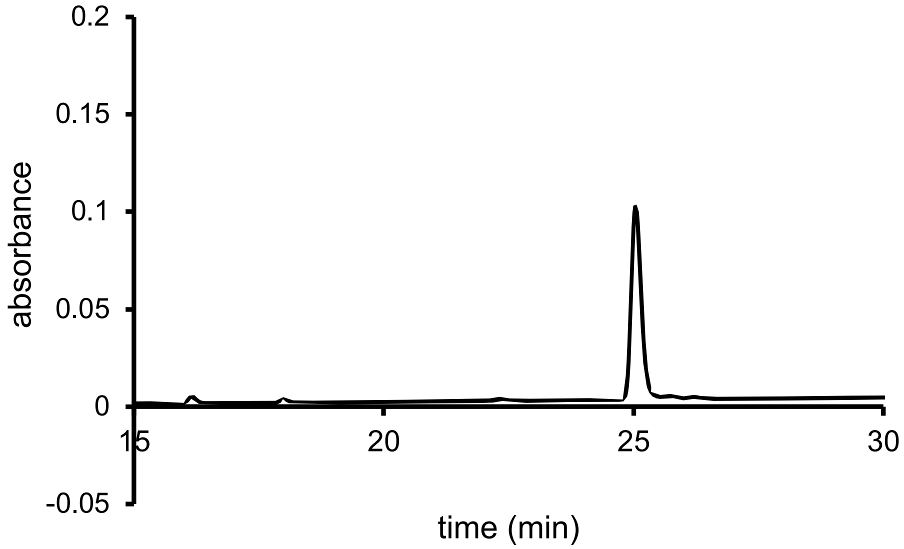


QTOF High Resolution Mass Spectrum for **Coum8** (m/z 641.3770 for $[M+H]^+$, found 641.3776)

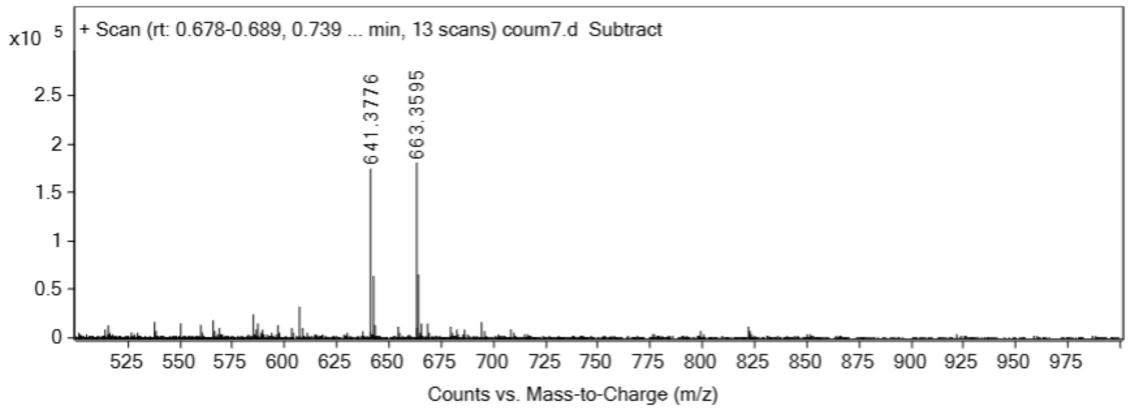


**Coum12**

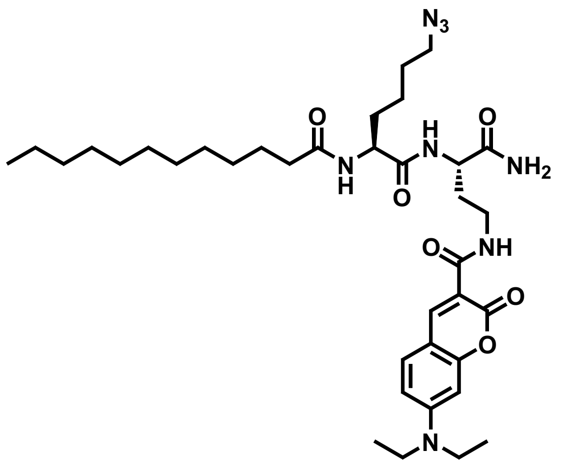


Analytical HPLC Chromatogram of **Coum12**

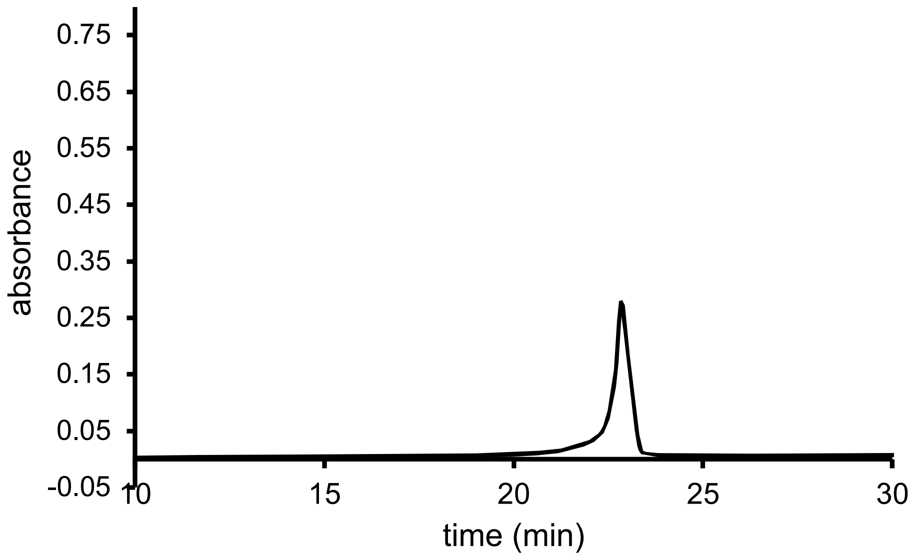


QTOF High Resolution Mass Spectrum for **Coum12** (m/z 697.4396 for $[M+H]^+$, found 697.4425)

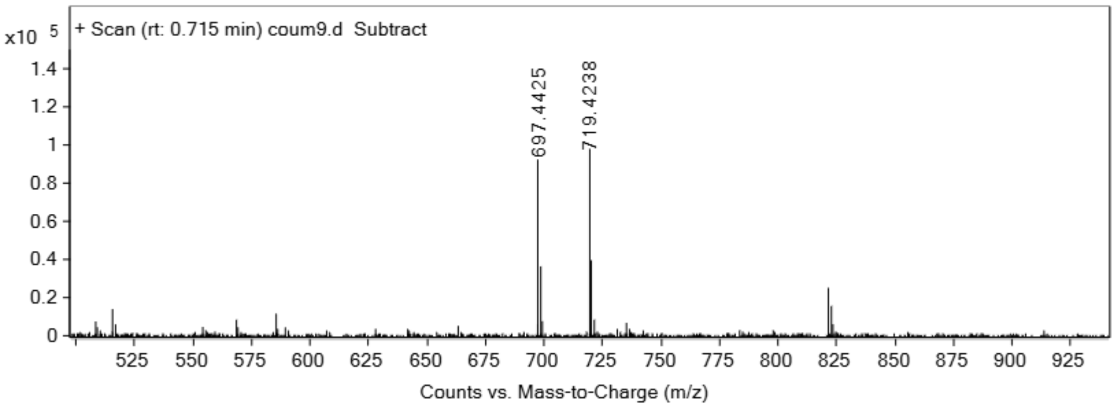

| Compound | $CP_{50}$ *E. coli* (μM) | $CP_{50}$ HeLa (μM) |
|---|---|---|
| **P0** | 13.81 ± 0.85 | >100 |
| **Sat1** | 65.21 ± 1.01 | >250 |

**Table S1.** Calculated $CP_{50}$ values from the curve-fits of **P0** and **Sat1.** Data are represented as mean +/- SD (n = 3) of biological replicates.

| Compound | $CP_{50}$ *E. coli* (μM) | $CP_{50}$ HeLa (μM) |
|---|---|---|
| **Sat1** | 65.21 ± 1.1 | >250 |
| **Sat2** | 35.53 ± 1.6 | >250 |
| **Sat5** | >250 | 34.16 ±.90 |
| **Sat7** | >250 | 8.998 ± 1.4 |
| **Unsat4** | >250 | 39.20 ± 1.1 |
| **Cyc2** | >250 | >250 |
| **Brn2** | >250 | 87.30 ± 1.6 |
| **Het2** | >250 | 20.89 ± 1.1 |
| **Mix3** | >250 | 21.86 ± 0.92 |

**Table S2.** Calculated $CP_{50}$ values from the curve-fits from the chosen sub library of compounds. Data are represented as mean +/- SD (n = 3) of biological replicates.
